# Tubular lysosomes harbor active ion gradients and poise macrophages for phagocytosis

**DOI:** 10.1101/2020.12.05.413229

**Authors:** Bhavyashree Suresh, Anand Saminathan, Kasturi Chakraborty, Chang Cui, Lev Becker, Yamuna Krishnan

**Affiliations:** Department of Chemistry, The University of Chicago, IL, 60637, USA; Grossman Institute of Neuroscience, Quantitative Biology and Human Behavior, The University of Chicago, Chicago, IL 60637, USA; Committee on Cancer Biology, Ben May Department for Cancer Research, The University of Chicago, IL, 60637, USA; Ben May Department for Cancer Research, The University of Chicago, Chicago, IL, USA

## Abstract

Lysosomes adopt dynamic, tubular states that regulate antigen presentation, phagosome resolution and autophagy. To date, tubular lysosomes have been studied either by inducing autophagy or by activating immune cells, both of which lead to cell states where lysosomal gene expression differs from the resting state. Therefore, it has been challenging to pinpoint the specific biochemical properties lysosomes acquire upon tubulation that could drive their functionality. We describe a DNA-based assembly that tubulates lysosomes in macrophages without activating them. Lumenal proteolytic activity maps at single lysosome resolution revealed that tubular lysosomes were less degradative. Further, they showed striking proximal to distal lumenal pH and Ca^2+^ gradients. Such gradients had been predicted, but never previously observed. We now identify a role for tubular lysosomes whereby they poise resting macrophages for phagocytosis. The ability to tubulate lysosomes without having to starve or activate immune cells may help reveal new roles for tubular lysosomes.

## Introduction

The lysosome is a multifunctional organelle, responding to environmental cues as well as cellular and organismal needs(*1, 2*). Its different functions have been linked to differences in lysosome size, shape, location, abundance, and composition(*3, 4*). Although lysosomes are generally vesicular, they can form reticulated tubules that are several microns long(*4–6*). New roles are rapidly emerging for tubular lysosomes(*5, 7–11*). In dendritic cells, tubular lysosomes fuse with the cell membrane leading to antigen presentation(*12–14*). Tubulation also promotes the lysosomal efflux of bacterial peptides into the cytosol to activate NOD-like receptors(*15*). In autophagic lysosome reformation (ALR), lysosomes tubulate and undergo scission, producing proto-lysosomes(*6, 7*). Tubular lysosomes are found even in invertebrates and protozoans revealing that the capacity to tubulate is conserved across phyla. For instance, tubular lysosomes are seen in remodeling muscle cells of *D. melanogaster*, the epidermis of aging *C. elegans*, as well as in *L. mexicana*(*9, 16–18*).

Tubular lysosomes are generally observed when the cell is undergoing either autophagy or immune activation(*4, 7, 9, 11, 16, 17, 19*). While tubulation *per se* occurs universally along microtubules, the biochemistry within tubular lysosomes in different contexts varies remarkably. For instance, autophagy-induced tubular lysosomes in fibroblasts were found to be less acidic and less degradative than vesicular lysosomes(*7*), whereas in muscles of *D. melanogaster*, they were as acidic and as degradative as vesicular ones(*16*). Inhibiting lysosomal acidification collapses tubular lysosomes in *L. mexicana*,^(*18*)^ whereas in muscles of *D. melanogaster* acidification is not critical for tubulation(*9*). In rodents, autophagy-induced tubular lysosomes in kidney cells are hypoacidic unlike those in activated dendritic cells(*7, 20*). Even though autophagy and the immune response activate different transcription programs that upregulate distinct sets of lysosomal proteins, the differences between tubular and vesicular lysosomes across many of the above studies are challenging to reconcile.

Thus, if we could tubulate lysosomes without inducing autophagy or the immune response, we would be able to better understand the nature and functionality of tubular lysosomes independent of cell state. If we could then compare the proteolytic activity and ionic composition of the resultant vesicular and tubular lysosomes it could reveal fundamental functional differences, if any, between both morphologies and perhaps pinpoint key properties acquired by lysosomes upon tubulation. Here we describe our studies arising from our serendipitous discovery of a DNA-based reagent that acutely tubulates lysosomes without activating macrophages. This DNA-based reagent, *Tudor* (<u>Tu</u>bular lysosome <u>D</u>NA rep<u>or</u>ter) binds Ku70/80 heterodimers at the plasma membrane and tubulates lysosomes via a pathway that is distinct from that triggered by lipopolysaccharide (LPS) that invariably activates macrophages(*21*).

*Tudor* enabled us to compare the properties of tubular lysosomes in activated and resting macrophages. Mapping enzyme activity at single lysosome resolution revealed that tubular lysosomes are proteolytically less active than vesicular ones. Given these differences in proteolytic activity, we analyzed lumenal pH and Ca^2+^ levels at single lysosome resolution, since lysosomal ionic content regulate proteolysis(*22–24*). Although the overall levels of pH and Ca^2+^ in vesicular and tubular lysosomes were similar, tubular lysosomes showed lumenal pH and Ca^2+^ gradients along the length of the tubule. Such gradients in tubular lysosomes had been previously predicted by others and we now provide experimental evidence that supports this prediction. What the model had not predicted, but our observations reveal, is that there are different classes of tubular lysosomes.

Tubular lysosomes are known to play important roles in the late stages of phagocytosis(*25, 26*). They enhance antigen presentation and facilitate phagolysosome resolution(*25–27*). Using *Tudor* to study the intrinsic properties of tubular lysosomes in resting macrophages, we found that tubular lysosomes poise macrophages for phagocytosis by promoting phagosome formation as well as phagosome-lysosome fusion. Our studies now reveal a role for tubular lysosomes even during the early stages of phagocytosis.

## Results and Discussion

### A DNA nanodevice, *Tudor*, tubulates lysosomes

We describe the design, endocytic uptake pathway and activity of a DNA nanodevice that acutely triggers tubulation of lysosomes in macrophages without activation. This DNA nanodevice, denoted *Tudor* (<u>Tu</u>bular lysosome <u>D</u>NA <u>r</u>eporter), is designed such that two complementary DNA strands A1 and A2 display an actuator domain and a fluorescent reporter respectively (Fig. 1a, SI Table 1). The actuator domain in A1 is a 43-base long DNA aptamer called SA43, known to specifically bind the Ku70/80 dimer at the plasma membrane with a Kd of 21 nM(*28*). Strand A1 is hybridized to A2, which displays the fluorescent dye Alexa 647, to form a 24-base pair (bp) duplex (dsDNA). The fluorophore on *Tudor* enables one to simultaneously evaluate tubulation, uptake efficiency and the sub-cellular distribution of *Tudor* (Fig. 1a). The formation, integrity and purity of *Tudor* was confirmed by gel electrophoresis (Fig. S1).

**Figure 1:**
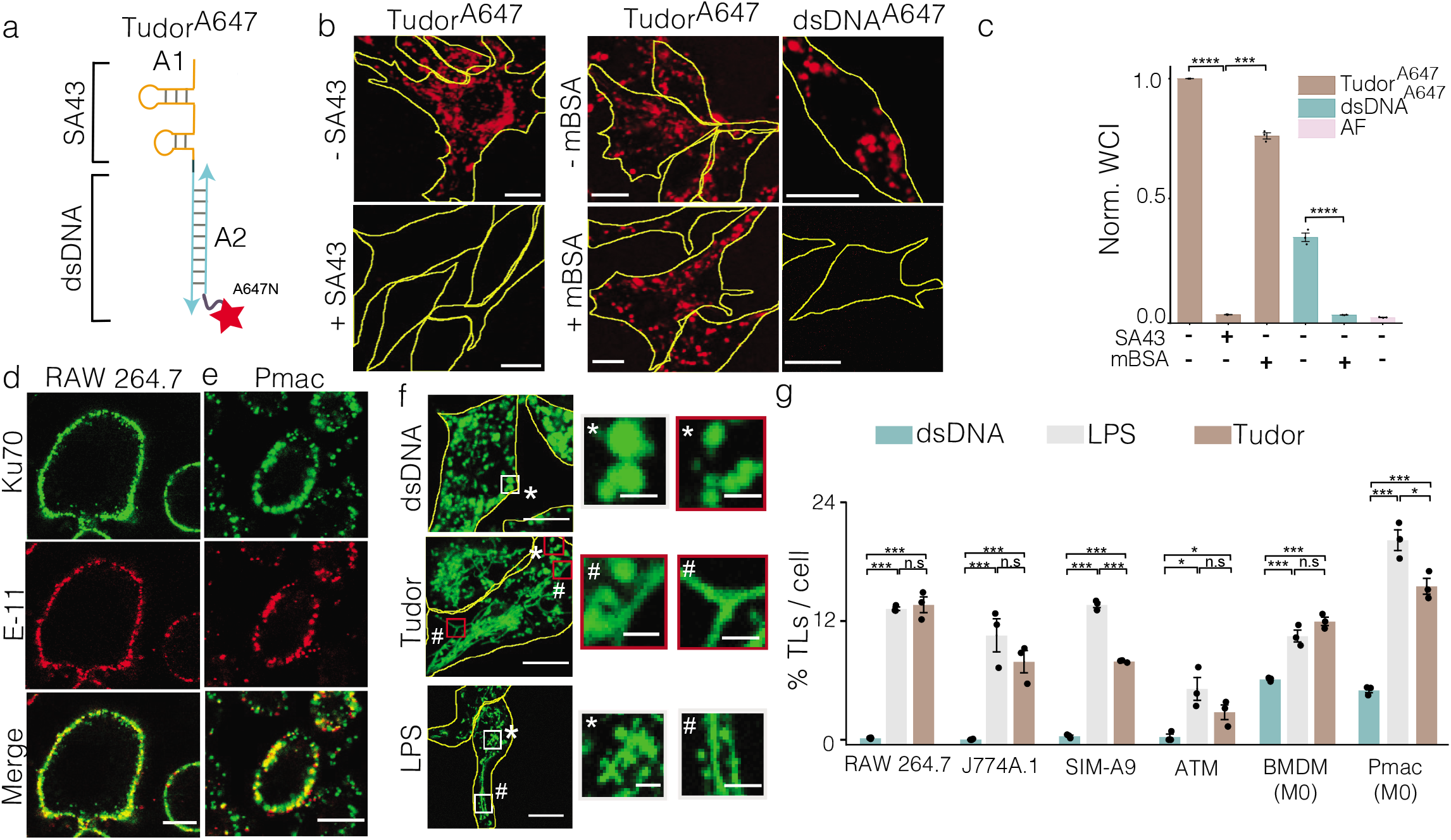
A DNA nanodevice, *Tudor*, tubulates lysosomes. (**a**) Schematic of *Tudor* containing modules A1, SA43 aptamer to Ku70/80 (actuator domain), hybridized to A2 bearing Alexa 647N (red star). (**b**) Representative confocal images of *Tudor^A647^* and dsDNA^A647^ uptake in presence or absence of indicated competitors (3 μM SA43 or 60 equivalents mBSA) in RAW 264.7 (**c**) Normalized whole cell intensities (WCI) from (b) (n=100 cells). (**d - e**) Representative images of Ku70 (green) and Pan Cadherin (E-11, red) in RAW 264.7 cells (d) and Pmac (e). (**f**) Representative images of TMR dextran labeled RAW 264.7 cells in the presence of dsDNA, *Tudor* and LPS. Magnified images of white boxed regions [top panel], vesicular lysosomes (VL,*) and tubular lysosomes (TL,#) red boxed regions [top and middle panel] and LPS treated [bottom panel]. (**g**) Quantification of % TLs per cell in indicated cell types treated with dsDNA, *Tudor* and LPS (*n* = 50 cells). n.s: non-significant; \*\*\*\**P*< 0.00005; ***P< 0.0005; \**P*< 0.05; (one-way ANOVA with Tukey *post hoc* test). Imaging experiments were performed in triplicate with similar results. Error bars represent standard error of mean (s.e.m) from three independent experiments. Scale bar, 10 μm; inset scale bars, 2 μm.

We found that *Tudor* was internalized by receptor mediated endocytosis in RAW 264.7 cells. However, *Tudor* uptake was not dependent on scavenger receptors that are most commonly involved in dsDNA uptake(*29*) but rather occurred via Ku70/80. We first tested the involvement of scavenger receptors by trying to compete out *Tudor* uptake with excess maleylated BSA (mBSA), a ligand for scavenger receptors(*30*). Unlike dsDNA which is uptaken by scavenger receptors and gets competed out effectively by mBSA, *Tudor* uptake was minimally affected indicating that uptake was not via scavenger receptors (Fig. 1b-c). However, uptake was completely abolished in the presence of 60 eq. of unlabeled SA43, revealing that *Tudor* internalization was mediated by specific interactions with Ku70/80 at the cell surface (Fig. 1b-c). In addition to its ubiquitous presence in the nucleus, the cell-surface abundance of the Ku70/80 heterodimer is documented in many cell types including macrophages(*31–37*). Indeed, immunostaining without permeabilizing the plasma membrane, revealed the presence of Ku70 at the plasma membrane of macrophage cell lines, such as RAW 264.7, J774A.1, SIM-A9.1. This also held true in all primary macrophages with different activation states tested, such as naïve murine bone marrow-derived macrophages (BMDM), peritoneal macrophages (Pmac), and their polarized states including LPS/INFg (pro-inflammatory) or the M2 phenotype with IL4 (anti-inflammatory) (Fig. 1d, e and Fig. S2).

Upon treating macrophages with 100 nM *Tudor* for 4h, we found that it labeled organelles with vesicular as well as highly tubular morphologies. When lysosomes in macrophages were pre-labeled with TMR dextran, they colocalized with internalized *Tudor* revealing that the vesicular and tubular *Tudor*-containing compartments were, in fact, lysosomes (Fig. 1f and S3). Tubulated lysosomes that were formed by *Tudor* treatment phenocopied those observed by lipopolysaccharide (LPS, 100 ng/mL) treatment (Fig. 1f-g). LPS is a canonical reagent that both activates macrophages via TLR4 and simultaneously tubulates lysosomes by a well-defined pathway(*5, 21, 38, 39*). Tubulation was not observed when RAW 264.7 cells were either treated with a *Tudor* variant lacking the actuator domain, denoted as dsDNA, or a single stranded DNA aptamer against a different cell surface protein, MUC-1(*40*) (Fig. 1f-g). Treating RAW 264.7 cells with CpG DNA which activates macrophages via TLR9 also failed to induce tubulation, suggesting that *Tudor*-induced tubulation was specifically triggered by SA43, and was not due to generic effects triggered by DNA (Fig. S4). Further, *Tudor* efficiently tubulated lysosomes in other macrophage cell lines such as J774A.1, SIM-A9, as well as primary macrophages such as BMDM, Pmacs and adipose tissue macrophages (ATM) (Fig. 1g and Fig. S5-S7).

### *Tudor* tubulates lysosomes in macrophages without immune activation

Since LPS not only tubulates lysosomes but also induces immune activation, we sought to test whether tubulating lysosomes with *Tudor* stimulates the immune response in macrophages. Tubulation efficiency is parametrized as the percentage area of tubular lysosomes 4h after treatment with *Tudor* or LPS. We first confirmed that tubulation induced by either ssDNA or dsDNA was negligible (Fig. 2a-c). Since lysosome tubulation by *Tudor* appeared to be unrelated to immunostimulation by the DNA scaffold, we tested whether *Tudor* treatment of resting macrophages (M0) polarized them into M1 or M2 states. Interestingly, mRNA expression data revealed no significant upregulation of M1 or M2 markers implying that *Tudor* did not polarize primary M0 macrophages towards either state in BMDM or Pmac (Fig. 2e and Fig. S8). Nevertheless, the kinetics and extent of *Tudor*–induced tubulation phenocopied that induced by LPS, suggesting that both agents could potentially act via a similar pathway (Fig. 2d).

**Figure 2:**
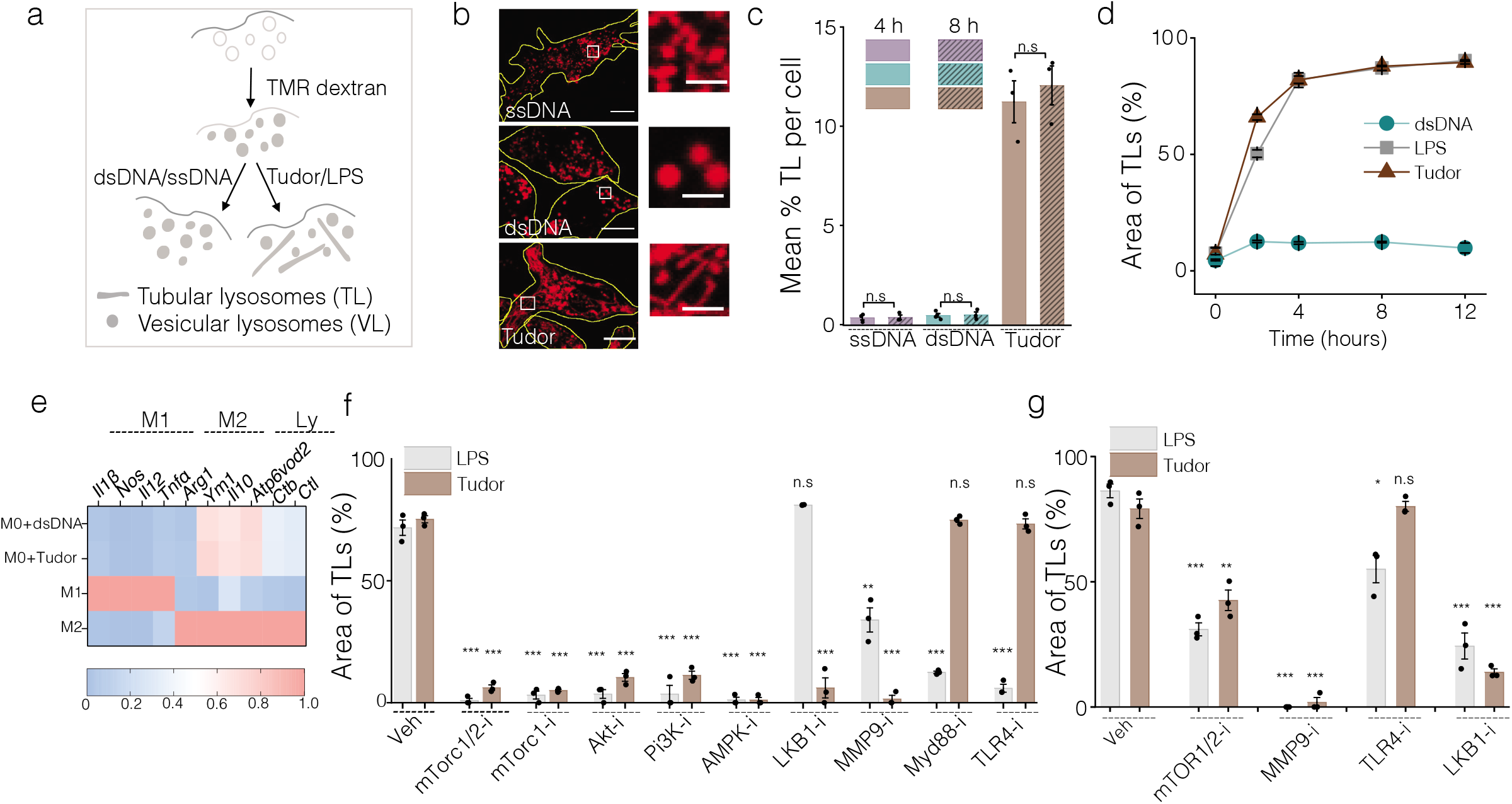
*Tudor* non-immunogenically tubulates lysosomes in macrophages. (**a**) Schematic of lysosomal tubulation assay (**b**) Representative images of TMR-dextran labeled lysosomes of RAW 264.7 upon treatment with ssDNA, dsDNA and *Tudor*. Inset: Magnified images of VLs and TLs. (**c**) Quantification of (b) as % TLs per cell at 4 and 8 hours of treatment (n = 25 cells). (**d**) % area of TLs w.r.t. time upon addition of *Tudor*, LPS or dsDNA in RAW 264.7 cells (n= 20 cells). (**e**) Heat maps representing fold change of mRNA levels of M1, M2 and lysosomal markers (Ly) in BMDM (M0) upon treatment with dsDNA and *Tudor*. Fold change of M1 markers are normalized to M1 BMDM, fold change of M2 markers are normalized to M2 BMDM, and fold change of lysosomal markers are normalized to M0+dsNDA. (**f** and **g**) Quantification of % area of TLs in the presence *Tudor* or LPS and pharmacological inhibitors for indicated protein (*protein-i)* in RAW 264.7 cells, (n=20 cells) (f) and in M0 BMDMs (n = 20 cells) (g). n.s: non-significant; \*\*\**P*< 0.0005; **P< 0.005; *P< 0.05 (one-way ANOVA with Tukey *post hoc* test). Error bars represent standard error of mean (s.e.m) for all experiments shown here. Data represented from at least 3 independent experiments. Scale bar: 10 μm and inset scale bar: 4 μm.

To identify the players responsible for *Tudor*-induced tubulation, we quantified lysosomal tubulation in RAW 264.7 cells in the presence of different pharmacological inhibitors. In the presence of TAK242, a potent TLR4 inhibitor, or the Myd88 inhibitory peptide, LPS failed to tubulate lysosomes, consistent with previous findings(*5*). In contrast, *Tudor*-induced tubulation was independent of TLR4 as well as Myd88 (Fig. 2f and Fig. S9a). To test the involvement of other TLRs, we pharmacologically inhibited TLR5, TLR3, TLR2/TLR6, TLR1/TLR2 prior to treatment with *Tudor*. We observed no significant effects, revealing that *Tudor* did not tubulate lysosomes via TLR stimulation (Fig. S9b, d).

Given that Ku80 in the Ku heterodimer interacts with the hemopexin domain of MMP9 to activate the latter at the plasma membrane, we tested the role of MMP9 in lysosome tubulation(*41, 42*). An MMP9 protease activity assay revealed that *Tudor* treatment activated MMP9 (Fig. S9e). Further, inhibiting MMP9 with MMP9 inhibitor-1(*43*) abolished both *Tudor* and LPS-induced lysosome tubulation (Fig. 2f and Fig. S9a). These findings were recapitulated in primary macrophages such as BMDMs (Fig. 2g), demonstrating that MMP9 was indispensable for both LPS and *Tudor*-mediated lysosome tubulation (Fig. 2g and Fig. S10).

To test the hypothesis that LPS and *Tudor* might work via common players late in the tubulation pathway, we pharmacologically inhibited mTOR, Akt or PI3K with Torin-1(*5, 44*), Akt-I(*5*) or ZSTK474(*45*) respectively and treated them with either LPS or *Tudor*. Indeed, both LPS and *Tudor*-mediated tubulation depended on the PI3K-Akt-mTOR cascade (Fig. 2f and Fig. S9a). SiRNA knockdown of *Arl8-b* further confirmed that *Tudor*-induced tubulation occurred via the same players as the LPS-induced pathway, downstream of PI3K(*5, 38*) (Fig. S11).

There are many ways to activate PI3K. For instance, cSrc activation of PI3K is mediated by c-Cbl(*46*),(*36*). Alternatively, TLR2 activation in innate immune cells activates RAC1 which in turn can activate Akt(*47*). JAK1/2 can also activate PI3K(*48*). Treatment with potent inhibitors for cSrc (Dasatinib), RAC1 (RAC1i) and JAK1/2 (barcitinib) revealed that neither *Tudor* nor LPS acted via these pro-inflammatory signaling pathways (Fig. S9c-d). We therefore targeted anti-inflammatory pathways that involved PI3K and Akt(*49*). In our hands, inhibiting AMPK with Dorsomorphin (compound C)(*50*), impeded tubulation mediated by both LPS and *Tudor* in RAW 264.7 macrophages, even though prior work showed that AMPK activation prevents LPS-mediated tubulation(*5*). AMPK activity is regulated by kinases such as LKB1 or TGFβ activating kinase 1 (TAK1)(*51*). We found that *Tudor*–induced tubulation was dependent on LKB1, but not on TAK1 (Fig. 2f and Fig. S9a, c-d). Our studies reveal two new players in the lysosome tubulation pathway, LKB1 and AMPK, that negatively regulate mTOR, the significance of which we discuss later.

### Single lysosome protease activity maps reveal that tubular lysosomes are less degradative

Tubular lysosomes facilitate cellular functions ranging from antigen presentation to autophagy, yet it is still not clear whether tubulation changes the lumenal biochemistry of the lysosome. Lysosomes house more than 50 different hydrolases that cumulatively degrade endocytosed cargo(*52*). When macrophages are stimulated with LPS lysosome tubulation is induced and many lysosomal enzymes upregulated(*53, 54*). We therefore quantitatively mapped enzymatic activity at the resolution of single lysosomes in live RAW 264.7 macrophages treated with either LPS or *Tudor*. We allowed cells with Alexa488-dextran-labeled lysosomes to endocytose DQ™-BSA and compared the proteolytic activity within tubular and vesicular lysosomes (Fig. 3a-b and Fig. S12). We found that regardless of how lysosomes were tubulated, increases in proteolysis was confined to vesicular lysosomes while the activity in tubular lysosomes was unaffected (Fig. 3b).

**Figure 3:**
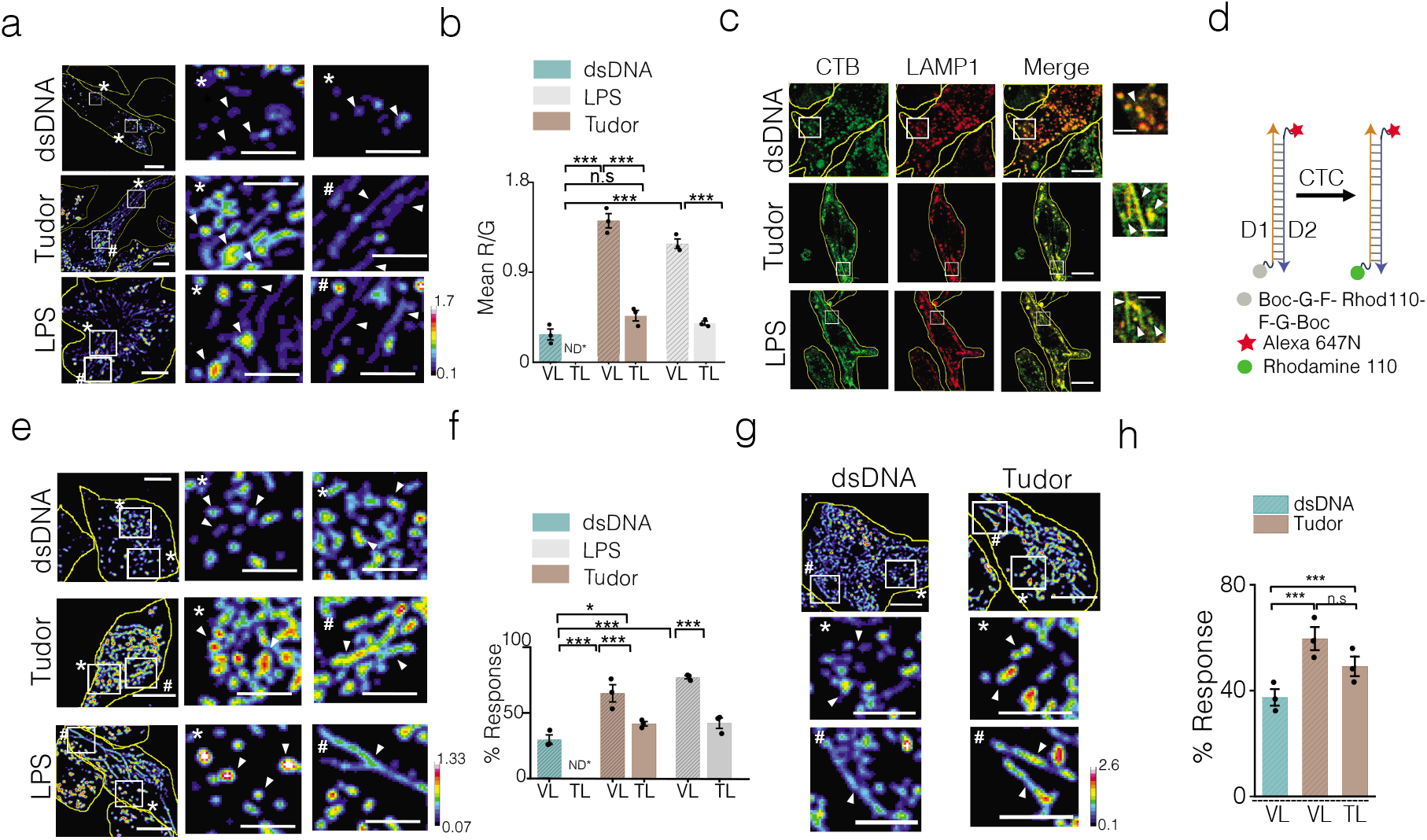
Single lysosome enzymatic cleavage maps show lower degradation in tubular lysosomes. (**a**) Representative pseudocolour R/G images of Alexa488 dextran (G) and DQ™ Red BSA (R) labeled lysosomes in dsDNA; *Tudor* and LPS treated RAW 264.7 (left). Magnified images shown in white box (right). (**b**) Quantification of (a) as mean R/G ratio at single lysosome resolution (n = 50 cells; m = 200 lysosomes). (**c**) Representative confocal images of RAW 264.7 immunostained for CTB (green) and LAMP1 (red) upon treatment with dsDNA, *Tudor* and LPS. Magnified images are shown in white box (right). (**d**) Schematic of DNA based CTC activity reporter consisting of DNA duplex (orange and blue ladder) with sensing module (caged Rhodamine 110, grey) and normalizing module (Alexa 647N, red) and cleaved or always on module (Rhodamine 110, green). (**e**) Representative pseudocolored G/R images of CTC activity reporter labeled RAW 264.7 pretreated with dsDNA; *Tudor* or LPS (left). Magnified images of the white boxed region (right). (**f**) Quantification of (e) as % response of CTC reporter in VLs and TLs in dsDNA, *Tudor* and LPS treated cells (n =50 cells, m=500 lysosomes). (**g**) Representative pseudocolored G/R images of CTC activity reporter labeled BMDMs (M0) pretreated with dsDNA or Tudor (top). Magnified images of the white boxed region (below). (**h**) Quantification of (g) as % response in VLs and TLs of dsDNA and *Tudor* treated cells (n =50, m=500 lysosomes), *P< 0.05; ***P< 0.0005 (one-way ANOVA with Tukey *post hoc* test); n.s: non-significant. ND: not defined. White arrowheads show VLs (*) and TLs (#). Error bars represents standard error of mean (s.e.m) from three independent experiments. Scale bars: 10 μm; inset scale bars: 4 μm.

While DQ™-BSA reveals the overall proteolytic activity in lysosomes, DNA-based enzyme activity reporters can selectively address the contribution of a specific lysosomal protease(*55*). Cathepsin C (CTC) is an aminopeptidase that plays key roles in inflammation, IL1β production, TNF-α production, and macrophage reprogramming(*56–58*). We therefore used a previously published ratiometric DNA-based lysosomal CTC reporter system to probe CTC activity in vesicular and tubular lysosomes in RAW 264.7 cells and BMDM (Fig. 3d-h, Fig. S14-S15) (*55*). The CTC activity reporter is a DNA duplex that localizes in the lysosome via scavenger receptor mediated endocytosis. It bears a reference fluorophore and a Rhodamine110 dye whose fluorescence is caged by phenylglycyl (FG) groups that are a substrate for CTC. CTC activity cleaves the FG dipeptides uncaging fluorescence only in the lysosomes. We found that upon tubulating lysosomes, proteolytic activity in the residual vesicular lysosomes was selectively increased without changing the activity in tubular lysosomes (Fig. 3e-f and Fig. S14a-c).

To test whether the differences in lysosomal activity between tubular and vesicular states arose potentially due to selective partitioning of hydrolases between both types of lysosomes(*7*), we probed the relative abundance of Cathepsin B (CTB) by immunofluorescence. Since tubular lysosomes tend to fragment into vesicular ones with traditional fixation methods, we developed a method that preserves lysosomes in their tubular forms (Fig. S13a-b). We observed no significant differences in the relative abundance of CTB normalized to LAMP1 between tubular and vesicular lysosomes regardless of whether cells were treated with dsDNA, LPS or Tudor (Fig. 3c and Fig. S13c).

### Lumenal pH and Ca^2+^ maps reveal two major kinds of tubular lysosomes

The differential enzymatic activity within tubular and vesicular lysosomes despite their comparable cathepsin content led us to test for potential differences in their lumenal pH. Additionally, we considered mapping Ca^2+^ since tubular lysosomes undergo active fission and fusion. Lysosomal Ca^2+^ channels such as P2X4 and TPC2 are implicated in fusion(*3, 59–61*), while TRPML1 regulates lysosomal fission(*3*). Moreover, these Ca^2+^ channels strongly depend on mTOR-activity, which is part of the tubulation cascade(*62–64*). We therefore reasoned that lumenal pH and Ca^2+^ maps at single lysosome resolution could provide insight on the formation or function of tubular lysosomes.

In order to map lumenal pH and Ca^2+^ levels in tubular lysosomes in macrophages, we used a DNA-based, pH-correctable Ca^2+^ sensor, *CalipHluor*2.0 (Fig. S18a). It consists of (i) a pH sensing dye, DCF, that is sensitive between pH 4.0 – 6.0 (ii) a Ca^2+^ sensing dye, namely Rhod-5F and (iii) a reference dye, Atto647N, for quantitative ratiometry. We measured the stability of *CalipHluor*2.0 and found that it was ~95% intact at t = 2h within lysosomes of RAW 264.7 cells (Fig. S16). We then mapped lumenal Ca^2+^ and pH in tubular and vesicular lysosomes as follows. We induced tubulation with *Tudor*, then labeled lysosomes in RAW 264.7 cells with *CalipHluor*2.0 and imaged cells in three channels, G, O and R (Fig. 4a and Fig. S17, S18d-S19a). According to the lysosomal pH and Ca^2+^ maps, the overall lumenal pH and Ca^2+^ levels in n~100 individual, tubular lysosomes was comparable to that of ~300 vesicular lysosomes (Fig. 4c-d).

**Figure 4:**
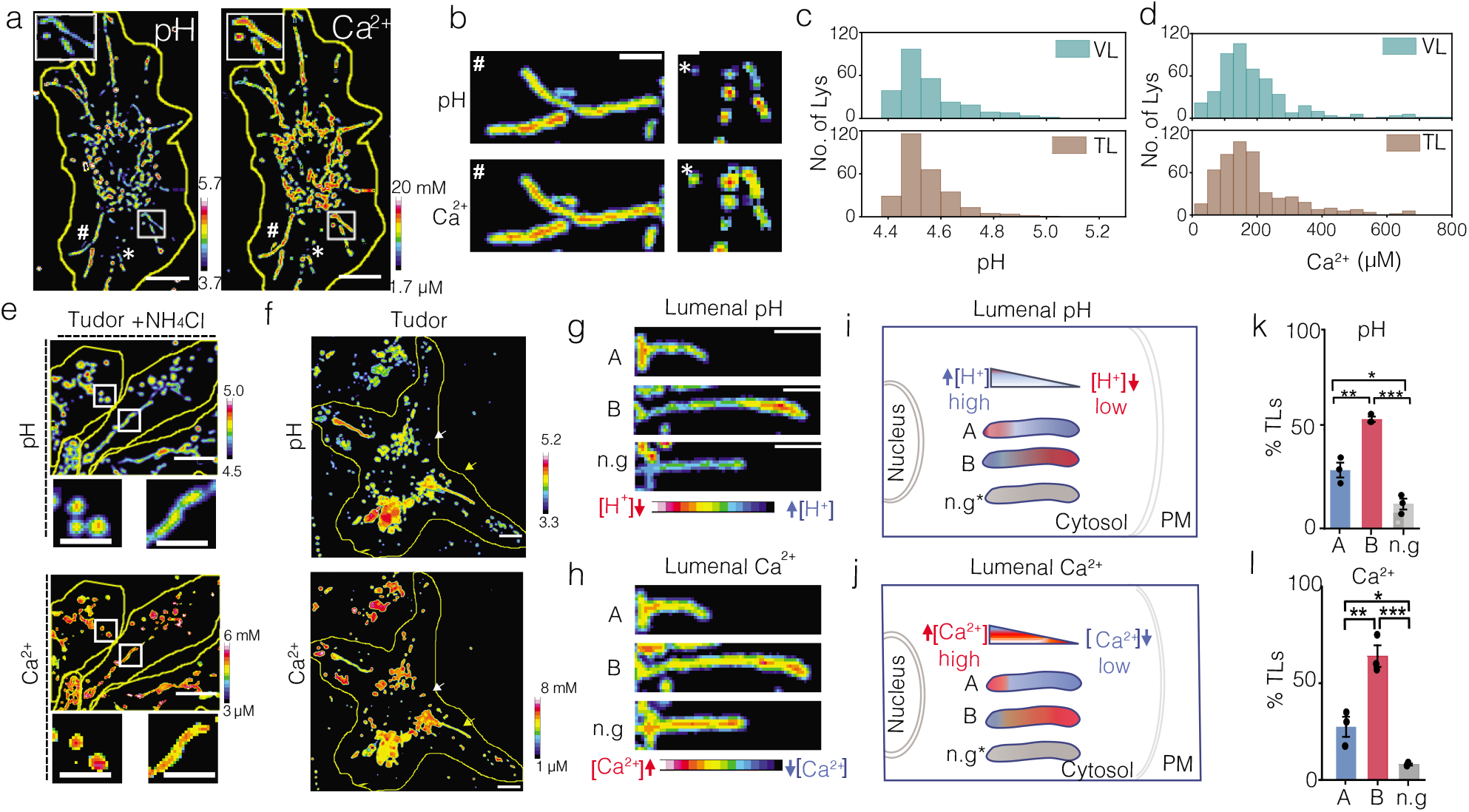
Heterogeneity of ionic gradient within tubular lysosomes. (**a**) Representative pH and Ca^2+^ images of *CalipHluor 2.0* labeled RAW 264.7 cells pretreated with *Tudor*. (**b**) Representative pH and Ca^2+^ maps in TLs and VLs. (**c** and **d**) Histograms of lysosomal pH (c) and Ca^2+^ (d) distribution in VLs of dsDNA treated RAW 264.7 and TLs of Tudor treated cells, (n= 20 cells, m = 300 VLs (n=20 cells; m=100 TLs). (**e**) Representative pH and Ca^2+^ images of *Tudor* treated RAW 264.7 cells in presence of 10 mM ammonium chloride. (**f)** Representative pH and Ca^2+^ images respectively of *Tudor* treated RAW 264.7 showing both VLs and TLs. (**g - h**) Representative images of pH and Ca^2+^ maps of TLs. (**i** and **j**) Schematics of the different TL classes according to their lumenal pH and Ca^2+^ gradients and orientation in the cell. n.g represents no gradient. (**k** and **l**) Quantification of TL populations (A), (B) and (n.g) respectively, (n=20 cells, m= 100 TLs). ***P< 0.0005; **P< 0.005; *P< 0.05 (one-way ANOVA with Tukey *post hoc* test). Error bars represents standard error of mean (s.e.m) from three independent experiments. Scale bar: 10 μm, inset scale bars: 4 μm.

Closer scrutiny of the pH and the Ca^2+^ maps in tubular lysosomes revealed a clear gradient for each ion along the long axis of the tubule (Fig. 4a, b, f-h). In every tubule, areas of high acidity corresponded to low Ca^2+^ and vice versa. In contrast, lumenal pH and Ca^2+^ levels were homogenous within any given vesicular lysosome (Fig. 4b-d). To rule out imaging artifacts, we repeated the experiment, but in the presence of NH4Cl, which neutralizes lysosomal pH and also releases lumenal Ca^2+^ from lysosomes, both of which would be expected to affect tubulation(*65*), and then generated pH and Ca^2+^ maps. NH4Cl treatment dramatically reduced the number of tubular lysosomes overall. However, in the remaining tubular lysosomes both the lumenal pH and Ca^2+^ gradients were dissipated (Fig. 4e and Fig. S19b), which reaffirmed the existence of lumenal pH and Ca^2+^ gradients.

Interestingly, the ion gradients in tubular lysosomes were not static, rather they changed as the tubules underwent growth or deformation such that regions of high Ca^2+^ always correlated with regions of low acidity. In contrast, pH and Ca^2+^ levels in vesicular lysosomes remained constant on similar timescales (Fig. 4g-h). Such Ca^2+^ gradients are consistent with the hypothesis that tubulation requires the stringent control of TRPML1 activity given that experiments by others show that either hyperactivating or inhibiting TRPML1 disintegrates tubular lysosomes(*66*). Our observations suggest that such tight regulation of TRPML1 observed by others, could potentially function to sculpt the Ca^2+^ gradient within tubular lysosomes.

Further analysis revealed at least three kinds of tubular lysosomes existed within cells. Since lysosomes are stretched along microtubules, we found ~95% of tubular lysosomes radiated from the nucleus to the plasma membrane. We therefore considered the nucleus and the plasma membrane as reference points and classified the radially oriented population of tubular lysosomes based on whether their high Ca^2+^ termini were nearer the nucleus (population A) or the plasma membrane (population B). Those tubules that showed no ion gradient were denoted n.g. (Fig. 4i, j). We found that 53% of tubules were oriented such that their high Ca^2+^/low acidity termini were positioned towards the plasma membrane (population B). However, ~29% of tubules were in the reverse orientation with their high Ca^2+^/low acidity termini closer to the nucleus (population A). About ~13% showed no lumenal pH or Ca^2+^ gradients (Fig. 4k, l). Our findings were recapitulated in similar experiments when lysosomes were tubulated with LPS (Fig. S19c-g).

The two different orientations of tubular lysosomes suggest either that there are different mechanisms of tubulation or that there are different kinds of tubular lysosomes or both. Tubulation requires opposing pulls generated by Arl8b-SKIP-kinesin along the plus end of microtubules and Rab7-RILP-dynein complexes along the minus end(*5, 38, 67, 68*). In fact, Arl8b-SKIP and Rab7-RILP complexes are already known to regulate the relative spatial positions of vesicular lysosomes in the cell, i.e., whether they are proximal to the perinuclear region or the cell periphery(*69*). Further, peripheral lysosomes regulate plasma membrane repair and nutrient availability, whereas, perinuclear lysosomes fuse with autophagosomes for autophagy(*66, 70*). It is therefore possible that the two major populations of tubular lysosomes could be associated with different functions.

### Tubular lysosomes promote phagocytosis and phagosome-lysosome fusion

Phagocytosis drives many important functions in innate immune cells including antigen presentation and pathogen killing, amongst others(*71*). It is known that when macrophages are activated with LPS phagocytosis is enhanced(*5, 38, 72*). However, since LPS also tubulates lysosomes, we do not know whether increased phagocytosis is due to immune activation or lysosome tubulation, or both. To test this, we treated cells with either dsDNA, LPS or *Tudor* and measured the rate of phagocytosis of pHrodo™ Red conjugated zymosan by RAW 264.7 cells and Pmac (Fig. 5a and Fig. S20-S21). We found that lysosome tubulation alone was sufficient to enhance phagocytosis in both cellular systems (Fig. 5a, e and Fig. S20-S21). The effect of lysosome tubulation on phagosome maturation was then probed by quantifying tubular lysosome - phagosome contacts and assaying content mixing. Both parameters were quantified by imaging lysosomes and phagosomes labeled with the spectrally distinct tracers Alexa488 Dextran and pHrodo-Red™ Zymosan (SI methods, Fig. 5d). In Pmac, ~30% of all tubular lysosomes in the cell contacted phagosomes while in RAW 264.7 cells it was 15-20% (Fig. 5b-f, h-i, Fig. S22). Further, more physical contacts led to greater content mixing indicating the occurrence of more productive fusion events in both cellular systems (Fig. 5c, g). We also confirmed that *Tudor* treatment did not perturb the fluid phase endocytosis (Fig S20 and S22g-i). Our results show that tubular lysosomes promote phagocytosis and aid phagosome-lysosome fusion.

**Figure 5:**
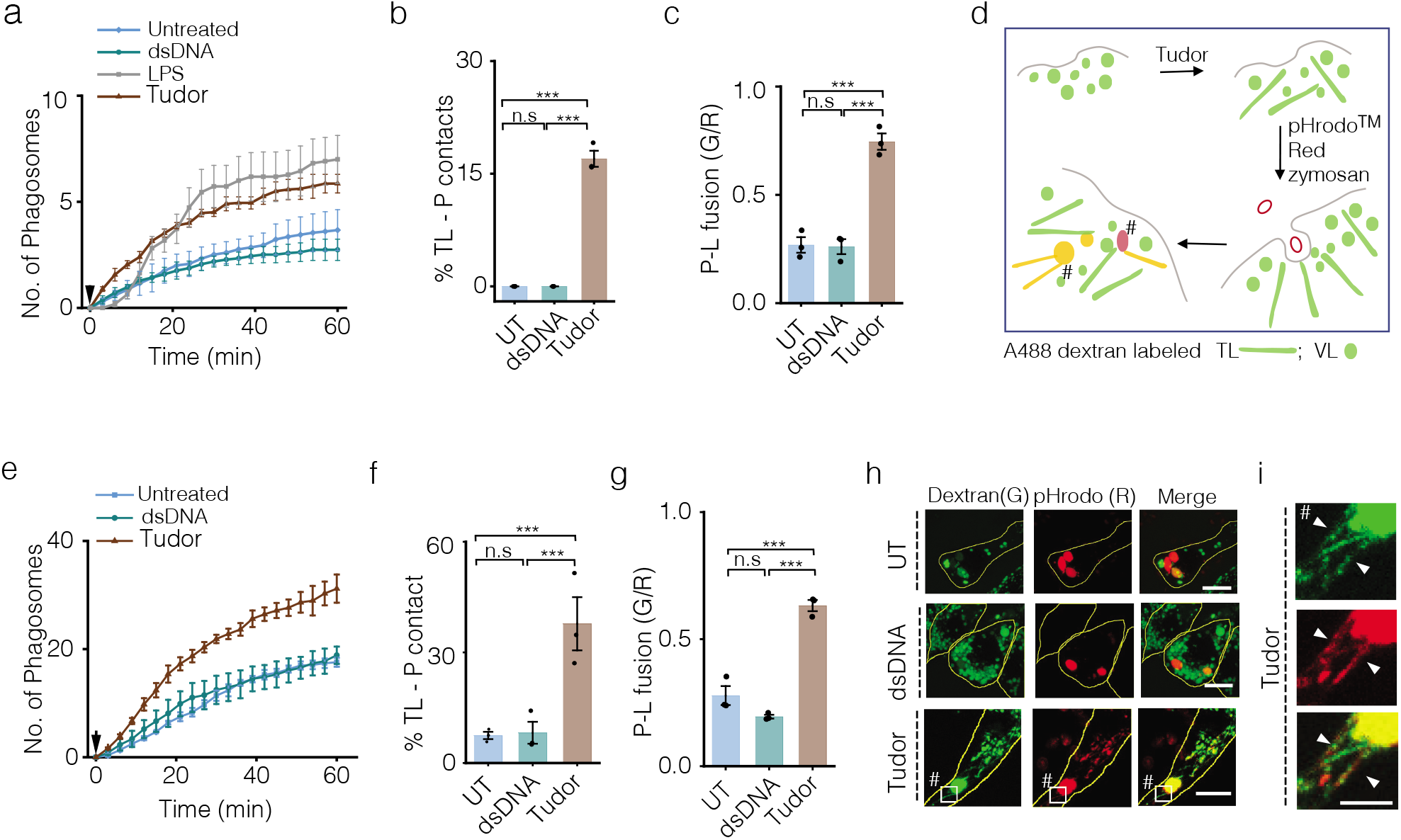
Tubulation promotes phagocytosis and phagosome-lysosome fusion in macrophages. (**a**) Average number of phagosomes as a function of time in RAW 264.7 cells pretreated with dsDNA, LPS or *Tudor*. Arrow shows zymosan addition time (n =30 cells). (**b**) Percentage TLs in contact with phagosome (% TL-P contact) (n =50 cells, m = 100 phagosomes). (**c**) Extent of phagosome lysosome fusion (P-L fusion) represented by mean G/R (n = 60 cells, m=100 phagosomes) in RAW 264.7 cells. (**d**) Schematic of fusion assay in *Tudor* treated cells, lysosomes were labeled with Alexa 488 dextran (G) and phagosomes were labeled with pHrodo™ Red zymosan (R). (**e**) Average number of phagosomes as a function of time in Pmac (M0) pretreated with dsDNA; *Tudor* or LPS. Arrow at t=0 min showing the addition of zymosan on to cells (n = ~30 cells, m=100 phagosomes). (**f**) Percentage TLs in contact with phagosomes (M0) (n = 30 cells, m = 100 phagosomes). (**g**) Extent of phagosome lysosome fusion (P-L fusion) (n =30 cells, m = 100 phagosomes). (**h**) Representative confocal images of lysosomes (G) and phagocytosed zymosan particles (R) in presence or absence of dsDNA; *Tudor* in RAW 264.7 cells (**i**) Inset of TLs contacting phagosomes. # showing TL making contact with phagosome. ***P< 0.0005 (one-way ANOVA with Tukey *post hoc* test). n.s: non-significant; Error bars represents standard error of mean (s.e.m) from three independent experiments. Scale bars: 10 μm and inset scale bars: 4 μm.

### Signal transduction in *Tudor*-mediated enhancement of phagocytosis

Since treating macrophages with *Tudor* tubulates lysosomes without polarizing them, we sought to study how tubular lysosomes enhance phagocytosis in the absence of background transcriptional changes driven by immunostimulation. We know that MMP9 is involved in plasma membrane remodeling which is vital for phagocytosis. The sigmoidal increase in phagocytosis over 4 h triggered by *Tudor* (Fig. 5a, e), suggests a progressive increase of MMP9 activity at the cell surface. We therefore tested whether MMP9 expression increased upon *Tudor* treatment, since MMP9 transcription can be stimulated by either NF-κB or Nrf2. The former can be activated upon immunostimulation(*73, 74*) while the latter is activated during cell starvation or oxidative stress(*11, 75*). Pharmacological inhibition of NF-κB and Nrf2 by JSH-23 and ML385 respectively revealed no impact on tubular lysosome formation induced by *Tudor* (Fig. S23 a-b) (*76, 77*). *Tudor* treated M0 BMDM showed no phosphorylation of NF-κβ or STAT1, reinforcing that *Tudor* does not trigger LPS-like signaling or its associated transcriptional changes (Fig. S23c). Further, *Tudor* treated RAW 264.7 cells showed no change in MMP9 mRNA levels, ruling out transcriptional regulation of MMP-9 (Fig. 6a-c). It is known that PI3K and Akt activation promotes MMP9 secretion. In combination with *Tudor* at the cell surface, which would activate newly secreted MMP9, this may explain a potential feedback mechanism that would also be consistent with the lack of increase in MMP9 at the transcriptional level(*78*).

**Figure 6:**
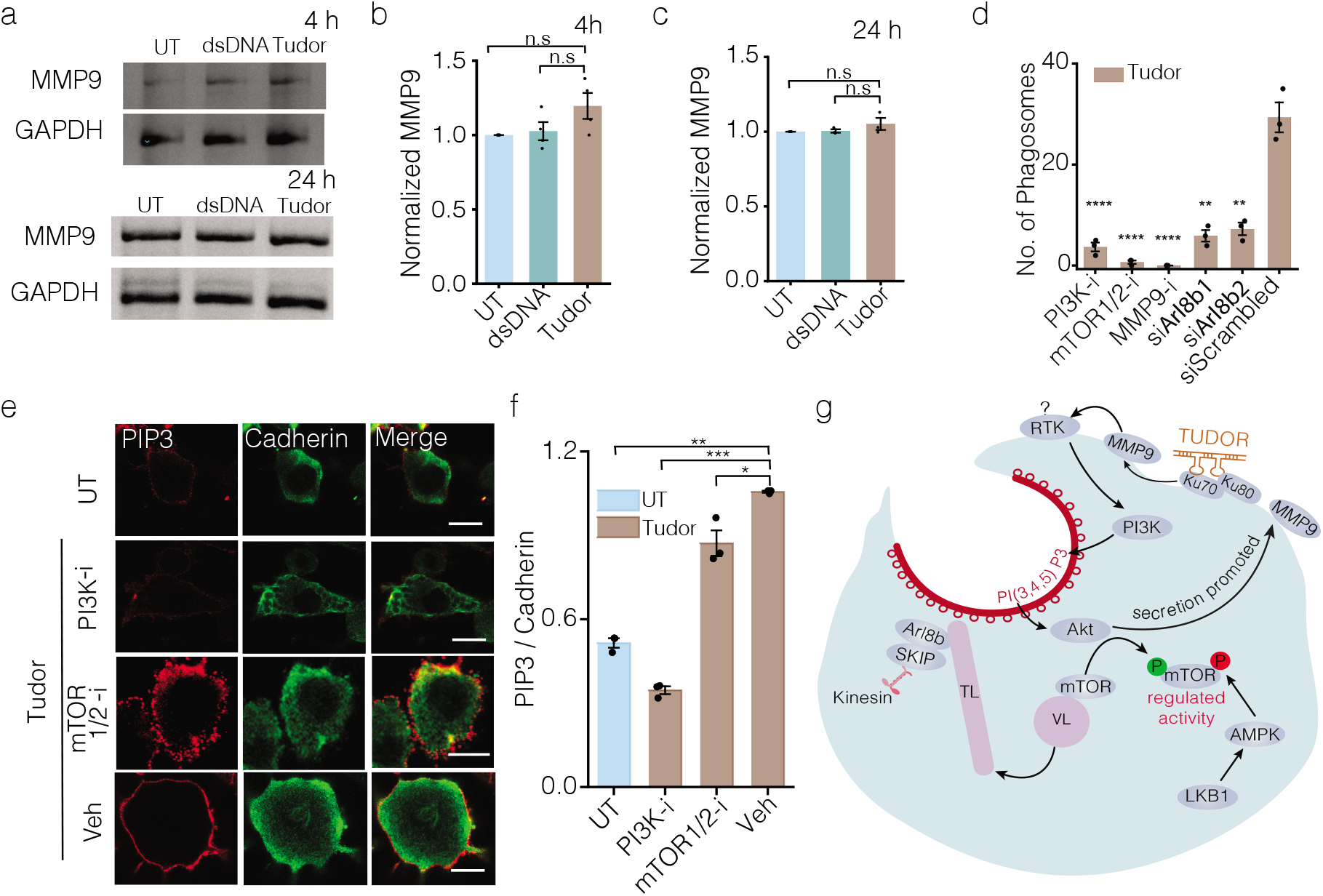
Role of the *Tudor* mediated pathway in enhancing phagocytosis. (**a**) Expression levels of MMP9 in RAW 264.7 cells with or without dsDNA or *Tudor* treatment at indicated times. GAPDH is the loading control. (**b** and **c**) Normalized intensity ratio of MMP9 to GAPDH at (a) 4 hrs and (b) 24 hrs (c). (**d**) Number of phagosomes in *Tudor* treated RAW 264.7 cells in presence or absence of indicated inhibitors and siRNA against *Arl8b*, (n=100 cells). (**e**) Representative confocal images of untreated (UT) or *Tudor* treated RAW 264.7 cells, immunostained for PI(3,4,5)P3 (red) and Cadherin (green) in the presence of indicated inhibitors. (**f**) Ratio of mean total cell intensities of PI(3,4,5)P3 to Cadherin in (e), (n=50 cells). (**g**) Proposed model of signaling pathway underlying how *Tudor* induced tubulation of lysosomes promotes phagocytosis in macrophages. Error bars represent standard error of mean (s.e.m) from three independent experiments. (***P< 0.0005; **P< 0.005; *P< 0.05 (one-way ANOVA with Tukey *post hoc* test).

To examine how tubular lysosomes promote phagocytosis, we treated RAW 264.7 cells with *Tudor* while pharmacologically inhibiting mTOR and PI3K and evaluated the phagocytic capacity of these cells (Fig. 6d). We observed a drastic decrease in phagocytosis in both cases (Fig. 6d). PI3K activation is known to promote phagocytosis by phosphorylating PI(4,5)P2 on inner leaflet of the plasma membrane and converting it to PI(3,4,5)P3, a lipid that facilitates phagosome cup formation(*79, 80*). Indeed, upon *Tudor* treatment, RAW 264.7 cells showed elevated levels of PI(3,4,5)P3 at the plasma membrane, consistent with PI3K activity. Importantly, inhibiting mTOR minimally affected PI(3,4,5)P3 levels at the plasma membrane, yet prevented lysosomal tubulation and impeded phagocytosis (Fig. 6e-f). In order to explicitly test the role of tubular lysosomes in promoting phagocytosis, we knocked down Arl8b using siRNA. The GTPase Arl8b is an adaptor connecting lysosomes to kinesin. It is downstream of mTOR activation and upstream of lysosome tubulation. When RAW 264.7 cells depleted of Arl8b were treated with *Tudor* we found that both lysosome tubulation and phagocytosis were suppressed, despite the rest of the signaling cascade proceeding normally (Fig. 6d and Fig. S24). These results show that tubular lysosomes are needed for effective phagocytic uptake.

### A model for how tubular lysosomes promotes phagocytosis

We therefore propose a model where externally introduced *Tudor* acts as a ligand for Ku70/80 localized on the plasma membrane. When the Ku heterodimer binds *Tudor*, Ku80 activates MMP9. Activated MMP9 is known to interact and activate a range of receptor tyrosine kinases (RTK) at the plasma membrane. Since PI3K has an SH2 domain, we posit that MMP9 could activate PI3K via an as yet unidentified RTK(*81*). PI3K activation in turn, promotes phagocytosis through multiple mechanisms. First, it enriches PI(3,4,5)P3 abundance in the inner leaflet of the plasma membrane to promote the formation of the phagosomal cup(*80, 82*). Second, PI(3,4,5)P3 in turn activates Akt to enhance MMP9 secretion at the cell surface. This sets up a positive feedback loop where *Tudor*-mediated MMP9 activation leads to even more cell-surface MMP9. Finally, PI3K activation leads to mTOR activation via Akt, which makes lysosomes tubulate(*5*), (*81*). Our results show that suppressing any of these processes impedes the early stages of phagocytosis, namely engulfment as well as phagosome lysosome fusion.

Our results also revealed that like lysosomal Ca^2+^ channels, mTOR activity is also stringently regulated and underpins tubular lysosome formation. LKB1 and AMPK activity were found to be important in tubulating lysosomes. AMPK is known to negatively regulate mTOR by phosphorylation at S722 and S792 positions of mTOR(*83, 84*). This likely prevents hyperphosphorylation and runaway activation of mTOR which, in turn, likely toggles lysosomal Ca^2+^ channel activity for effective tubulation (Fig. 6g)(*85*). Recent work shows that lysosomes are also involved in the early steps of phagocytosis in addition to their more well-documented roles in phagosome maturation(*86, 87*). Lysosomal Ca^2+^ channels induce local Ca^2+^ surges that facilitate phagosome formation by activating dynamin or by providing the extra membrane needed for phagocytosis(*86*). With their Ca^2+^-rich termini localized near the cell periphery, tubular lysosomes could facilitate the signaling that leads to cell membrane remodeling.

### Summary

Thus far, tubular lysosomes have been studied either by inducing autophagy or activating immune cells, both of which lead to the cell adopting states with different lysosomal gene expression patterns from that in resting cells. Our studies show that *Tudor* acutely tubulates lysosomes in macrophages without activating them. This allows one to decouple tubular lysosome formation and function from the background transcriptional reprogramming associated with autophagy or immune activation. Although, *Tudor* initiates tubulation without cell starvation or activating TLR receptors, the molecules downstream of PI3K that finally tubulate the lysosome are ultimately the same as in the ALR or LPS-mediated pathways.

The overall proteolytic activity of tubular lysosomes is lower than in vesicular lysosomes, despite their similar cathepsin content. Interestingly, while it is well known that overall proteolytic activity increases upon LPS stimulation(*53*), our studies reveal that this elevation occurs selectively in vesicular lysosomes. Differential proteolysis has also been observed in ALR, where tubular lysosomes are initially devoid of specific cathepsins. Thereafter, they formed proto-lysosomes that were posited to acquire hydrolytic enzymes upon fusion with late endosomes(*7*). The hypoactive proteolysis within tubular lysosomes and the hyperactive proteolysis within vesicular lysosomes indicate that both forms of the lysosome likely perform distinct functions in phagocytosis.

Quantitative pH and Ca^2+^ maps of vesicular and tubular lysosomes revealed that the differential proteolytic activity could not be attributed simply to differences in overall lumenal ionic content. However, we found that tubular lysosomes showed spatial gradients of lumenal pH and Ca^2+^ along their long axis. Such spatial gradients of Ca^2+^ have been previously observed within primary cilia, a sub-cellular structure similar in shape and size to tubular lysosomes(*88*). The tip of the primary cilia maintains high Ca^2+^ levels due to PKD1 channel activity, while at the base of the cilia cytosolic Ca^2+^ acts as a sink. Together this generates an active Ca^2+^ gradient along the long axis of the cilium and enables mechano-sensation(*88*). Specifically in tubular lysosomes, Botelho *et al* have predicted that both pH and Ca^2+^ gradients should arise from continuously toggling the activity of P2X4 and TRPML1(*3*). Our studies provide the first experimental proof of this model(*3*). What the model did not predict, but our observations now reveal is that there is more than one class of tubular lysosomes. We found a major population where their low pH/high Ca^2+^ termini were close to the plasma membrane, a minor population where their low pH/high Ca^2+^ termini were close to perinuclear region, and an even smaller population of tubular lysosomes without ion gradients. Our studies also revealed that the chemical nature and population distributions of tubular lysosomes induced by LPS and *Tudor* are similar.

We found a role for tubular lysosomes in the early stages of phagosome formation by poising resting macrophages for phagocytosis. *Tudor* enhances phagocytosis by activating PI3K downstream, which enriches PI(3,4,5)P3 on the cell membrane, promoting phagosome cup formation(*79, 89*). Macrophages strongly express MMP9 and our studies reveal that MMP9 activation is crucial to phagocytosis. We further show that MMP9 activation proceeds via positive feedback, which ensures the sustained tubulation needed to support the extensive cell membrane ruffling and remodeling needed for phagocytosis. The ruffled border in osteoclasts is actually formed by secretory lysosomes where Snx10 is implicated in transporting secretory lysosomes to the plasma membrane(*90, 91*). In fact, Snx10 is also critical for MMP-9 secretion in osteoclasts(*92*). We therefore suggest that a model where the movement of the much larger tubular lysosomes on force-generating microtubules could similarly push against the fluid, PI(3,4,5)P3-rich cell membrane thereby causing the large-scale remodeling needed to engulf phagocytic cargo.

We were initially surprised that an aptamer to Ku would stimulate phagocytosis. A model to explain this observation is that a nominal amount of Ku on the surface of macrophages could act as a sensor for the ends of fragmented DNA released by dead or dying cells in their proximity. These endogenous triggers of cell-surface Ku could stimulate phagocytosis in order to promote their clearance. Given the different types of tubular lysosome populations, it is possible their roles in cell function may be more widespread than previously anticipated. The ability to switch on or switch off lysosome tubulation using *Tudor* and MMP-9 inhibition respectively in diverse cell types will help uncover new roles of tubular lysosomes and potentially modulate immune cell function.

## Acknowledgements

We thank J. Zou, C. Labno for technical advice; the integrated light microscopy and BioPhysics core facilities at the University of Chicago.

## Funding

This work was supported by the University of Chicago Women’s Board (YK); FA9550-19-0003 from AFOSR (YK), NIH grants R21NS114428 (YK),1R01NS112139-01A1 (YK), R01DK102960 (LB), R01HL137998 (LB), the Mergel Funsky award (YK) and the Ono Pharma Foundation Breakthrough Science Award (YK).

## Author Contribution

B.S, A.S, K.C and YK designed the project. B.S and A.S characterized *CalipHluor 2.0*. A.S performed pH/Ca2+ imaging experiments. K.C synthesized conjugatable Cathepsin C and B probes and qRTPCR. C.C. isolated and polarized primary macrophages from mice and performed qRTPCRs and western blots. B.S performed biochemical/cell biology assays, immunostaining, and imaging experiments. B.S, L.B and YK analyzed the data. B.S, A.S and Y.K wrote the paper. All authors discussed the results and gave inputs on the manuscript.

## Competing interests

The authors declare no competing interests.

## Data availability

The data that support the plots within this paper and other finding of this study are available from the corresponding author upon reasonable request.

## Notes

### Competing Interest Statement

The authors have declared no competing interest.

## References

1. R. M. Perera, R. Zoncu, The lysosome as a regulatory hub. Annu. Rev. Cell Dev. Biol. 32, 223–253 (2016).

2. C.-Y. Lim, R. Zoncu, The lysosome as a command-and-control center for cellular metabolism. J. Cell Biol. 214, 653–664 (2016).

3. G. T. Saffi, R. J. Botelho, Lysosome fission: planning for an exit. Trends Cell Biol. 29, 635–646 (2019).

4. V. E. B. Hipolito, E. Ospina-Escobar, R. J. Botelho, Lysosome remodelling and adaptation during phagocyte activation. Cell Microbiol. 20 (2018), doi:10.1111/cmi.12824.

5. A. Saric et al., mTOR controls lysosome tubulation and antigen presentation in macrophages and dendritic cells. Mol. Biol. Cell. 27, 321–333 (2016).

6. Y. Chen, L. Yu, Recent progress in autophagic lysosome reformation. Traffic. 18, 358–361 (2017).

7. L. Yu et al., Termination of autophagy and reformation of lysosomes regulated by mTOR. Nature. 465, 942–946 (2010).

8. L. Bonet-Ponce et al., LRRK2 mediates tubulation and vesicle sorting from lysosomes. Sci. Adv. 6 (2020), doi:10.1126/sciadv.abb2454.

9. A. E. Johnson, H. Shu, A. G. Hauswirth, A. Tong, G. W. Davis, VCP-dependent muscle degeneration is linked to defects in a dynamic tubular lysosomal network in vivo. Elife. 4 (2015), doi:10.7554/eLife.07366.

10. V. E. B. Hipolito et al., Lysosome expansion by selective translation of lysosomal transcripts during phagocyte activation. BioRxiv (2018), doi:10.1101/260257.

11. Y. Sun et al., Lysosome activity is modulated by multiple longevity pathways and is important for lifespan extension in C. elegans. Elife. 9 (2020), doi:10.7554/eLife.55745.

12. C. V. Harding, H. J. Geuze, Class II MHC molecules are present in macrophage lysosomes and phagolysosomes that function in the phagocytic processing of Listeria monocytogenes for presentation to T cells. J. Cell Biol. 119, 531–542 (1992).

13. M. Boes et al., T-cell engagement of dendritic cells rapidly rearranges MHC class II transport. Nature. 418, 983–988 (2002).

14. J. M. Vyas, A. G. Van der Veen, H. L. Ploegh, The known unknowns of antigen processing and presentation. Nat. Rev. Immunol. 8, 607–618 (2008).

15. N. Nakamura et al., Endosomes are specialized platforms for bacterial sensing and NOD2 signalling. Nature. 509, 240–244 (2014).

16. T. Murakawa, A. A. Kiger, Y. Sakamaki, M. Fukuda, N. Fujita, An Autophagy-Dependent Tubular Lysosomal Network Synchronizes Degradative Activity Required for Muscle Remodeling. BioRxiv (2020), doi:10.1101/2020.04.15.043075.

17. R. Miao, M. Li, Q. Zhang, C. Yang, X. Wang, An ECM-to-Nucleus Signaling Pathway Activates Lysosomes for C. elegans Larval Development. Dev. Cell. 52, 21–37.e5 (2020).

18. K. A. Mullin et al., Regulated degradation of an endoplasmic reticulum membrane protein in a tubular lysosome in Leishmania mexicana. Mol. Biol. Cell. 12, 2364–2377 (2001).

19. J. Swanson, A. Bushnell, S. C. Silverstein, Tubular lysosome morphology and distribution within macrophages depend on the integrity of cytoplasmic microtubules. Proc. Natl. Acad. Sci. USA. 84, 1921–1925 (1987).

20. E. S. Trombetta, M. Ebersold, W. Garrett, M. Pypaert, I. Mellman, Activation of lysosomal function during dendritic cell maturation. Science. 299, 1400–1403 (2003).

21. F. Meng, C. A. Lowell, Lipopolysaccharide (LPS)-induced macrophage activation and signal transduction in the absence of Src-family kinases Hck, Fgr, and Lyn. J. Exp. Med. 185, 1661–1670 (1997).

22. K. Chakraborty, K. Leung, Y. Krishnan, High lumenal chloride in the lysosome is critical for lysosome function. Elife. 6, e28862 (2017).

23. N. Narayanaswamy et al., A pH-correctable, DNA-based fluorescent reporter for organellar calcium. Nat. Methods. 16, 95–102 (2019).

24. K. Leung, K. Chakraborty, A. Saminathan, Y. Krishnan, A DNA nanomachine chemically resolves lysosomes in live cells. Nat. Nanotechnol. 14, 176–183 (2019).

25. A. R. Mantegazza et al., TLR-dependent phagosome tubulation in dendritic cells promotes phagosome cross-talk to optimize MHC-II antigen presentation. Proc. Natl. Acad. Sci. USA. 111, 15508–15513 (2014).

26. R. Levin-Konigsberg et al., Phagolysosome resolution requires contacts with the endoplasmic reticulum and phosphatidylinositol-4-phosphate signalling. Nat. Cell Biol. 21, 1234–1247 (2019).

27. A. R. Mantegazza, J. G. Magalhaes, S. Amigorena, M. S. Marks, Presentation of phagocytosed antigens by MHC class I and II. Traffic. 14, 135–152 (2013).

28. S. Aptekar et al., Selective Targeting to Glioma with Nucleic Acid Aptamers. PLoS One. 10, e0134957 (2015).

29. M. Fukasawa et al., SRB1, a class B scavenger receptor, recognizes both negatively charged liposomes and apoptotic cells. Exp. Cell Res. 222, 246–250 (1996).

30. P. J. Gough, S. Gordon, The role of scavenger receptors in the innate immune system. Microbes Infect. 2, 305–311 (2000).

31. S. Monferran, C. Muller, L. Mourey, P. Frit, B. Salles, The Membrane-associated form of the DNA repair protein Ku is involved in cell adhesion to fibronectin. J. Mol. Biol. 337, 503–511 (2004).

32. E. Weterings et al., A novel small molecule inhibitor of the DNA repair protein Ku70/80. DNA Repair (Amst.). 43, 98–106 (2016).

33. J. Fransson, C. A. K. Borrebaeck, The nuclear DNA repair protein Ku70/80 is a tumor-associated antigen displaying rapid receptor mediated endocytosis. Int. J. Cancer. 119, 2492–2496 (2006).

34. B. S. Prabhakar, G. P. Allaway, J. Srinivasappa, A. L. Notkins, Cell surface expression of the 70-kD component of Ku, a DNA-binding nuclear autoantigen. J. Clin. Invest. 86, 1301–1305 (1990).

35. C. Muller, J. Paupert, S. Monferran, B. Salles, The double life of the Ku protein: facing the DNA breaks and the extracellular environment. Cell Cycle. 4, 438–441 (2005).

36. Y. G. Y. Chan, M. M. Cardwell, T. M. Hermanas, T. Uchiyama, J. J. Martinez, Rickettsial outer-membrane protein B (rOmpB) mediates bacterial invasion through Ku70 in an actin, c-Cbl, clathrin and caveolin 2-dependent manner. Cell Microbiol. 11, 629–644 (2009).

37. J. J. Martinez, S. Seveau, E. Veiga, S. Matsuyama, P. Cossart, Ku70, a component of DNA-dependent protein kinase, is a mammalian receptor for Rickettsia conorii. Cell. 123, 1013–1023 (2005).

38. A. Mrakovic, J. G. Kay, W. Furuya, J. H. Brumell, R. J. Botelho, Rab7 and Arl8 GTPases are necessary for lysosome tubulation in macrophages. Traffic. 13, 1667–1679 (2012).

39. M. A. Gray et al., Phagocytosis enhances lysosomal and bactericidal properties by activating the transcription factor TFEB. Curr. Biol. 26, 1955–1964 (2016).

40. M. S. Jani, J. Zou, A. T. Veetil, Y. Krishnan, A DNA-based fluorescent probe maps NOS3 activity with subcellular spatial resolution. Nat. Chem. Biol. 16, 660–666 (2020).

41. S. Monferran, J. Paupert, S. Dauvillier, B. Salles, C. Muller, The membrane form of the DNA repair protein Ku interacts at the cell surface with metalloproteinase 9. EMBO J. 23, 3758–3768 (2004).

42. J. Paupert, S. Dauvillier, B. Salles, C. Muller, Transport of the leaderless protein Ku on the cell surface of activated monocytes regulates their migratory abilities. EMBO Rep. 8, 583–588 (2007).

43. E.-J. Lee, H.-S. Kim, Inhibitory mechanism of MMP-9 gene expression by ethyl pyruvate in lipopolysaccharide-stimulated BV2 microglial cells. Neurosci. Lett. 493, 38–43 (2011).

44. Y. Zhang et al., Coordinated regulation of protein synthesis and degradation by mTORC1. Nature. 513, 440–443 (2014).

45. D. Kong, T. Yamori, ZSTK474, a novel phosphatidylinositol 3-kinase inhibitor identified using the JFCR39 drug discovery system. Acta Pharmacol Sin. 31, 1189–1197 (2010).

46. J. J. Song et al., c-Cbl acts as a mediator of Src-induced activation of the PI3K-Akt signal transduction pathway during TRAIL treatment. Cell Signal. 22, 377–385 (2010).

47. L. Arbibe et al., Toll-like receptor 2-mediated NF-kappa B activation requires a Rac1-dependent pathway. Nat. Immunol. 1, 533–540 (2000).

48. O. Yamada, K. Ozaki, M. Akiyama, K. Kawauchi, JAK-STAT and JAK-PI3K-mTORC1 pathways regulate telomerase transcriptionally and posttranslationally in ATL cells. Mol. Cancer Ther. 11, 1112–1121 (2012).

49. Y. P. Zhu, J. R. Brown, D. Sag, L. Zhang, J. Suttles, Adenosine 5’-monophosphate-activated protein kinase regulates IL-10-mediated anti-inflammatory signaling pathways in macrophages. J. Immunol. 194, 584–594 (2015).

50. C.-O. Yi et al., Resveratrol activates AMPK and suppresses LPS-induced NF-κB-dependent COX-2 activation in RAW 264.7 macrophage cells. Anat. Cell Biol. 44, 194–203 (2011).

51. D. B. Shackelford, R. J. Shaw, The LKB1-AMPK pathway: metabolism and growth control in tumour suppression. Nat. Rev. Cancer. 9, 563–575 (2009).

52. V. Turk et al., Cysteine cathepsins: from structure, function and regulation to new frontiers. Biochim. Biophys. Acta. 1824, 68–88 (2012).

53. B. M. Creasy, K. L. McCoy, Cytokines regulate cysteine cathepsins during TLR responses. Cell Immunol. 267, 56–66 (2011).

54. T. Jakoš, A. Pišlar, A. Jewett, J. Kos, Cysteine Cathepsins in Tumor-Associated Immune Cells. Front. Immunol. 10, 2037 (2019).

55. K. Dan, A. T. Veetil, K. Chakraborty, Y. Krishnan, DNA nanodevices map enzymatic activity in organelles. Nat. Nanotechnol. 14, 252–259 (2019).

56. B. Korkmaz et al., Therapeutic targeting of cathepsin C: from pathophysiology to treatment. Pharmacol. Ther. 190, 202–236 (2018).

57. Q. Liu et al., Cathepsin C promotes microglia M1 polarization and aggravates neuroinflammation via activation of Ca2+-dependent PKC/p38MAPK/NF-κB pathway. J. Neuroinflammation. 16, 10 (2019).

58. S. Alam et al., Up-regulated cathepsin C induces macrophage M1 polarization through FAK-triggered p38 MAPK/NF-κB pathway. Exp. Cell Res. 382, 111472 (2019).

59. Q. Cao et al., Calcium release through P2X4 activates calmodulin to promote endolysosomal membrane fusion. J. Cell Biol. 209, 879–894 (2015).

60. J. Yang, Z. Zhao, M. Gu, X. Feng, H. Xu, Release and uptake mechanisms of vesicular Ca2+ stores. Protein Cell. 10, 8–19 (2019).

61. X. Wang et al., TPC proteins are phosphoinositide-activated sodium-selective ion channels in endosomes and lysosomes. Cell. 151, 372–383 (2012).

62. H. Xu, M. Delling, L. Li, X. Dong, D. E. Clapham, Activating mutation in a mucolipin transient receptor potential channel leads to melanocyte loss in varitint-waddler mice. Proc. Natl. Acad. Sci. USA. 104, 18321–18326 (2007).

63. R.-J. Li et al., Regulation of mTORC1 by lysosomal calcium and calmodulin. Elife. 5 (2016), doi:10.7554/eLife.19360.

64. P. Li, M. Gu, H. Xu, Lysosomal ion channels as decoders of cellular signals. Trends Biochem. Sci. 44, 110–124 (2019).

65. K. A. Christensen, J. T. Myers, J. A. Swanson, pH-dependent regulation of lysosomal calcium in macrophages. J. Cell Sci. 115, 599–607 (2002).

66. X. Li et al., A molecular mechanism to regulate lysosome motility for lysosome positioning and tubulation. Nat. Cell Biol. 18, 404–417 (2016).

67. W. Du et al., Kinesin 1 drives autolysosome tubulation. Dev. Cell. 37, 326–336 (2016).

68. N. A. Kaniuk et al., Salmonella exploits Arl8B-directed kinesin activity to promote endosome tubulation and cell-to-cell transfer. Cell Microbiol. 13, 1812–1823 (2011).

69. B. Cabukusta, J. Neefjes, Mechanisms of lysosomal positioning and movement. Traffic. 19, 761–769 (2018).

70. D. E. Johnson, P. Ostrowski, V. Jaumouillé, S. Grinstein, The position of lysosomes within the cell determines their luminal pH. J. Cell Biol. 212, 677–692 (2016).

71. M. Guilliams et al., Dendritic cells, monocytes and macrophages: a unified nomenclature based on ontogeny. Nat. Rev. Immunol. 14, 571–578 (2014).

72. V. E. B. Hipolito et al., Enhanced translation expands the endo-lysosome size and promotes antigen presentation during phagocyte activation. PLoS Biol. 17, e3000535 (2019).

73. C. He, Molecular mechanism of transcriptional activation of human gelatinase B by proximal promoter. Cancer Lett. 106, 185–191 (1996).

74. P. K. Shihab et al., TLR2 and AP-1/NF-kappaB are involved in the regulation of MMP-9 elicited by heat killed Listeria monocytogenes in human monocytic THP-1 cells. J. Inflamm. (Lond.). 12, 32 (2015).

75. H. Endo, S. Owada, Y. Inagaki, Y. Shida, M. Tatemichi, Glucose starvation induces LKB1-AMPK-mediated MMP-9 expression in cancer cells. Sci. Rep. 8, 10122 (2018).

76. A. Singh et al., Small Molecule Inhibitor of NRF2 Selectively Intervenes Therapeutic Resistance in KEAP1-Deficient NSCLC Tumors. ACS Chem. Biol. 11, 3214–3225 (2016).

77. H.-M. Shin et al., Inhibitory action of novel aromatic diamine compound on lipopolysaccharide-induced nuclear translocation of NF-kappaB without affecting IkappaB degradation. FEBS Lett. 571, 50–54 (2004).

78. Y. Wang et al., Geraniin inhibits migration and invasion of human osteosarcoma cancer cells through regulation of PI3K/Akt and ERK1/2 signaling pathways. Anticancer Drugs. 28, 959–966 (2017).

79. R. Levin, S. Grinstein, J. Canton, The life cycle of phagosomes: formation, maturation, and resolution. Immunol. Rev. 273, 156–179 (2016).

80. R. S. Flannagan, V. Jaumouillé, S. Grinstein, The cell biology of phagocytosis. Annu. Rev. Pathol. 7, 61–98 (2012).

81. E. Castellano, J. Downward, RAS Interaction with PI3K: More Than Just Another Effector Pathway. Genes Cancer. 2, 261–274 (2011).

82. V. Carricaburu et al., The phosphatidylinositol (PI)-5-phosphate 4-kinase type II enzyme controls insulin signaling by regulating PI-3,4,5-trisphosphate degradation. Proc. Natl. Acad. Sci. USA. 100, 9867–9872 (2003).

83. D. M. Gwinn et al., AMPK phosphorylation of raptor mediates a metabolic checkpoint. Mol. Cell. 30, 214–226 (2008).

84. K. Inoki, J. Kim, K.-L. Guan, AMPK and mTOR in cellular energy homeostasis and drug targets. Annu. Rev. Pharmacol. Toxicol. 52, 381–400 (2012).

85. M. J. Munson et al., mTOR activates the VPS34-UVRAG complex to regulate autolysosomal tubulation and cell survival. EMBO J. 34, 2272–2290 (2015).

86. L. C. Davis, A. J. Morgan, A. Galione, NAADP-regulated two-pore channels drive phagocytosis through endo-lysosomal Ca2+ nanodomains, calcineurin and dynamin. EMBO J. 39, e104058 (2020).

87. X. Sun et al., A lysosomal K+ channel regulates large particle phagocytosis by facilitating lysosome Ca2+ release. Sci. Rep. 10, 1038 (2020).

88. S. Su et al., Genetically encoded calcium indicator illuminates calcium dynamics in primary cilia. Nat. Methods. 10, 1105–1107 (2013).

89. J. G. Marshall et al., Restricted accumulation of phosphatidylinositol 3-kinase products in a plasmalemmal subdomain during Fc gamma receptor-mediated phagocytosis. J. Cell Biol. 153, 1369–1380 (2001).

90. C. Sobacchi, A. Schulz, F. P. Coxon, A. Villa, M. H. Helfrich, Osteopetrosis: genetics, treatment and new insights into osteoclast function. Nat. Rev. Endocrinol. 9, 522–536 (2013).

91. J. Lacombe, G. Karsenty, M. Ferron, Regulation of lysosome biogenesis and functions in osteoclasts. Cell Cycle. 12, 2744–2752 (2013).

92. C. Zhou et al., SNX10 Plays a Critical Role in MMP9 Secretion via JNK-p38-ERK Signaling Pathway. J. Cell Biochem. 118, 4664–4671 (2017).

