## Supplementary Information for "Tubular lysosomes harbor active ion gradients and poise macrophages for phagocytosis"

This File includes

Materials and Methods

Supplementary text

Figure S1 to S24

Tables S1 to S5

Caption for Movie S1

References

**Chemicals and reagents:** All oligonucleotides, were purchased from Integrated DNA Technologies (IDT, USA). Fluorophore labeled oligonucleotides were ethanol precipitated before use. All other oligonucleotides were HPLC purified and used as it is. Oligonucleotides were quantified by UV spectrophotometer (Shimadzu UV-2700), dissolved in milli-q water, aliquoted and stored at -20 °C for further use.

Details of the chemicals used in this study are mentioned in table S4 and S5. Maleylated BSA (mBSA) was synthesized as described in previous protocol(*1*–*3*).

Inhibitors were dissolved in DMSO (67-68-5, Sigma) at concentrations of 3 mM, stored at -20 °C. Myd88 peptide inhibitor kit was dissolved in 1X sterile PBS. For inhibitors studies involving incubations longer than 24 hours; inhibitors were replenished in culture media every 24 hours with the same concentrations. Details of all the inhibitors used in this study are mentioned in table S3.

**Mammalian cell culture:** SIM-A9, COS-7 cells were obtained from American Type Culture Conditions (ATCC). RAW 264.7 macrophages were kind gift from Dr. Christine A. Petersen, Department of Epidemiology, College of Public Health, University of Iowa. J774A.1 were a kind gift from Prof. Deborah Nelson, Department of Pharmacological and Physiological Sciences, University of Chicago. HepG2 cells were kind gift from Dr. Bryan Dickinson, Department of Chemistry, University of Chicago.

RAW 264.7, J774A.1, Cos7 and HepG2 cells were cultured in Dulbecco's modified Eagles medium/F12 (1:1) (DMEM-F12) with 10% FBS as per ATCC protocol. SIM-A9 was cultured in DMEM-F12 with 10% Fetal Bovine Serum (FBS) with 5 % Horse serum (Invitrogen co-operation, USA). DMEM Media were supplemented with 100 U/mL of penicillin, 100 μg/mL of streptomycin (Life Technologies).

**Bone marrow-derived macrophage (BMDM) isolation and activation.** BMDMs were differentiated from bone marrow stem cells with L-cell conditioned media for six days as previously described(*4*). BMDMs were stimulated by LPS (5 ng/mL, Sigma) and INF **γ** (12 ng/mL, R&D Systems) for 24 hrs to be activated to M1 BMDMs, or 20 ng/mL IL-4 for 48 hrs to M2 BMDMs.

**Adipose tissue-macrophages (ATM) isolation.** Adipose tissue was minced and digested with 1 mg/mL type I collagenase in 1% BSA/PBS at 37°C shaker at 160 rpm for 40 minutes. Cell pellet was centrifuged, lysed with red blood cell lysis buffer, and passed through 40 µm filter. ATMs were isolated using CD11b microbeads (Miltenyi Biotec) as previously described(*4*), and purity was assessed by flow cytometry.

**Thioglycolate-elicited peritoneal macrophage (Pmac) isolation.** Pmac were isolated as previously described(*5*). Briefly, Pmac were isolated by lavaging the peritoneal cavity with PBS containing 2% endotoxin-free BSA 5 days after 4% thioglycolate injection (3 mL/mouse).

**Assembly and characterization of *Tudor*:** Equimolar ratios of A1 and A2 oligos were mixed to final concentration of 20 µM in 20 mM sodium phosphate buffer, pH 7.2 containing 10 mM KCl, 10 mM $MgSO_4$. Annealing was done by heating it to 90 °C for 5 mins followed by cooling to RT over 3 hours at the rate of 5 °C/15 mins. This was equilibrated at 4 °C overnight before use. *Tudor* was characterized by mobility shift assay in 12% native poly acrylamide gel electrophoresis (PAGE) (refer Supplementary note. 1)(*6*, *7*).

**Competition assays:**

**Ku70/80 mediated uptake assay:** RAW264.7 were pretreated with 60 equivalence of unlabeled SA43 (aptamer against Ku70/80 heterodimer proteins) in Opti-MEM$^{TM}$ for 30 mins after which cells were treated with 50 nM *Tudor* for 30 mins in Opti-MEM$^{TM}$. Cells were washed and chased for 1 hour in complete media (DMEM containing 10% FBS) containing unlabeled SA43. Cells without SA43 were treated with only the Opti-MEM$^{TM}$ as pre-pulse after which cells were treated with same concentrations of *Tudor* and chased as mentioned above. Imaging was performed as described in methods section.

**Scavenger receptor mediated uptake assay:** RAW264.7 cells were pretreated with 60 equivalence of mBSA for 30 mins in Opti-MEM$^{TM}$ after which cells were treated with 100 nM *Tudor* or dsDNA for 30 mins in Opti-MEM$^{TM}$ containing 60 equivalence of mBSA. Cells were washed and chased for 1 hour in complete medium containing mBSA. Cells without mBSA were pretreated with Opti-MEM$^{TM}$ alone followed by pulse and chase with *Tudor* or dsDNA devoid of mBSA.

All analysis for uptake assays were performed using Fiji(*8*). To quantify uptake, images were background subtracted, whole cell intensity for each cell were measured using cell outlines drawn in the brightfield channel.

**Lysosomal tubulation assay:** RAW 264.7, J774A.1, SIM-A9, HepG2, COS-7 cells; murine ATM; BMDM and Pmac were pulsed with 0.5 mg/ mL TMR dextran for 1 hour and chased in complete media for 16 hours to specifically label lysosomal compartments. Cells were then treated with 100 nM *Tudor* in culture media for 4 hours. Cells were imaged using either a widefield or confocal microscope. LPS (100 ng/mL) was used as a positive control for lysosomal tubulation assay where cells were incubated with LPS for 4 hours at 37 °C in culture media. For time dependent tubulation assay, cells were treated with unlabeled 100 nM *Tudor*; 100 nM dsDNA or 100 ng/ mL LPS (t=0 mins) in complete media. Cells were imaged at different time points (0; 1; 2; 4; 8 and 12 hours) for the formation of tubular lysosomes.

**Fluorescence microscopy imaging:**

IX83 inverted wide field microscope (Olympus Corporation of the Americas) was used with 60x, 1.42 NA or 100X, 1.42 NA, differential interference contrast (DIC) objective (PLAPON, Olympus Corporation of the Americas) and Evolve Delta 512 EMCCD camera (Photometrics). The microscope, filter wheel, shutter, and charge-coupled device camera were controlled using MetaMorph Premier Ver 7.8.12.0 (Molecular Devices LLC, USA). Alexa 488 was imaged with 500/20 band-pass excitation filter, 535/30 band-pass emission filter and 89016 dichroic mirror. Atto 647N was imaged with 640/30 band-pass excitation filter and 705/72 band-pass emission filter with 89016 dichroic. TMR dextran and pHrodo$^{TM}$ Red zymosan was imaged with 530/30 band pass excitation filter with 575/40 band pass emission filter and 49014 long pass dichroic filter. Images in Alexa 488 channel were acquired with 300 ms exposure time and 300 EM Gain. TMR dextran, pHrodo$^{TM}$ red and Atto 647N channel were acquired with 100 ms exposure and 100 ms EM Gain.

The confocal microscope used in the study is Leica TCS SP5 II STED laser scanning confocal microscope (Leica Microsystems, Inc.) with an Argon ion laser for 488-nm excitation, DPS laser for 564-nm excitation and an He-Ne laser for 594-nm, 633-nm excitation, using HCX PIApo 63x/1.4 UV oil 0.14mm WD objective. ER Tracker$^{TM}$ Green, Mito Tracker$^{TM}$ Green, FITC dextran, Alexa 488 dextran, DCF, Rhodamine 110 was excited with Argon laser at 488 nm; TMR Dextran, pHrodo$^{TM}$ Red, Rhod5F was excited using DPSS laser at 561 nm; DQ$^{TM}$ BSA Red was excited with orange HeNe laser 594 nm. Lyso Tracker$^{TM}$ deep red, Alexa 647, Atto 647N was excited using Red HeNe laser 633 nm. Acousto-optical beam splitter (AOBS) was used to filter all emission signals with suitable settings for each fluorophore. Images were recorded using hybrid detectors (HyD).

Time lapse imaging: Time lapse imaging for RAW 264.7 with tubulated lysosomes labeled with *CalipHluore 2.0* was imaged in Leica TCS SP5 II STED laser scanning confocal microscope with 63X, 1.4 NA objective in G, O and R channels. The images were acquired for upto 10 mins with 15 secs time intervals. Images were background subtracted, bleach corrected and processed to construct pH and calcium (log) Images according to previously established procedure(*9*).

**Quantification of Tubular lysosomes:** Tubeness plugin from Fiji(*8*) was used to highlight any curvilinear structures in the images(*10*, *11*). The images were thresholded and Feret value of 0-10 was used to identify all structures between 0-10 μm in length and circularity of 0-0.5 for only tubular structures and circularity of 0-1 is used to identify all tubular and vesicular lysosomes in an image. *Analyze Particles* was used with above mentioned parameters to display the results of area, feret length, intensity for all lysosomes analyzed. Lysosomes of Feret length ≥ 4.0 μm were considered to be a TL(*12*).

Analysis description: (i) %TLs per cell - Number of TLs (≥ 4.0 μm) were divided by total number of lysosomes (VLs + TLs). (ii) % Area of TLs per cell - Mean area of TLs divided by mean area of total lysosomes (VLs + TLs) for each cell.

**Colocalization experiments:** Lysosomes in RAW 264.7 were marked with TMR dextran as mentioned above. Cells were stimulated for tubulation of lysosomes with *Tudor* at 37° C. Cells were then loaded with either 200 nM Mito Tracker™ Green or 50 nM ER Tracker™ Green in HBSSA, incubated for 15-20 mins, washed in HBSS and then imaged in HBSS using Leica TCS SP5 II STED laser scanning confocal microscope.

**Inhibitor assay:** Lysosomes in RAW 264.7, BMDMs were pre-pulsed with TMR dextran as previously mentioned. Cells were then treated with specific inhibitors at 37° C followed by (100 nM) *Tudor* for 4 hours at 37° C in the presence of the inhibitors. Cells were then imaged using a Leica TCS SP5 II STED laser scanning confocal microscope. Details of the concentration and incubation times of each inhibitor used are provided in Table S3.

**Specificity assays:** Equimolar ratios of MUC1-dsDNA; CpG-dsDNA; SA43 aptamer, dsDNA; ssDNA and *Tudor* (refer to Table S1 and S2 for sequence and combinations of DNA used) were annealed as per protocol discussed below at a final concentration of 10 μM in 10 mM sodium phosphate buffer, pH 7.2 containing 100 mM KCl and MgCl.

Lysosomes were marked by TMR dextran, Cells were then treated with (100 nM) MUC1-dsDNA; CpG-dsDNA; SA43, ssDNA, dsDNA and *Tudor* for 4 hours at 37 °C. Cells were then imaged by Leica TCS SP5 II STED laser scanning confocal microscope. Images were background subtracted. Tubeness was used to analyze %TLs/Cells and %Area of TL/cell as discussed above.

**siRNA gene silencing:** siRNA gene silencing in RAW 264.7 were performed using Trans IT-TKO (Mirus Biol LLC) as per supplier's instructions. siRNA used was DsiRNA (IDT DNA, USA) against mouse *Arl8b* (GENE ID: 67166). Two specific siRNA oligonucleotides were used against *Arl8b* along with negative control from DsiRNA for transfection. Complete media was added 15 mins post addition of transfection mixture. Gene silencing was confirmed by quantitative real time PCR for *Arl8b* after 72 hours of transfection. Lysosomal tubulation assay was performed after 72 hours of transfection using the above mentioned protocol.

**MMP9 activity assay:** MMP9 activity assay was performed as per supplier's instructions provided with the MMP 9 assay kit. Cells were seeded at a density of ~100,000 cells per well in 96 well plate and grown with standard culturing conditions. Media was replaced with assay buffer (50 mM Tris, 10 mM $CaCl_2$, 150 mM NaCl, 0.05 % BrijW L23, pH 7.5) containing either APMA (final concentration of 1 mM) for 2 hours; MMP 9inhibitor-I (100 μM for 1 hour) or 500 nM *Tudor* for 4 hours. This was followed by addition of 200 X final concentration of peptide substrate diluted in assay buffer. The substrate containing solution was incubated on cells for 24 hours. The reaction was stopped using stop solution provided in the kit. (Relative fluorescence unit (RFU) was measured using Synergy™ Neo 2 Multi -Mode Microplate Reader with Ex/Em at 480 /520 nm. Mean fluorescence unit (MFU) was calculated and plotted where signal from the APMA containing wells were normalized to 1. Normalized percentage activity where the MFU of background hydrolysis (BH) was subtracted from the other samples (APMA, MMP9-i, dsDNA and *Tudor*) was set to 0 % and APMA to 100 %.

**Measurement of M1 and M2 markers gene expression in BMDM and Pmac by qRT-PCR:** Cell pellets were lysed in RLT buffer and total RNA was isolated using the RNAeasy kit (Qiagen) with on-the-column DNAse digestion (Qiagen). RNA was converted to cDNA using reverse transcription kit (Qiagen), and amplified using QuantiTect SYBR Green PCR Kits (Qiagen). 18S was used as internal control. The primers were used were as follows (F=forward, R= reverse):

| | | |
|---|---|---|
| *18s* | F: GCCGCTAGAGGTGAAATTCTT; | R: CGTCTTCGAACCTCCGACT |
| *Ctb* | F: CTGCGCGGGTATTAGGAGT; | R: CAGGCAAGAAAGAAGGATCAAG |
| *Ctl* | F: AGACCGGCAAACTGATCTCA; | R: ATCCACGAACCCTGTGTCAT |
| *Lamp1* | F: ACATCAGCCCAAATGACACA; | R: GGCTAGAGCTGGCATTCATC |
| *Atp6vod2* | F: CAGAGCTGTACTTCAATGTGGAC; | R: AGGTCTCACACTGCACTAGGT |
| *Tnfa* | F: CACCACGCTCTTCTGTCTACTG; | R: GCTACAGGCTTGTCACTCGAA |
| *Ilb* | F: AACTCAACTGTGAAATGCCACC; | R: CATCAGGACAGCCCAGGTC |
| *Il12* | F: GGAGCACTCCCCATTCCTACT; | R: GAACGCACCTTTCTGGTTACAC |
| *Nos2* | F: GCTCCTCTTCCAAGGTGCTT; | R: TTCCATGCTAATGCGAAAGG |
| *Arg1* | F: CTCCAAGCCAAAGTCCTTAGAG; | R: AGGAGCTGTCATTAGGGACATC |
| *Il10* | F: GCTCTTACTGACTGGCATGAG; | R: CGCAGCTCTAGGAGCATGTG. |
| *Ym1* | F: GCCCACCAGGAAAGTACACA; | R: TGTTGTCCTTGAGCCACTGA. |
| *Cd11b* | F: CCATGACCTTCCAAGAGAATGC, | R: ACCGGCTTGTGCTGTAGTC. |
| Arl8b | F: AGATCTGGGACATAGGCGGA, | R: AGGACCATGTCCTGGGAAGT |

**Western blot analyses**. Cells were lysed with 1% SDS containing protease and phosphatase inhibitors (Sigma), and protein was quantified with the BCA Protein Assay Kit (Pierce). Proteins (10-20 μg) were resolved on 10% SDS-PAGE gels, transferred to PVDF membranes (Millipore), blocked with 5% BSA (Sigma) in 0.1% TBS/Tween-20 at RT for 2hrs, stained with primary and secondary antibodies, and visualized using the ECL detection kit (Biorad) and a LI-COR imager. Antibodies include: pSTAT1 (7649), tubulin (2125), pNF-kB (3033) are from Cell Signaling Technologies.

**RT PCR:** Total RNA was isolated using Trizol as per instructions by manufacturer (Invitrogen). First strand synthesis was performed using Super Script III as per manufacturer's instructions (Thermo Scientific). 10 μL of PCR product was run on 2.0 % agarose gel in TAE buffer.

MMP9 and GAPDH specific primers used were as follows:

| | | |
|---|---|---|
| MMP9 | F: CCTGTGTGTTCCCGTTCATCT, | R: CGCTGGAATGATCTAAGCCCA |
| GAPDH | F: CCCAGAAGACTGTGGATGG, | R: CACATTGGGGGTAGGAACAC |

**Fixation protocols for TLs:**

**3% Glyoxal fixation:** Fixation protocol was modified from previously described method(*13*) Briefly, for 4 mL of total fixative solution; 0.789 mL of absolute ethanol, 0.313 mL 40% glyoxal and 0.03 mL acetic acid were added and the final volume was made up to 4 mL with 1X PBS. pH was set to between 4 and 5 using NaOH. Fixative was prepared freshly just before the experiment. Cells were treated with 3% glyoxal fixative for 20 mins at RT.

**1% Glyoxal Fixation:** Fixative was prepared similar to above mentioned method by just changing the mount of 40% glyoxal solution added to 0.1 mL. Cells were treated with 1% glyoxal fixative exactly for 5 mins at RT.

**0.5% PFA(v/v) +0.45% GA (v/v):** The final concentration 0.5 % PFA (Electron Microscopy Sciences) + 0.45 % of GA (Sigma) and was prepared in 1X PBS. Cells were treated with 0.5 % PFA + 0.45 % GA for 15 mins at RT as reported previously(*14*).

**2% PFA(v/v) + 0.2 % GA (v/v):** The final concentration 2% of PFA and 0.2 % GA was prepared in 1X PBS. Cells were treated with 2 % PFA+ 0.2 % GA for 5 mins at RT.

Post fixation in the above-mentioned fixatives; cells were washed in 1X PBS and imaged.

**Immunofluorescence:**

**Plasma membrane labeling of Ku70:** RAW 264.7 cells were fixed with 2% PFA for 10 mins on ice and gently washed 3 times with ice cold 1X PBS. Cells were blocked with 1% BSA and 0.3 M Glycine in 1X PBS for 30 mins at RT. Cells were labeled with Ku70 in blocking buffer for 1 hour in RT followed by 3 washes in 1X PBS. Cells were then stained with secondary antibody in blocking buffer for 30 mins at RT. Cells were then blocked with 4% FBS+3% BSA in 1X PBS for 30 mins followed by Pan Cadherin antibody overnight incubation. Cells were incubated in secondary antibody for 30 mins post washes. Cells were washed and imaged.

**Cathepsins B labeling in VLs and TLs:** RAW 264.7 were treated with either 100 ng/ mL LPS, 100 nM *Tudor*, dsDNA or culture media (untreated) for 4 hours at 37 °C. Cells were fixed with 2% PFA, 0.2% GA in 1X PBS for 15 mins in room temperature followed by treatment with 0.1% glycine and 3% BSA in 1X PBS for 5 mins, RT. Next, cells were permeabilized with 0.2% Triton$^{TM}$ X100 for 5 mins in 1X PBS and blocked in 4% FBS and 3% BSA in 1X PBS for 2 hours followed by incubation with primary Cathepsin B antibody in blocking buffer overnight at 4 °C in a moist chamber. Cells were then treated with secondary antibody for 1 hour in RT. Again, blocked-in blocking buffer for 2 hours followed by LAMP1 antibody for 1-hour, RT. The secondary antibody was then added at RT for 30 mins. Between every step mentioned above cells were thoroughly washed in PBST. Cells were then imaged.

**Plasma membrane labeling of Phosphatidylinositol (3,4,5) triphosphate:** Cells were treated with 1 μM ZSTK474 for 30 mins; 100 nM Torin 1 for 1 hour followed by 100 nM *Tudor* in presence or absence of the inhibitor for 4 hours in culture media. Cells were then fixed in 5% PFA+0.45% GA for 10 mins at RT and incubated in 100 mM Glycine and 1% BSA for 5 mins. Permeabilization was performed using 0.2% saponin for 3 mins followed by blocking in 5% FBS and 1% BSA for 1 hour. Cells were then incubated with  PIP3 antibody for1 hour at RT, followed by secondary antibody for30 mins, RT. Cells were blocked again with 4% FBS and 3% BSA for 30 mins prior to addition of Pan cadherin antibody overnight at 4° C. Cells were then incubated with secondary antibody for 1 hour, RT, washed and imaged. Between every step mentioned above cells were thoroughly washed in PBST.

Image acquisition for immunofluorescence: Cells were imaged in Leica TCS SP5 II STED laser scanning confocal microscope. All images were processed using Fiji.

**DNA sensors preparation and characterization:**

**Measurement of extinction coefficient of 5(6)-Carboxy-2',7'-dichlorofluorescein (DCF):** DCF was dissolved into dry DMSO to create a primary stock of 50 mM and was stored at -20 °C until used. Different dilutions of DCF were prepared in deionized water and absorption spectra for each were measured using a UV spectrophotometer. Using Beer-Lambert's law, extinction coefficient was estimated from different concentrations of DCF in deionized water and found to be 90000 $M^{-1}cm^{-1}$.

**Conjugation of DCF to DNA and *ImLy2.0* preparation:** DCF was modified with NHS ester according to previous protocol(*15*). 20 µM of the amine labeled 57 base strand (C1) was coupled to DCF-NHS ester (40 eq.), in 20 mM sodium phosphate buffer pH 7.0 and stirred overnight at RT. DCF conjugated DNA was purified by ethanol precipitation(*16*) and quantified using UV-absorption spectroscopy by measuring absorbance at 260 nm for DNA and 504 nm for DCF. The reaction mixture was purified by amicon ultra 0.5 mL centrifugal unit with filter MWCO 3kDa followed by ethanol precipitation to remove any residual free dye. The ethanol precipitated DNA conjugated to DCF was reconstituted in 20 mM Sodium phosphate buffer, pH 7.2. The efficiency of conjugation of DNA to DCF was further confirmed by 20% denaturing PAGE. Once DCF is conjugated to C1 DNA. Equimolar ratios of C1 (DCF containing DNA), C2 (Atto 647 dye containing DNA) and C3 (DBCO- modified DNA) was mixed to final concentration of 10 µM and annealed in 10 mM Sodium phosphate buffer (pH 7.2). The formation of *ImLy 2.0* was confirmed by 15% native PAGE.

***In vitro* fluorescence measurements of *ImLy 2.0:*** *In vitro* calibration for *ImLy2.0* was performed using Fluoromax spectrophotometer (Horiba Scientific) as reported earlier(*17*). Briefly, 30 nM of *ImLy 2.0* was diluted in pH clamping buffer (CaCl2 (50 µ M to 10 mM), HEPES (10 mM), MES (10 mM), sodium acetate (10 mM), EGTA (10 mM), KCl (140 mM), NaCl (5 mM), and MgCl2 (1 mM)) between pH 3.5 and pH 7.2 and allowed to equilibrate at RT for 30 mins. Fluorescence spectra was collected for each sample for DCF (G) by exciting at 504 nM and collecting emission spectra from 512nm to 560 nm and Alexa 647 (R) by exciting at 647 nm and collecting emission spectra from 650 nm to 700 nm. The ratio of emission maxima of G and R which is 520nm:665 nm was measured. The normalized G/R values from three independent experiments were plotted as a function of pH to generate *in vitro* calibration curve.

***In cellulo* clamping of *ImLy 2.0*:** RAW 264.7 were labeled with 500 nM *ImLy 2.0* (DCF: in Opti-MEM™ for 30 min followed by a chase of 30 mins in complete media. Cells were washed, fixed in 4% PFA in 1X PBS for 20mins. After thorough washing, cells were clamped in clamping buffer (120 mM potassium chloride, 5 mM sodium chloride, 1 mM magnesium chloride, 1 mM calcium chloride, 20 mM HEPES, 20 mM MES, 20 mM sodium acetate) at various pH containing 50 µM Nigericin and 50 µM Monensin for 1 hour at RT. Cells were imaged using a widefield microscope. Image analysis: Images were background subtracted and thresholded which was used to obtain ROIs for vesicular lysosomes. The ROIs were applied to background subtracted images of G and R separately. The G values and R values were noted for each lysosome. G/R was plotted for each pH point in each experiment.

***In cellulo* pH measurements by *ImLy 2.0*:** RAW 264.7 pulsed with 500 nM *ImLy 2.0* for 30 mins in Opti-MEM™ followed by chase in complete media for 30 mins. Cells were imaged by wide field microscope in HBSS. Lysosomes in cells were tubulated with *Tudor* followed by incubation with 500 nM *ImLy 2.0* in Opti-MEM$^{TM}$ and a chase of 30 mins in complete media. Cells were washed and imaged in HBSS. Image analysis: Images were background subtracted. Tubeness, a plugin in Fiji was used to highlight any tubular and vesicular structures in the R channel image. The image was then thresholded which was used to obtain ROIs for vesicular and tubular lysosomes. The ROIs were applied to background subtracted images of G and R separately. The G values and R values were noted for each lysosome. G/R was plotted for each pH point in each experiment.

***CalipHluor 2.0* preparation:** 1 mM of Rhod-5F-Azide was conjugated to 10 µM DBCO-C3 in 100 µL of 20 mM sodium phosphate buffer pH 7.2 and stirred overnight at RT(*9*). The reaction mixture was ethanol precipitated to remove any free dye. DNA conjugated to Rhod-5F was reconstituted in 20 mM sodium phosphate buffer pH 7.2. Conjugation was confirmed by 12% denaturing PAGE. *CalipHluor 2.0* was prepared by mixing eq molar concentrations of each oligonucleotides (5 µM) containing DCF, Rhod5F and a ratiometric dye (Atto 647N) in annealing buffer containing 100 mM KCl and 10 mM sodium phosphate buffer pH, 7.2. The formation of *CalipHluor 2.0* was confirmed by gel mobility shift assay in 15% native PAGE.

***In vitro* bead calcium calibration:** The protocol followed for *in vitro* calibration of *CalipHluor 2.0* is as per (*9*). Briefly, 500 nM of *CalipHluor 2.0* was incubated with 0.6 µm monodisperse silica beads in 20 mM sodium phosphate buffer, pH 5.1 containing 500 mM NaCl for 30 mins at RT. The beads were washed thrice by spinning at 10000 rpm for 10 mins each at room temperature. Beads adsorbed with *CalipHluor 2.0* were incubated with clamping buffer (HEPES (10 mM), MES (10 mM), sodium acetate (10 mM), EGTA (10 mM), KCl (140 mM), NaCl (500 mM), and $MgCl_2$ (1 mM)) for 30 mins, RT containing 0.1 µM or 10 mM free calcium buffers at pH (4.0, 4.6, 5.1, 6.0 and 7.2. The 2 µL of beads- *CalipHluor 2.0* solution was imaged on a glass slide in widefield microscope. Rhod-5F(O), Atto 647N (R) and DCF (G) was excited at 545 nm, 647 nm and 504 nm respectively. O/R (calcium) and G/R (pH) from three independent experiments were plotted for each calcium concentrations as function of pH from individual images.

***In vitro* Calcium calibration:** 100 nM of *CalipHluor 2.0* was incubated in calcium clamping buffer (HEPES (10 mM), MES (10 mM), sodium acetate (10 mM), EGTA (10 mM), KCl (140 mM), NaCl (5 mM), and $MgCl_2$ (1 mM)). 0.1 µM and 10 mM free calcium buffers were prepared at pH (4.0, 4.6, 5.1, 6.0 and 7.2). Rhod-5F(O), Atto 647 (R) and DCF (G) was excited at 545 nm, 647 nm and 504 nm respectively. Emission spectra for Rhod-5F, Atto 647N and DCF was collected from 570 nm to 620 nm, 650 to 700 nm and 512 to 560 nm respectively. Mean emission maxima of O/R and G/R from three independent experiments were plotted for each calcium concentrations as function of pH comparing with the *in vitro* bead calibration performed on widefield microscope. Free calcium at given pH for both in vitro calibration was found using https://somapp.ucdmc.ucdavis.edu/pharmacology/bers/maxchelator/CaEGTA-NIST.htm

***In cellulo* pH and Calcium clamping:** *In cellulo* clamping for calcium was performed as mentioned in(*9*). Cells were treated with 500 nM of *CalipHluor 2.0* for 30 mins followed by a chase of 30 mins to make sure the *CalipHluor 2.0* has been targeted to lysosomes. Cells were then fixed in 4% PFA for 20 mins, RT and washed. Cells were incubated in clamping buffer, pH 6.5 containing nigericin (50 µM), monensine (50 µM), ionomycin (20 µM) in clamping buffer containing ethylene glycol-bis(β-aminoethyl ether)-N,N,N′,N′-tetraacetic acid) (EGTA) (10 mM). Cells were incubated with clamping buffer containing 10 mM free calcium for 1 hour, RT. Cells were imaged using confocal microscope. Approximately over 500 endosomes were considered from three independent experiments to compute mean G/R and O/R where G corresponds to mean fluorescence intensity of DCF; O corresponds to Rhod-5F and R to Atto 647N. A pH calibration curve was built using mean G/R from clamped lysosomes at pH 6.5 obtained from *CalipHluor 2.0* and comparing the values with previous calibration curve from *ImLy 2.0*. This calibration curve was used to measure the pH in real time using *CalipHluor 2.0*. O/R values were recorded and considered to be $O/R_{max}$ at pH 6.5.

**pH and calcium measurements:** Cells were either treated with 100 nM *Tudor* for 4 hours to trigger tubulation of lysosomes or treated with 100 nM dsDNA. 500 nM *CalipHluor 2.0* was pulsed and chased of 30 mins such that all lysosomes (TLs and VLs) are marked with *CalipHluor 2.0*. Cells were images in confocal microscope. Quantification and calculation of free calcium in lysosomes of RAW 264.7 were performed according to previously reported method(*9*). For ammonium chloride treatment, lysosomes (VLs and TLs) were pulsed with *CalipHluor 2.0* were chased for 20 mins in DMEM followed by 20 mins of chase in Medium1(M1: 150 mM NaCl, 5 mM KCl, 1 mM $CaCl_2$, 1mM $MgCl_2$, 20 mM HEPES, pH 7.2) buffer containing 10 mM Ammonium chloride at 37 °C. Cells were imaged in confocal microscope in Opti-MEM™ or HBSS.

**Analysis of pH/$Ca^{2+}$ gradient within TLs:** All TLs in a cell is considered for analysis except for those which are parallel to the nucleus. All images were background subtracted. Tubeness plugin in Fiji was used to highlight any tubular and vesicular structures in the R (Atto 647N) channel. Images were then thresholded and used to obtain ROIs for VL and TLs. The ROIs were applied to background subtracted images of G, O and R separately. G/R and O/R images were constructed by dividing G channel image and O channel image with the R channel image. Nucleus was marked with a ROI. A box of 5 X 5 pixels (length and breadth) ROI, which is the average size of a VL was used to measure the G/R and O/R value along the TL starting from the side closest to the nucleus and progressing towards the plasma membrane. The length of 5 X 5-pixel ROI was kept constant throughout the analysis process although width varied based on the width of TLs. The mean intensity of each box was noted as a function of length of the tubule in G, R and O channels and were computed to obtain G/R and O/R values. G/R values were converted into pH using the equation obtained from the pH calibration curve and O/R values were converted into free lumenal calcium concentration using equations established previously(*9*). Both pH and Calcium values were normalized to its respective first value. Normalized pH and calcium values of each TLs were fitted to a straight line to obtain a slope. TLs were segregated based on positive; negative or no change (increase/decrease or no change) given by the slopes of pH and calcium values for each TL. Therefore, TLs were segregated into population A, B or no gradient (n.g).

**Stability of *Tudor* in TLs:** Conjugation of A2-NH2 to DBCO-PEG-ssDNA was performed as per per previously reported literature(*18*). A2-PEG-DBCO was conjugated to azido-Alexa 488 using click chemistry(*19*). Unconjugated azide containing Alexa 488 was removed and DNA was concentrated by amicon ultra 0.5 mL centrifugal filters MWCO 3 kDa (Millipore Sigma). Concentration of Alexa 488 conjugated oligo was measured by UV quantification. 10 μM of Alexa 488 A2 DNA was annealed with Atto-647N labeled A1 in 10 mM potassium phosphate buffer, 100 mM KCl, pH 7.4. Annealing of dual labeled *Tudor* with Atto 647N and PEG-Alexa 488 was performed as mentioned above. Lysosomes in RAW 264.7 were preloaded with 0.5 mg/ mL of TMR dextran. Cells were then pretreated with 100 nM unlabeled *Tudor* for 4 hours for formation of TLs. After 4 hours of incubation with unlabeled *Tudor*, cells were pulsed with 500 nM of dual labeled *Tudor* containing Atto 647N (R) and PEG-Alexa 488(G) for 30 mins and chased over time. Cells were imaged with time in wide field microscope. Image Analysis: Cells were background subtracted. Tubeness from Fiji was used to highlight the tubular lysosomes. ROI generated by analyze particles were used to obtain the G and R mean intensity values respectively. The G and R values were plotted as a function of chase time.

**Stability of dsDNA in lysosomes of RAW 264.7 macrophages:** Conjugation of azide labeled Alexa488 to DBCO-PEG-ssDNA (D1) was performed as reported previously. 10 μM of Alexa 488 ss-DNA (D1) was annealed with 10 μM of Atto 647N (D2) labeled DNA in 10 mM Potassium phosphate buffer, 100 mM KCl, pH 7.4. Annealing of ds DNA was performed as mentioned above. 500 nM of dsDNA was pulsed for 30 mins in RAW 264.7 with lysosomes labeled with (0.5 mg/mL) TMR dextran and chased over time. Cells were imaged at different time points in wide field microscope. Alexa 488 was considered as (G) and Atto 647N (R). Image Analysis: Images were background subtracted. ROI was drawn around the whole cell and whole cell intensities were plotted for both G and R as a function of chase time.

**Preparation of Alexa 488 conjugated dextran:** 2 mg of amino dextran (10 kDa) was mixed with 10 mM of Alexa488 carboxylic succinimidyl ester (Molecular Probes) in final concentration of 20 mM of sodium phosphate buffer, pH 7.2. The mixture was shaken in dark for approximately 8 hours. The excess dye was removed by amicon ultra 0.5 mL centrifugal filter with MWCO 3 kDa. The final concentration and purity of conjugation was quantified by UV spectrophotometer.

**DQ$^{TM}$ BSA assay:** The lysosomes in RAW 264.7 were labeled with Alexa 488 conjugated dextran (3 kDa). Cells were treated with either *Tudor* or dsDNA in complete media and then pulsed with DQ$^{TM}$-BSA red (10 μg/ mL) for 10 mins in HBSS and chased for 30mins in HBSS such that DQ$^{TM}$ BSA is targeted to lysosomes. Cells were again washed and imaged using a confocal microscope.

**Conjugations of azido-Rhodamine110 to DBCO D1 DNA:** 30 μM of DBCO containing D1 DNA was added to 5 equivalence of carboxy rhodamine110 azide in 10 mM sodium phosphate buffer, pH 7.2. The reaction mixture was mixed overnight at RT in dark. Unconjugated dye was removed by ethanol precipitation. Concentration and purity of conjugation was quantified by UV spectrophotometer. Extent of conjugation was also confirmed by 15% denaturing native PAGE. Similar protocol was used for conjugation of azide containing cathepsin C probe to DBCO containing D1 DNA. Success and concentration of conjugation was checked by UV spectrophotometer. Both DNA conjugated with Rhodamine 110 and Alexa 647N containing D2 were annealed as per protocol mentioned above with complementary D2 DNA containing Alexa 647N. Annealed DNA nanostructures ($Cat_{ON}$, $Cat_C$) were confirmed by 12% native PAGE.

**Cathepsin C activity assay:** Cells were pre-treated with either unlabeled 100 nM *Tudor* or dsDNA for 4 hours in complete media. Cells were then labeled with either 500 nM $Cat_C$ or $Cat_{ON}$ in Opti-MEM$^{TM}$ for 30 min followed by a chase of 30 mins at 37 °C in complete media. Cells were washed and imaged in HBSS using a confocal microscope. Image analysis was performed as follows; Images were background subtracted. Alexa 647 channel is considered to be red (R) (excitation maxima $\lambda_{max}$ = 650 nm) and Rhodamine 110 as green (G) (excitation maxima $\lambda_{max}$ = 500 nm). $Cat_{OFF}$ measurement: cells were pretreated with 50 μM of E64 inhibitor for 24 hours. Cells were then treated with 500 nM of $Cat_C$ in presence of 50 μM of E64 for 30 mins. Cells were washes and chased for 30 mins in complete media containing 50 μM E64.

**Image analysis for enzyme activity:** Images were background subtracted. Tubeness plugin in Fiji was used to highlight any tubular and vesicular structures in the R channel image. The image was then thresholded which was used to obtain ROIs for vesicular and tubular lysosomes. The ROIs were applied to background subtracted images of G and R separately. Mean G/R was plotted was computed for each experiment in $Cat_{off}$ ($G/R_{min}$), $Cat_{ON}$ ($G/R_{max}$) and real time measurements of activity of either $Cat_C$ ($G/R_{probe}$) for vesicular and tubular lysosomes. % Response was calculated using the following equation.

$$\% \text{ Response} = (G/R_{probe} - G/R_{min})/(G/R_{max} - G/R_{min})*100$$

**Zymosan pHrodo conjugation:** 5 mg/mL zymosan was freshly dissolved in 10 mM sodium phosphate buffer, pH 7.2 containing 0.2% tween 20 and sonicated for 1 min. 0.5 mg/mL of zymosan was mixed with 100 nM of pHrodo$^{TM}$ Red succinimidyl ester in 10 mM sodium phosphate buffer, pH 7.2 for 4 hours with continuous shaking. The conjugated zymosan was centrifuged at 5000 rpm for 5 mins at RT and stored at 4 °C until used.

**Zymosan uptake assay:** ~80000-100000 RAW 264.7, BMDMs, Pmac were plated in coverslip containing culture dishes. Cells were either treated with 100 nM *Tudor*, 100 nM dsDNA, 100 ng/mL LPS or only culture media (untreated) for 4 hours followed by addition of pHrodo$^{TM}$ Red conjugated zymosan (t=0 min) (excitation maxima $\lambda_{max}$ = 560 nm). Cells were images post addition of zymosan at 37°C over 1 hour using a widefield microscope.

**Imaging conditions for pHrodo$^{TM}$ Red conjugated zymosan and its uptake analysis:** pHrodo$^{TM}$ red conjugated zymosan particles were incubated in universal buffer (UB) ($CaCl_2$ (1 mM), HEPES (20 mM), MES (20 mM), sodium acetate (20 mM), KCl (120 mM), NaCl (5 mM), and MgCl2 (1 mM)) at pH 5.0 for 5 mins. 0.5 µL of this solution was then imaged on glass slide to set up the appropriate imaging conditions. Multiple stage positions were set to image various fields of cells. Zymosan was added (t=0 mins). Cells were then imaged using the above imaging conditions with time intervals of 3 mins up to 60 mins. The images obtained from uptake assay were z-projected with maximum intensity projection. The images were background subtracted in each stack. Each z-stacked image from time t=0 mins upto 60 mins were further stacked together to form a time lapse showing internalizing of zymosan into phagosomes. Number of pHrodo$^{TM}$ Red zymosan particles uptaken into cells were counted with time.

**Zymosan uptake in presence of inhibitors:** RAW 264.7 were cells were treated with PI3K inhibitor (1 µM, Zstk474, 30 mins); mTOR1/2 inhibitor (100 nM Torin1, 1 hour); MMP9 inhibitor (100 µM, MMP9-I, 1 hour) or siRNA against *Arl8b* for 72 hours. Cells were then treated with 100 nM *Tudor* for 4 hours in presence or absence of inhibitor, scrambled or siRNA against *Arl8b*. Cells were then pulsed with pHrodo$^{TM}$ Red conjugated zymosan for 30 mins at 37 °C. Cells were imaged to score for internalized zymosan. Zymosan uptake was analyzed for ~100 cells in each condition.

**Phagosome lysosome fusion assay:** Cells were treated with 2 mg/mL, 10 kDa Alexa 488 conjugated dextran with 1 hour of pulse and chased overnight to mark all lysosomes. Cells were then either treated with *Tudor*, dsDNA or only culture media (untreated) followed by the addition of 0.5 µL pHrodo$^{TM}$ Red conjugated zymosan for 30 mins at 37 °C and imaged by confocal microscopy. Imaging conditions before each experiment were set as mentioned above. Image Analysis: All images were background subtracted. Alexa 488 dextran was considered to be green (G) and pHrodo$^{TM}$ Red was considered to be red (R) and the fusion of phagosome to lysosome was analyzed in single plane confocal images where the ROI was drawn in the R channel using Fiji. The same ROI was used to obtain intensity values of both G and R channels and ratios were plotted for fusion. For phagosome lysosome contacts analysis; The number of TLs making contact with a single phagosome out of total Alexa 488 dextran containing TLs in a cell were counted.

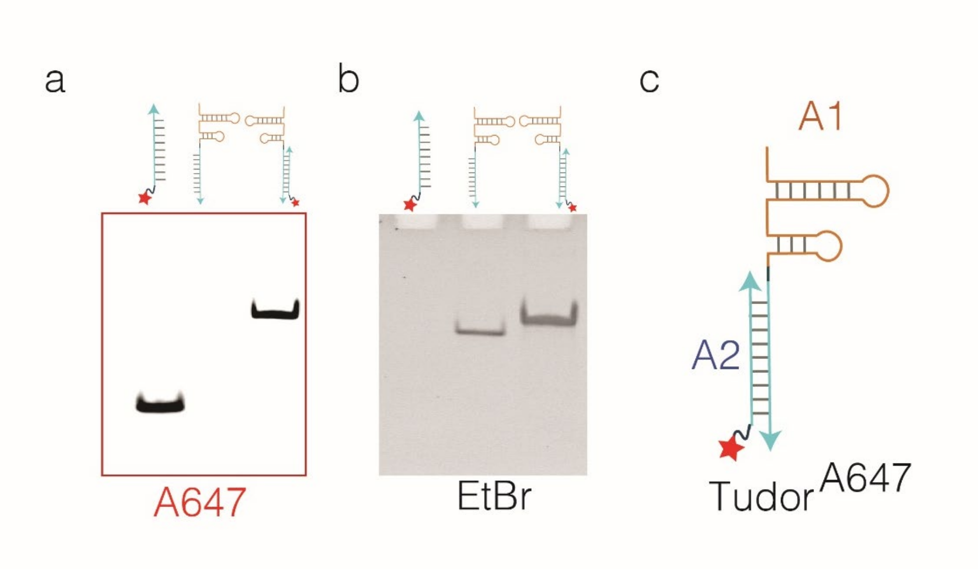


**Figure S1: Characterization of *Tudor*. (**a, b**)** Gel mobility shift assay characterizing the assembly of *Tudor* using 10% Native PAGE. Imaged in A647 (red) and EtBr (black) channels. Lane 1 showing the mobility of A2; Lane 2: A1 and Lane 3: equimolar A1 and A2 annealed product as indicated in the schematic of *Tudor* as shown in (c).

**Supplementary Note 1: Gel characterizations.** The formation of *Tudor* was confirmed by gel mobility shift assay with Native Polyacrylamide gel electrophoresis (PAGE) (Fig S1 a, b). *Tudor* consists of two single stranded oligonucleotides, namely, A2 strand: cyan strand containing Alexa 647 dye and A1 strand: orange strand contains the aptamer, SA43 which binds to Ku70/80 heterodimer on the cell surface followed by a trimer linker into (A3) sequence complementary to A2 (Fig S1c). A1 and A2 oligonucleotides and *Tudor* were stained with EtBr and imaged in both A647 and EtBr channels. A1 showed lower mobility shift compared to A2 in both A647 and EtBr channels while *Tudor* showed higher mobility shift compared to A1 and A2 strands.

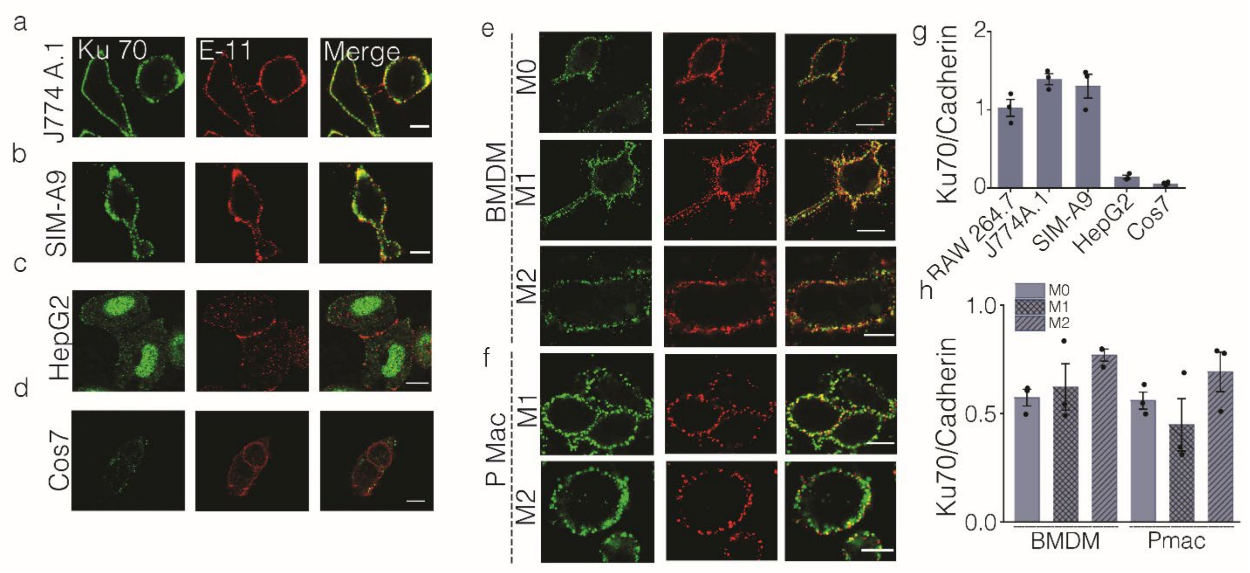

**Figure S2: Ku70 localizes on the plasma membrane of various cell lines and primary macrophages. (a-f)** Representative images showing colocalization of Ku70 (green) with Pan Cadherin (E-11, red) in (a) J774A.1; (b) SIM-A9; (c) HepG2; (d) COS-7; (e) naïve (M0), LPS/INFg activated- (M1), or IL4-activated (M2) BMDM and (f) naïve (M0), LPS/INFg activated- (M1), or IL4-activated (M2) Pmac, Scale bar = 10 µm. (g, h) Normalized intensity ratio of Ku70/Pan Cadherin for each indicated cell types. Data represents three independent experiments shown here (n=50 cells).

**Supplementary Note 2. Plasma membrane localization of Ku70 protein in different cell types:** Ku70/Ku80 heterodimers which is a DNA repair protein, performs nonhomologous end joining (NHEJ) in nucleus and are also found at the plasma membrane in certain cancer and immune cells(*20–23*). DNA aptamer SA43 was raised against the Ku70/Ku80 heterodimers present on the plasma membrane(*24*). The presence of Ku70 protein on the surface of plasma membrane was confirmed by immunofluorescence without permeabilization in various cell lines mentioned above (Fig S2).

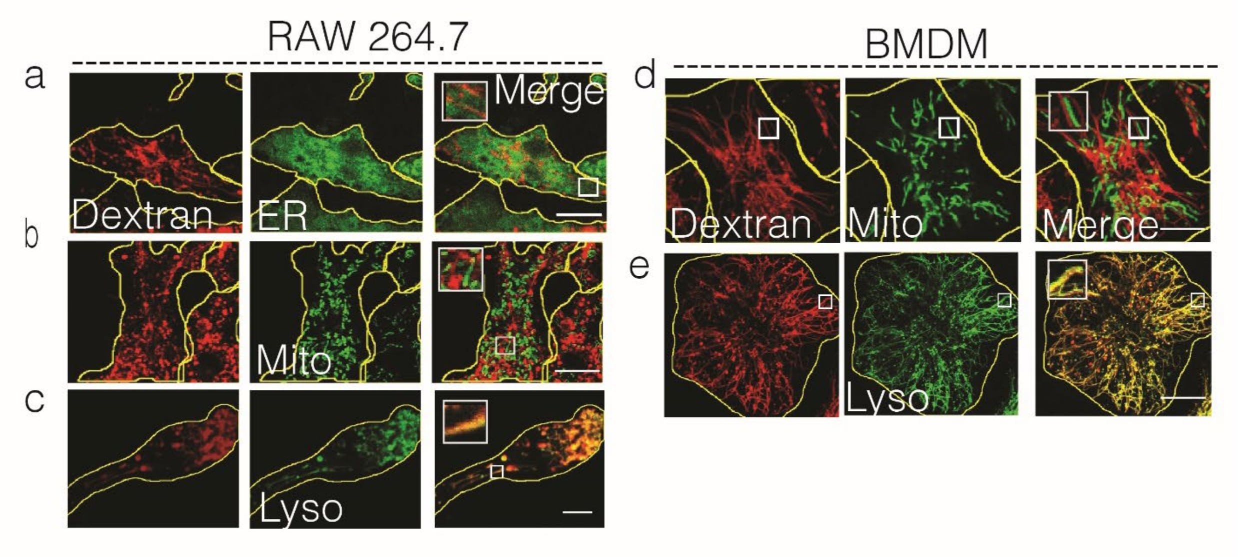


**Figure S3**: ***Tudor* specifically mediates tubulation of lysosomes.** Representative colocalization images of TMR-dextran labeled tubular lysosomes (red) with indicated organelle markers [green; (ER) – ER Tracker$^{TM}$; Mitochondria (Mito)-MitoTracker$^{TM}$ green; Lysosomes (Lyso) – LysoTracker$^{TM}$] in *Tudor* treated RAW 264.7 (a-c) and BMDM (d, e). All independent experiments were repeated at least three times. Scale bar = 10 µm.

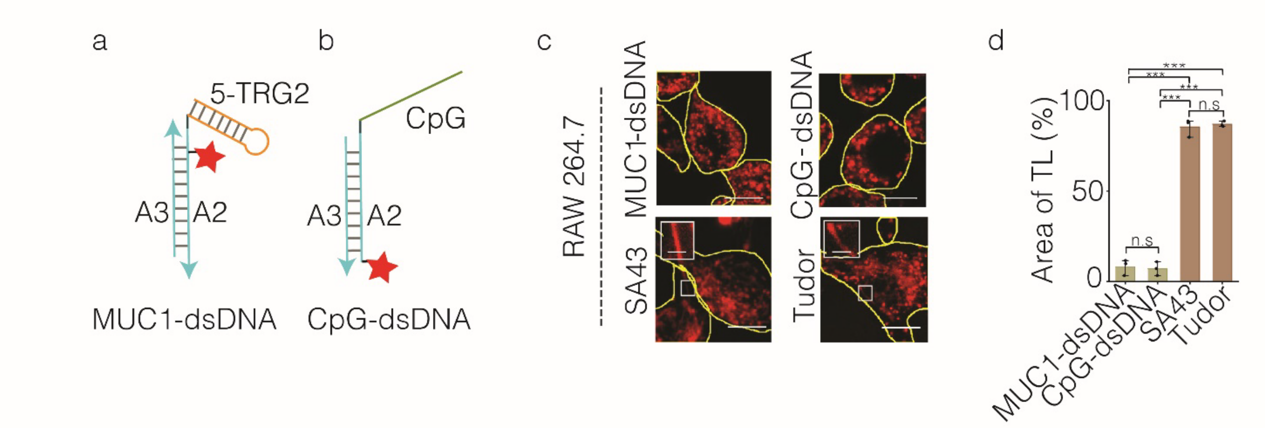


**Figure S4: Lysosomal tubulation is specifically triggered by *Tudor*.** (a and b) Schematic showing MUC1-dsDNA made of 5-TRG2 linked to A2 DNA complementary to A3 and CpG-dsDNA where CpG strand is linked to A2 DNA complementary to A3. (c) Representative confocal images of RAW 264.7 in presence of MUC1-dsDNA, CpG-dsDNA, SA43 aptamer and *Tudor*. Scale bar: 10 µm, inset scale bar: 4 µm. (d) Normalized % Area of TLs over total lysosomes in presence of indicated ligands. Error bars represent s.e.m from 3 independent experiments, (n= 20 cells per experiment); ***P< 0.0005; **P< 0.005 (one-way ANOVA with Tukey *post hoc* test). n.s: non-significant.

**Supplementary note 3: SA43 aptamer trigger tubulation of lysosomes.**

MUC1-dsDNA which was adopted from the prior art(*25*) incorporates 5-TRG2, a DNA aptamer which binds to hypo-glycosylated MUC-1 protein (*K*d: 18 nM) upregulated on the plasma membrane of certain cancer cells(*26*). CpG-ODN, a TLR-9 ligand, can trigger innate immune response in mammalian cells(*27*). Briefly, 5-TRG2 aptamer is fused to 24mer DNA (A2) through a short tri-mer oligonucleotide linker. 5-TRG2 fused to A2 strand along with its complementary A3 strand forms MUC1-dsDNA. CpG-dsDNA was also adopted from prior work(*28*) where CpG strand is also linked through short tri-mer linker to 24mer (A2) oligonucleotide which is complementary to A3 strand forming CpG-dsDNA. CpG-dsDNA is visualized by Alexa 647N present on A2 strand while MUC1-dsDNA was visualized by Alexa 647N present internally on A2 strand. *Tudor*, MUC1-dsDNA and CpG-dsDNA have similar design which involves a single strand overhang followed by a dsDNA module whose length and sequence are similar. To confirm the specificity of *Tudor* in triggering tubular lysosomes we treated RAW264.7 with 100 nM of MUC1-dsDNA, CpG-dsDNA, SA43 aptamer and *Tudor* for 4 hours. SA43 and *Tudor* treated cells showed lysosomes predominantly tubulated compared to MUC1-dsDNA and CpG-dsDNA treated cells suggesting that it's SA43 aptamer which is internalized by Ku70/80 on the plasma membrane of the macrophages to trigger tubular lysosome formation.

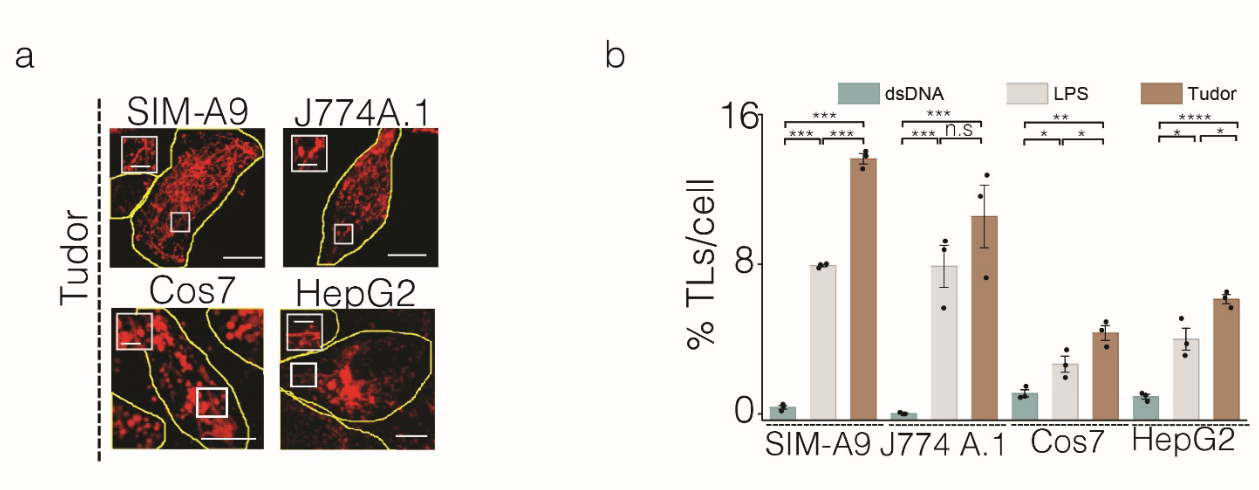


**Figure S5: *Tudor* triggers tubulation of lysosomes in various cell types.** (a) Representative fluorescence images of TMR dextran labeled lysosomes in *Tudor* treated SIM-A9, J774 A.1, Cos7 and HepG2 cells. Scale bar: 10 µm; inset scale bar: 4 µm. (b) Quantification of % TLs per cell on indicated cell types in presence of dsDNA, LPS or *Tudor* Error bars represent s.e.m from 3 independent experiments, n= 20 cells per experiment; ***P< 0.0005; **P< 0.005 (one-way ANOVA with Tukey *post hoc* test). n.s: non-significant.

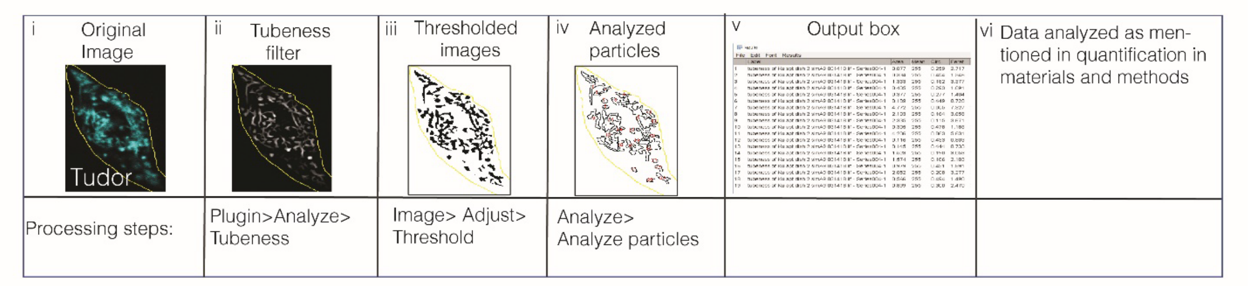


**Figure S6: Image analysis framework for quantification of tubular lysosomes.** (i) Fluorescent images of *Tudor* treated cells were background subtracted. (ii) the image was subjected to Tubeness filter which highlights all curvilinear structures. (iii) image in (ii) was then converted into a binary image by thresholding (0, 255). (iv) image in (iii) was used to find all structures (VLs and TLs) using analyze particles in Fiji based on two parameters: Feret values (0-10) and circularity (range:0.0-0.5). (v) Tubular structures only ≥4 µm considered for statistical analysis and quantification. (vi) The data obtained were analyzed in multiple methods, %TLs per cells, Number of TLs per cell, % Area of TLs per cell.

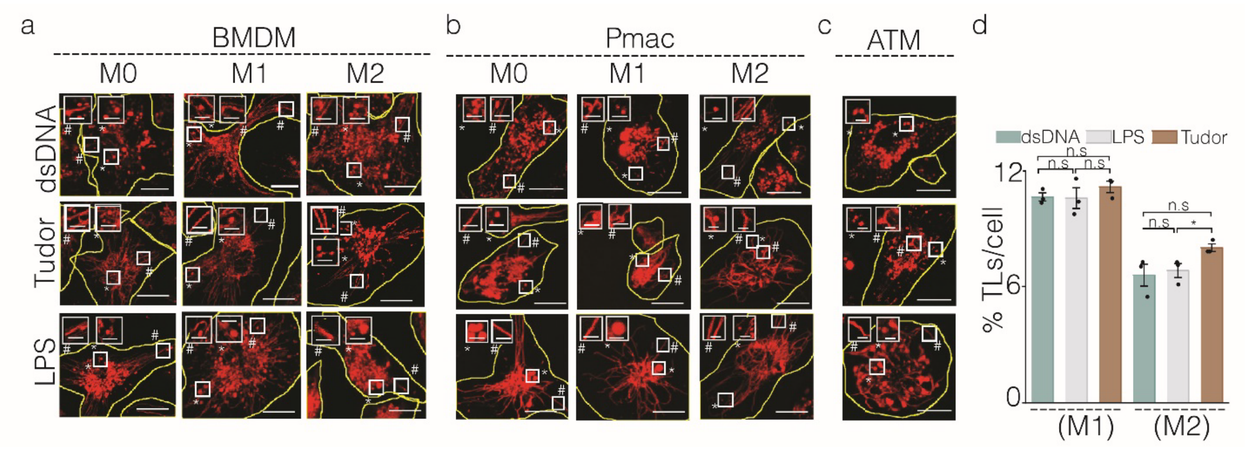


**Figure. S7: *Tudor* tubulates lysosomes in murine primary macrophages.** (a) Representative confocal images of TMR dextran labeled lysosomes in BMDMs**,** (b) Pmacs and **(c)** ATMs upon treatment with dsDNA, LPS or *Tudor*. Inset magnified image of section shown in the white box with * representing VLs and # representing TLs. Scale bar: 10 µm, inset scale bar: 4 µm. (d) Quantification of % TLs per cells for M1 and M2 macrophages of BMDM (n = 50 cells), error bars represent s.e.m from 3 independent experiments.

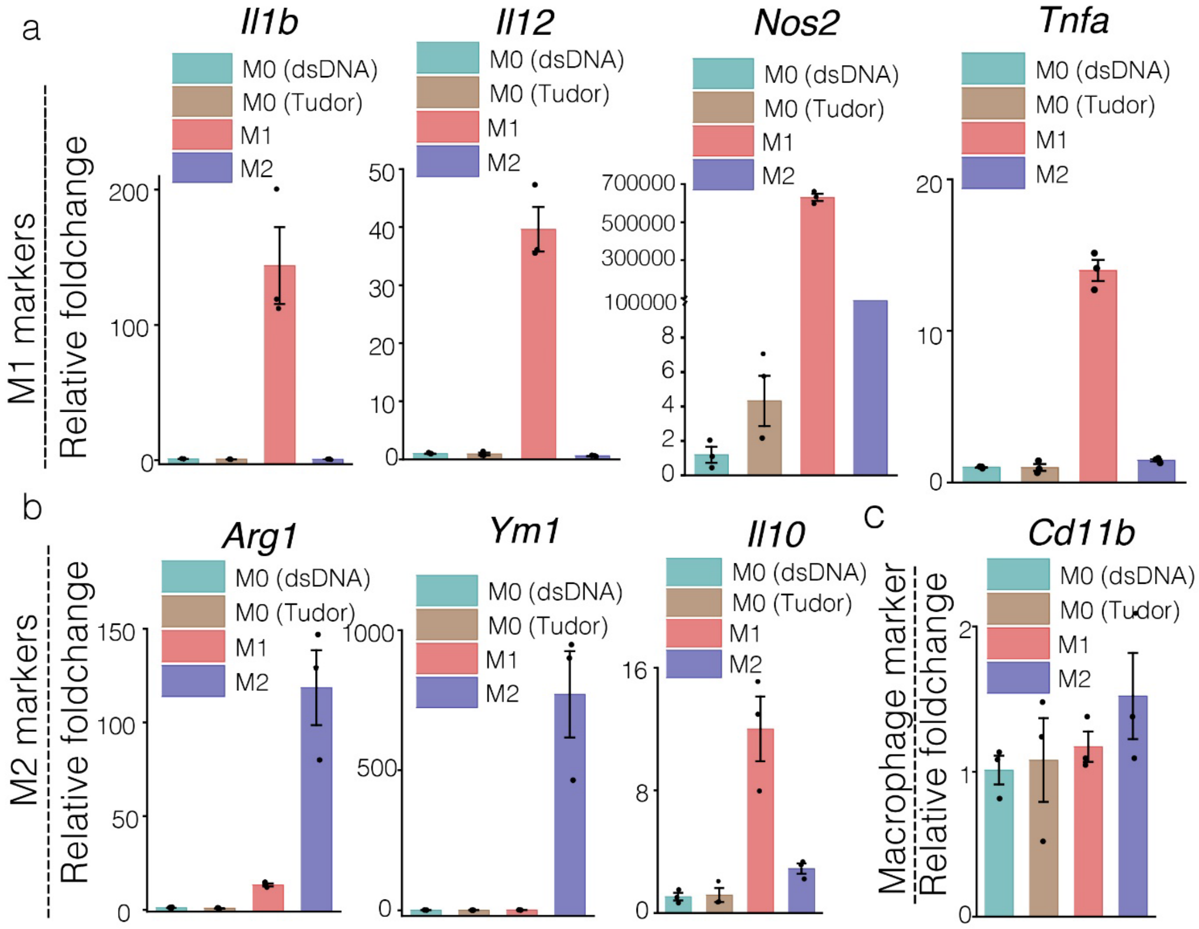

**Figure. S8: mRNA expression profiles of *Tudor* or dsDNA treated Pmac (M0). (a, b** and **c)** Expression levels of M1 (a) and M2 (b) and macrophage marker (c) genes shown in Pmac (M0) upon dsDNA and *Tudor* treatment. All error bars represent s.e.m from three independent experiments.

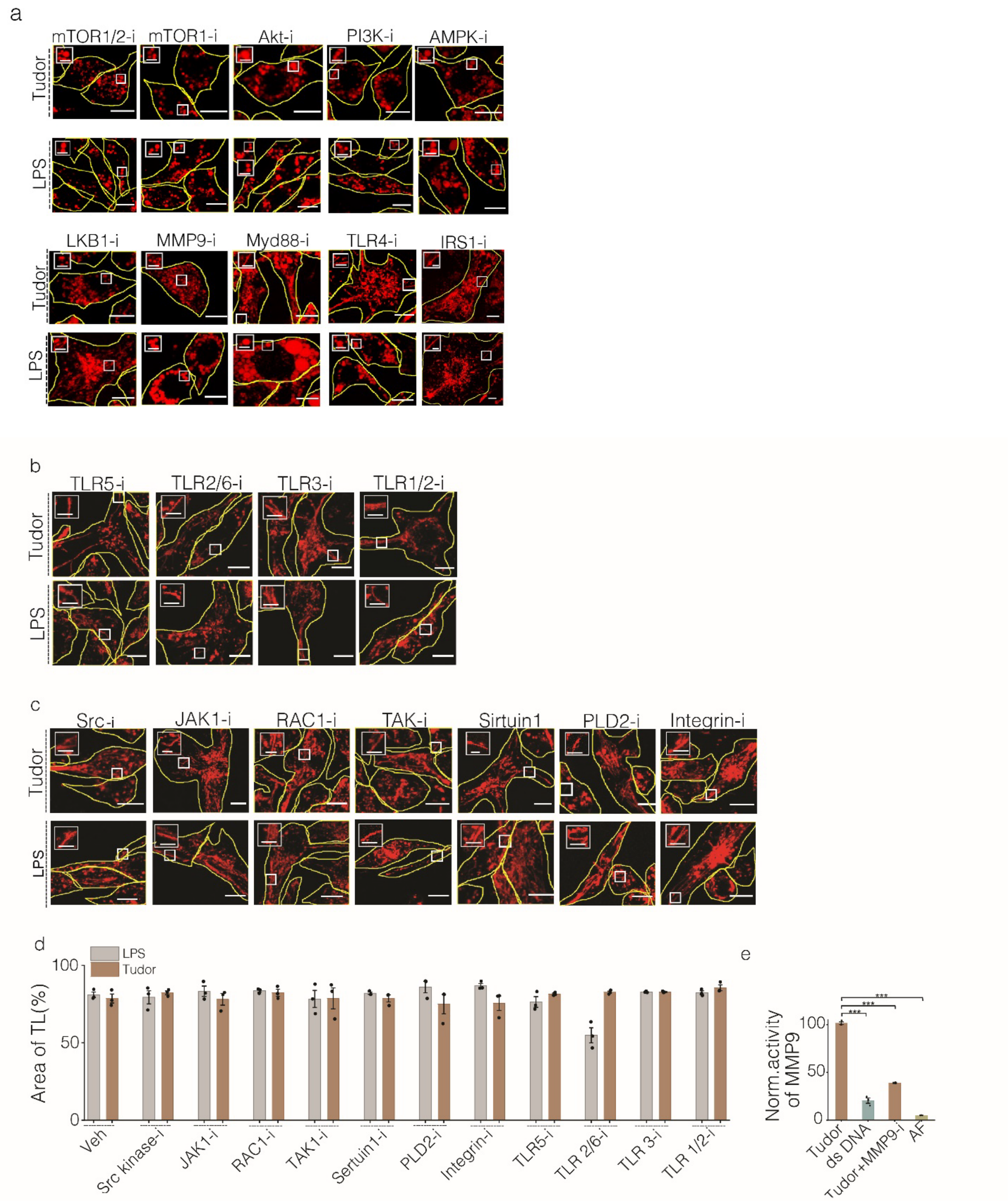


**Figure. S9: Pharmacological perturbations in Tudor or LPS treated RAW 264.7 cells.** (a-c) Representative confocal images of TMR dextran labeled lysosomes in *Tudor* or LPS treated cells in the presence of indicated pharmacological inhibitors. Scale bar: 10 µm, inset scale bar: 4 µm. (d) Quantification of mean % Area of TL over total lysosomes shown in (b-c), (n=20 cells), (Veh=DMSO). (e) Normalized activity of MMP9 in RAW 264.7 upon treatment with *Tudor* (in absence or presence of MMP9 inhibitor-1) and dsDNA where mean fluorescence unit of *Tudor* was normalized to maxima (100%). AF represents autofluorescence of cells without any treatment. ***$P < 0.0005$; (one-way ANOVA with Tukey *post hoc* test). Error bars represent s.e.m from three independent experiments.

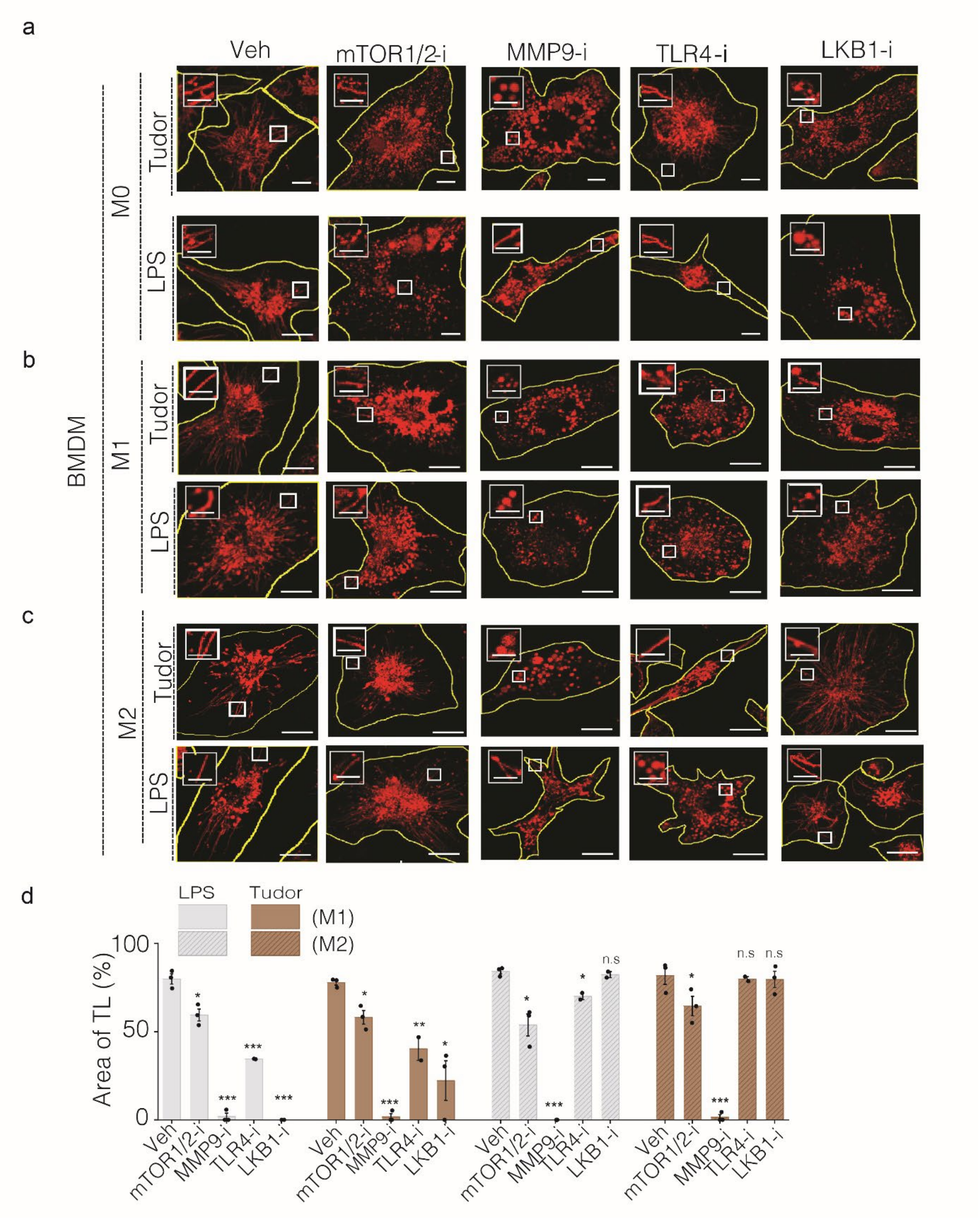


**Figure. S10: Lysosome tubulation pathway triggered by *Tudor* is conserved in BMDMs.** (a-c) Representative confocal images of TMR-dextran-labeled lysosomes of murine BMDMs treated either with *Tudor* or LPS where the indicated proteins are pharmacologically inhibited. (d) % Area of TL over total lysosomes shown for M1 (left) and M2 (right) obtained from (b and c). ****P*< 0.0005; **P< 0.005; *P< 0.05 (one-way ANOVA with Tukey *post hoc* test). Veh=DMSO; n.s: non-significant. Error represents s.e.m from three independent experiments with n= 20 cells per experiment. Scale bar: 10 µm, Inset scale bars: 4 µm.

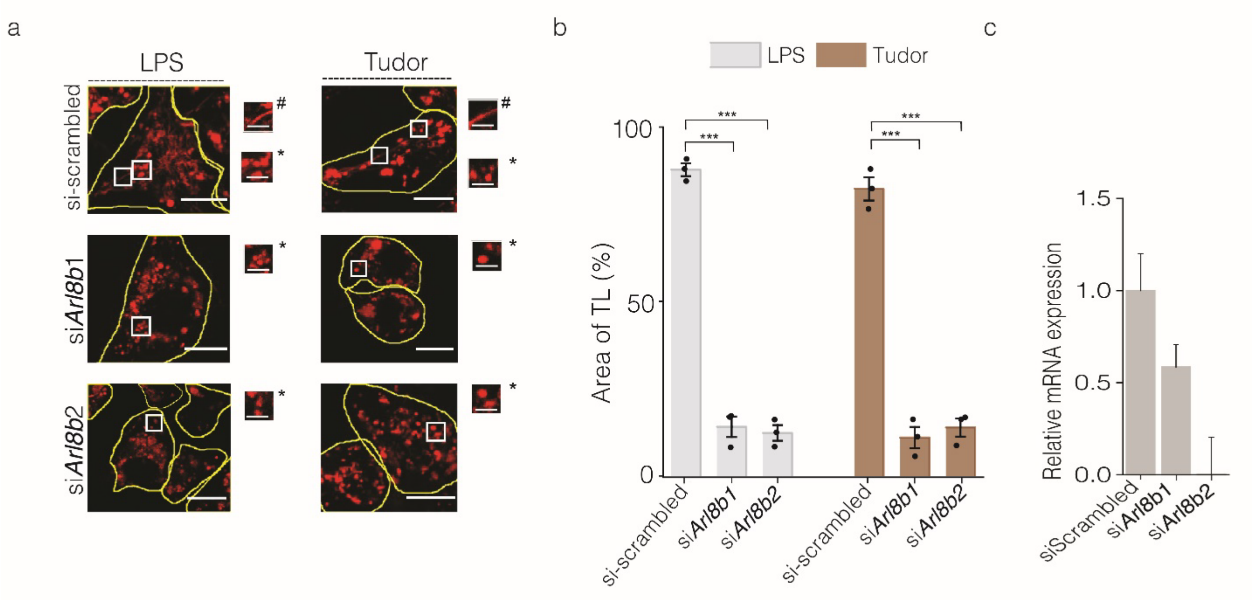


**Figure. S11: Arl8b regulates *Tudor* mediated tubulation of lysosomes.** (a) Representative images of TMR dextran labeled lysosomes in *Tudor* or LPS treated RAW 264.7 transfected with siRNA against *Arl8b* (si*Arl8b1* and si*Arl8b2*). Scale bar: 10 µm, with inset scale bar: 4 µm. * in inset represent VLs; # represents TLs. (b) Quantification of % Area of TLs as a measure of extent of lysosomal tubulation upon *Tudor* and LPS treatment and *Arl8b* knockdown, (n=20 cells). ***$P< 0.0005$; (one-way ANOVA with Tukey *post hoc* test). (c) Relative mRNA expression levels in RAW 264.7 treated with siRNA against *Arl8b* (si*Arl8b1* and si*Arl8b2*) and scrambled siRNA. Error bars represent s.e.m. from 3 three independent experiments.

**Supplementary Note 4: Arl8b is essential for *Tudor* triggered tubulation of lysosomes:** Lysosomal motility protein, Arl8b (ADP Ribosylation Factor like GTPase 8b) is a small Arf like GTPase which regulates the lysosomal positioning within the cytosol. Arl8b aids in lysosomal movement towards the periphery of the cell by governing the motility of lysosomes towards the "+" end of microtubule. This is due to its interaction with motor protein Kinesin1, through an adapter protein, SifA Kinesin interacting protein (SKIP)(*29*). Role of Arl8b in formation and movement of LPS triggered TLs is previously demonstrated(*12*, *30*). To study if *Tudor* trigged TLs formation also involved the recruitment of Arl8b was studied in RAW 264.7 where the cells were transfected with siRNA specific to Arl8b. Lysosomes in these cells were marked with TMR-dextran followed by treatment with *Tudor* to trigger tubulation of lysosomes. TLs in these cells were scored using Tubeness plugin as describes in Methods (Fig. S6). Cells treated with siRNA for Arl8b showed drastically reduced TLs formation suggesting the involvement of Arl8b in *Tudor* triggered TL formation.

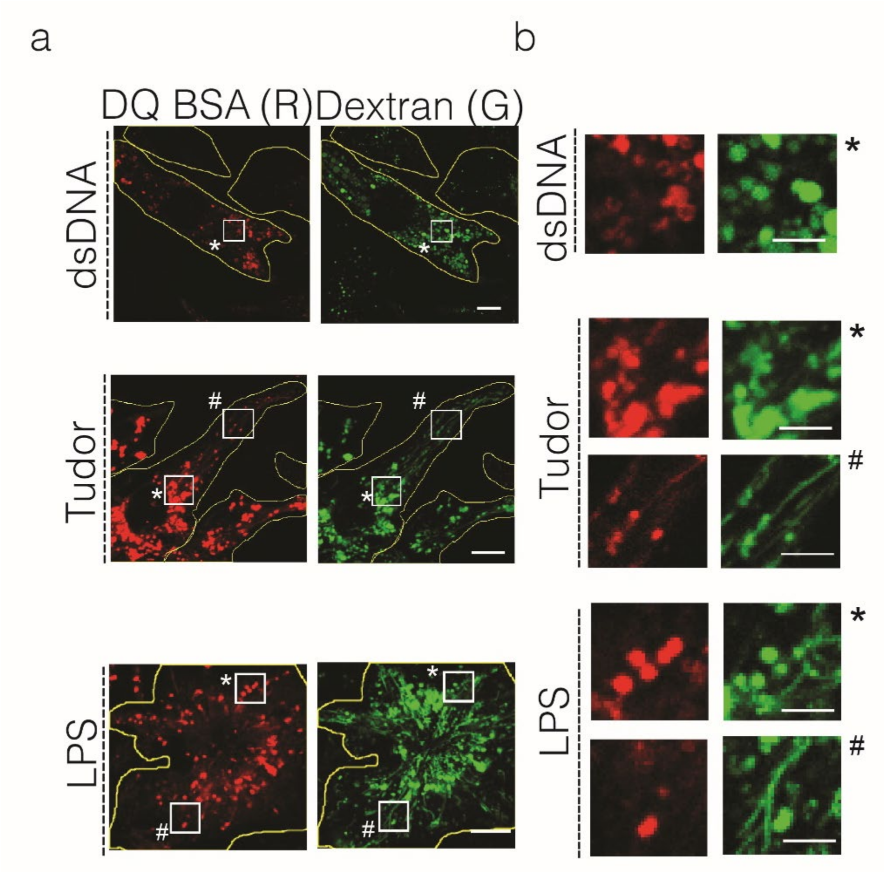


**Figure. S12: Differential hydrolysis in vesicular and tubular lysosomes of RAW 264.7 cells.** (a) Representative confocal images of Alexa 488 dextran-labeled lysosomes (G) in RAW 264.7 cells treated with dsDNA, *Tudor* or LPS followed by 10 µg/mL of DQ™ BSA Red (R). Scale bar: 10 µm. (b) Insets show magnified regions indicated * represents VLs and # represents TLs. Scale bar: 10 µm and inset scale bar: 4 µm.

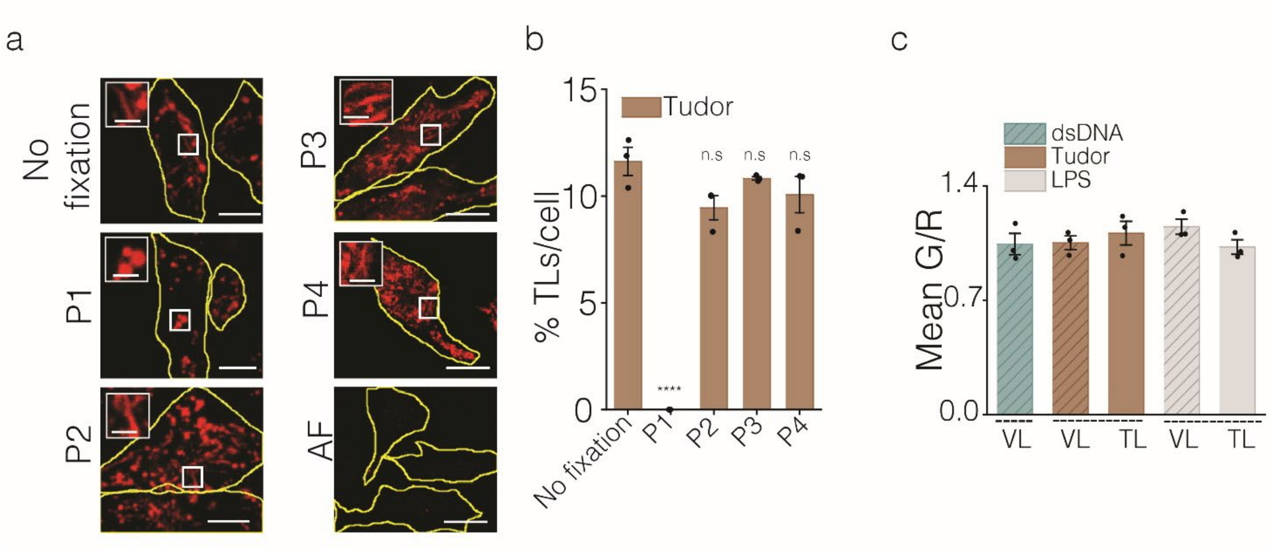


| Fixation methods used |
|---|
| P1: 3% Glyoxal |
| P2: 1% Glyoxal |
| P3: 0.5% PFA+0.45% GA |
| P4: 2% PFA+0.2 GA |

**Figure. S13: Fixation protocol for tubular lysosomes**. (a) Representative confocal images of TMR dextran labeled lysosomes in RAW 264.7 cells either untreated (no fixation and AF) or fixed using various mentioned fixatives (P1-P4). Insets show magnified regions indicated. Describe bottom table, either use a new panel # or describe as upper and bottom. (b) Quantification comparing the %TLs/cell between cells treated with various fixative compositions and un-fixed cells (n = 20 cells). ****P< 0.0005 (one-way ANOVA with Tukey *post hoc* test), n.s: non-significant. (c) Mean G/R plot showing fluorescence intensity from immunofluorescence of Cathepsin B (G) and LAMP1 (R) in presence of dsDNA, *Tudor* and LPS (n=15 cells) in RAW 264.7. Error bar represents s.e.m from three independent experiments. AF: Autofluorescence. Scale bar: 10 µM, with inset scale bar: 4 µm.

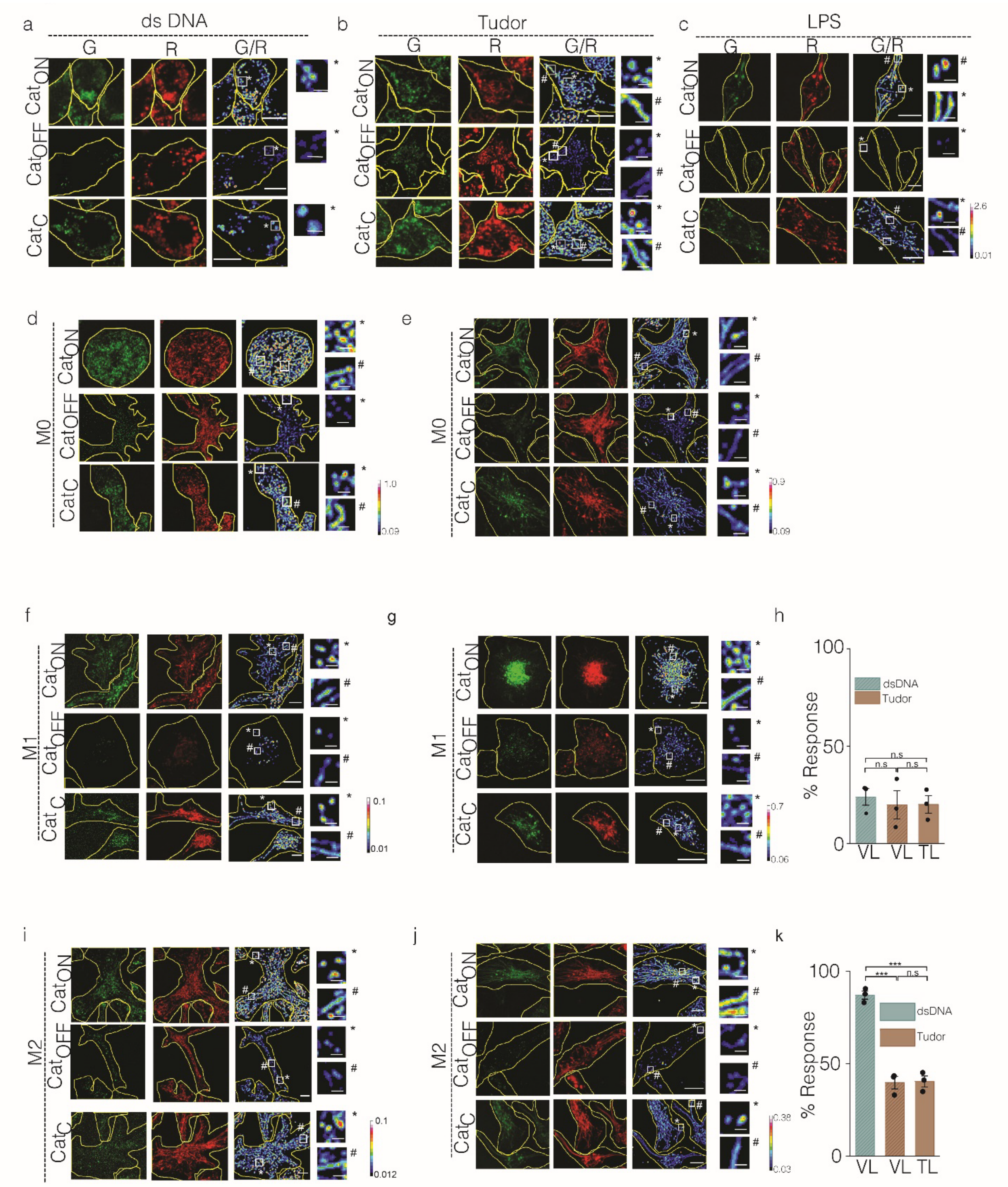


**Figure. S14: Cathepsin C activity in RAW 264.7 and BMDMs.** Representative confocal images of lysosomes in (a) dsDNA; (b) *Tudor* or (c) LPS treated RAW 264.7 cells labeled with CTC sensors ($Cat_{ON}$ $Cat_{C}$ and $Cat_{OFF}$) with or without E64. CTC activity measurement in dsDNA or *Tudor* treated BMDM (d, e) for M0; (f, g) for M1 and (i, j) for M2 macrophages. Inset shown in white box with * representing VLs and # represent TLs. (h, k) Quantification of % response of CTC in VLs and TLs upon treatment with dsDNA and *Tudor* in M1 and M2 BMDM respectively. All data obtained from three independent experiments with error representing s.e.m (n = 50 cells, m= 500 endosomes). ***P< 0.0005 (one-way ANOVA with Tukey *post hoc* test), n.s: non-significant. Scale bar: 10 µm. Inset scale bar: 4 µm.

**Supplementary Note 5: Cathepsin C (CTC) enzyme activity in VLs and TLs.** DQ™ BSA degradation assay revealed the overall enzyme activity within tubular lysosomes is lower as compared to vesicular lysosomes (Fig 3a-c). Previous literature shows that in autophagy, stimulated tubules lacked cathepsins and acid phosphatase(*31*). Yet immunofluorescence of cathepsin B showed equal levels of staining in VLs and TLs of *Tudor* and LPS stimulated RAW 264.7 (Fig S14c). We thus choose to study the enzymatic activity of CTC, one of the abundant lysosomal cysteine cathepsins. We used previously described DNA based CTC sensor ($Cat_C$) in this study(*6*) which consists of 2 modules namely, (i) sensing module made of azido Rhodamine 110 which is caged by a CTC cleavage motif, Gly-Phe dipeptides and a (ii) ratiometric module comprising of Alexa 647N (denoted as R) which is insensitive to any perturbations during this process (Fig 3e). Lysosomal CTC cleaves the N-terminus of Gly-Phe dipeptide and renders Rhodamine 110 free which allows it to fluoresce (denoted as G). DNA based Cathepsin C ON probe ($Cat_{ON}$) which consist of azido-Rhodamine 110 denoted as G, and Alexa 647N as R (Fig 3e). $Cat_{ON}$ sensor provides the measure of the maximum cleavage based fluorescence signal and hence provide the maximum G/R ratio. $Cat_C$ in presence of E64 (pan cathepsin inhibitor) shows the lowest or basal cleavage of $Cat_C$ sensor and therefore provides minimum G/R ratio. % Response of CTC activity within the VLs and TLs upon different treatments were calculated as described in Methods section with single lysosomal resolution.

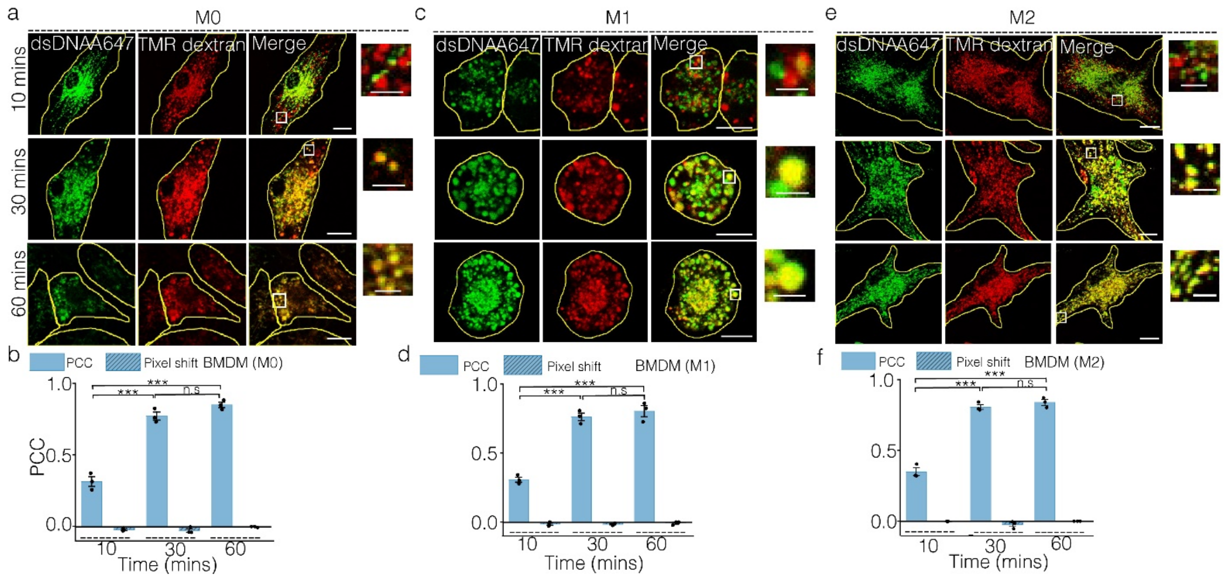


**Figure. S15**: **Time dependent colocalization of dsDNA with lysosomes in BMDM**. Representative confocal images of TMR dextran labeled lysosomes colocalized with dsDNA-A647 at different chase times of 10 mins; 30 mins and 60 mins in M0 (a); M1 (c) or M2 (e) BMDMs. Pearson's correlation coefficient (PCC) and pixel shift measured at each indicated chase time for M0 (b); M1 (d) and M2 (f). Images and data represented from three independent experiments and error bars represent s.e.m (n = 12 cells per experiment). ***P< 0.0005 (one-way ANOVA with Tukey *post hoc* test), n.s: non-significant. Scale bar: 10 µm. Inset scale bar: 4 µm.

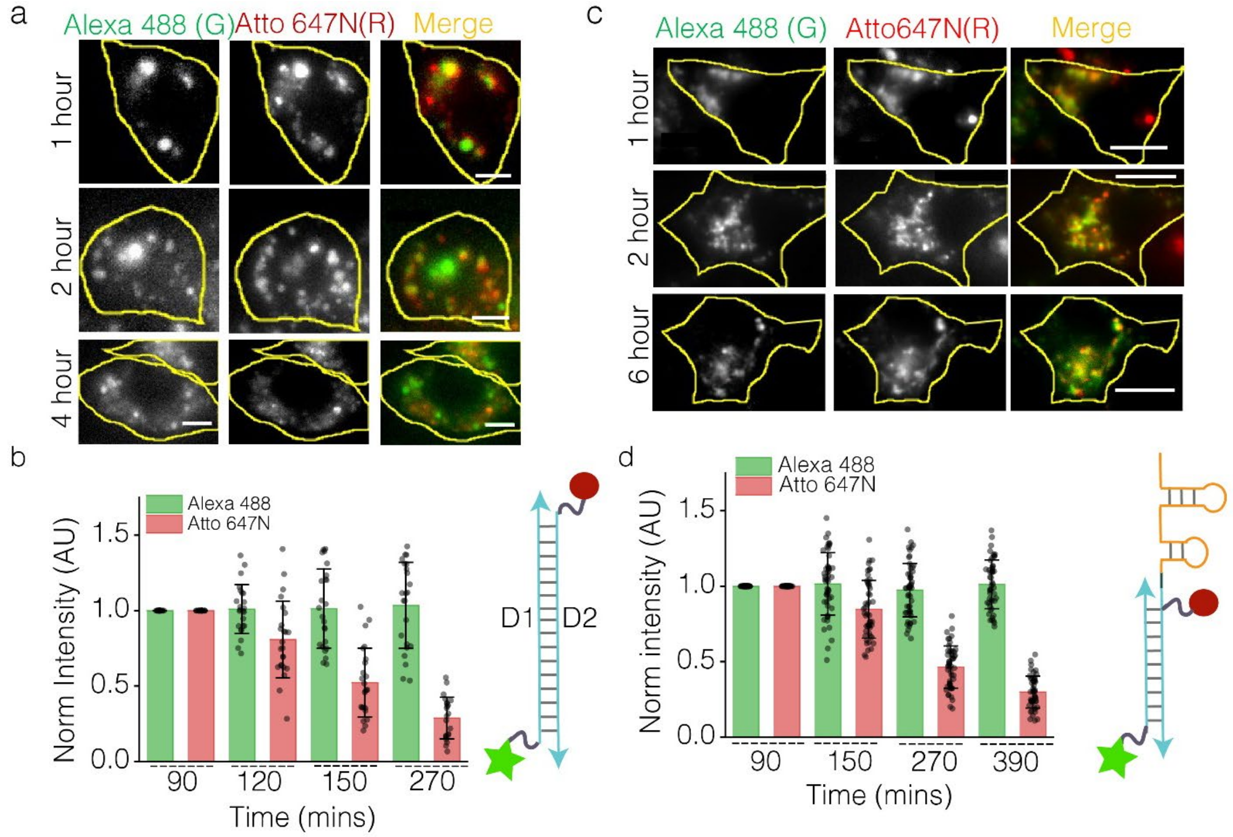


**Figure. S16: Stability of DNA nanodevices in VLs and TLs** (a) Representative images of RAW 264.7 cells showing dsDNA (Schematic shown below) uptake containing PEG-Alexa 488 (G) and Atto 647N (R) at different indicated chase times. Scale bar: 5 µm. (b) Quantification showing whole cell intensities of G and R as a function of chase times. (n= 20 cells) (c) Representative single plane wide field images of *Tudor* treated RAW 264.7 labeled with *Tudor* containing Atto 647N (R) and PEG-Alexa 488 (G) (Schematic shown on the right) in TLs at various chase times. (d) Quantification showing normalized intensity of TLs in R and G channels at different chase times. (n = 15 cells, m = 50 TLs per experiment). Error bars indicate standard deviation (SD). Scale bar: 10 µm. Data represented from three independent experiments.

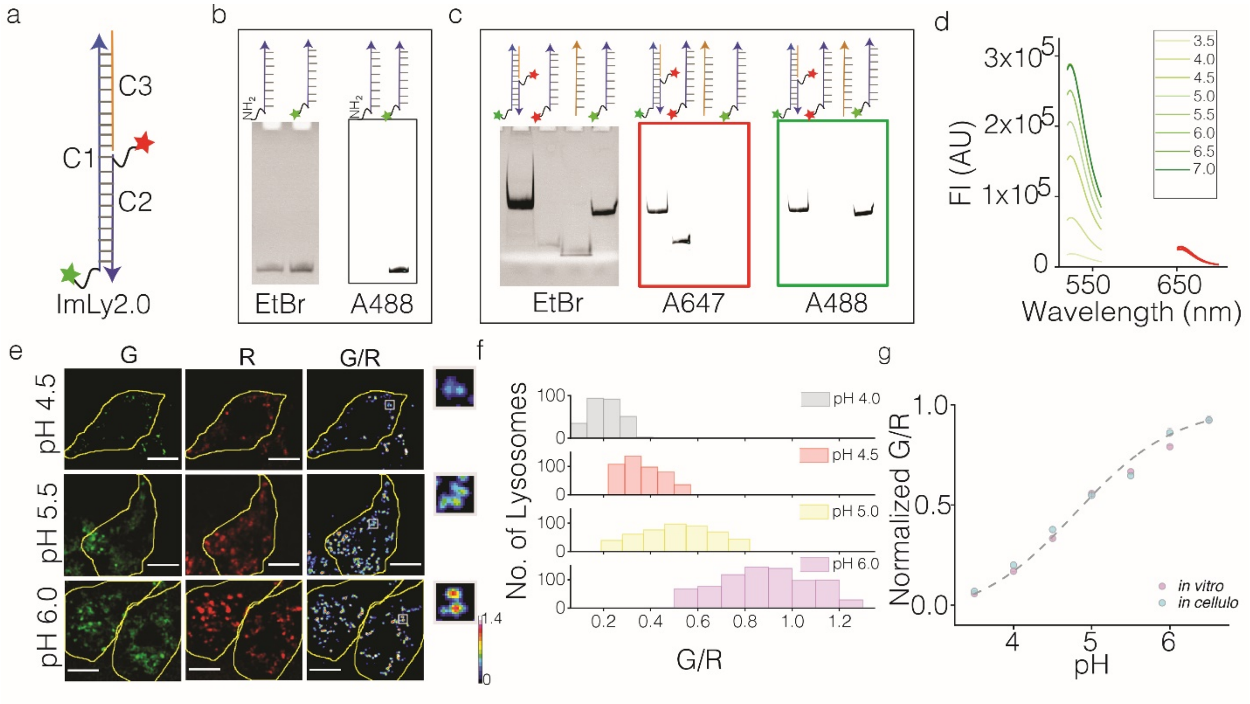


**Figure. S17: Characterization of *ImLy2.0*.** (a) Schematic of *ImLy 2.0* showing 5(6)-Carboxy-2',7'-dichlorofluorescein (DCF) on 58 mer (C1) oligo (green), Atto 647N (red) on the complimentary 28mer oligo (C2) and unlabeled 30mer oligo (C3). (b) Denaturating PAGE (15%) showing the conjugation of DCF to amine containing C1 in EtBr and Alexa 488 channels. (c) Gel mobility shift assay showing the formation of *ImLy 2.0* by 15% native PAGE imaged in EtBr, Alexa 647 and Alexa 488 channels. (d) Emission spectra of *ImLy 2.0* at pH ranging from 3.5 and 7.0. (e) Representative images of RAW 264.7 showing the uptake of *ImLy 2.0* and pixel wise pseudocoloured images of G/R clamped at indicated pH. Inset showing the zoomed in area shown in the white box. Scale bar: 10 µm. (f) Histogram of G/R ratios of lysosomes clamped at indicated pH (n= ≥ 90 cells, m = ≥ 500 lysosomes). (g) pH calibration (*in vitro*) for *ImLy 2.0* showing normalized G/R ratios versus indicated pH values. Error bars represents s.e.m from three independent experiments.

**Supplementary note 6: Calibration and lysosomal pH measurement using *ImLy 2.0:***

Previously reported DNA based pH sensor, *ImLy* senses pH reliably between pH 3.8 and pH 5.2(*17*). In order to probe pH between pH 4 and pH 6, we designed a new pH sensor *ImLy2.0* which is ideal for endo-lysosomal pH measurements. *ImLy2.0* comprises of 3 DNA strands, D1, 58 mer DNA strand consists of amino modification on 5' end which is used for conjugation with DCF (the pH sensing moiety). Conjugation of DCF to amino labeled DNA was confirmed by denaturing polyacrylamide gel electrophoresis. C2 is a 28 mer strand complementary to one half of C1 and is labeled with Atto 647N (pH and calcium insensitive dye) which acts as ratiometric moiety. C3 is a 30 mer strand which is complementary to the other half of C1. The formation of *ImLy2.0* was confirmed by mobility shift assay by native PAGE. *In vitro* calibration of *ImLy2.0* was performed by measuring the excitation and emission spectra in DCF (G) and Atto 647N (R) channels in universal buffer by varying pH ranging from pH 3.5 to pH 6.5. The ratio of G/R was plotted which was fitted to sigmoidal curve with Boltzmann fit. *In cellullo* calibration for *ImLy 2.0* was performed in RAW 264.7 using protocol mentioned in the methods section. The R/G ratios were taken at single lysosomal resolution from cells treated clamped at varying pH. The ratios were plotted similar to *in vitro* calibration curve. The *in cellulo* calibration curve recapitulated *in vitro* calibration curve suggesting optimal sensing properties *in cellulo*. *ImLy2.0* was used to measure pH of both vesicular and tubular lysosomes in RAW 264.7 using the pH calibration curve generated using pH clamping of cells at varying pH points.

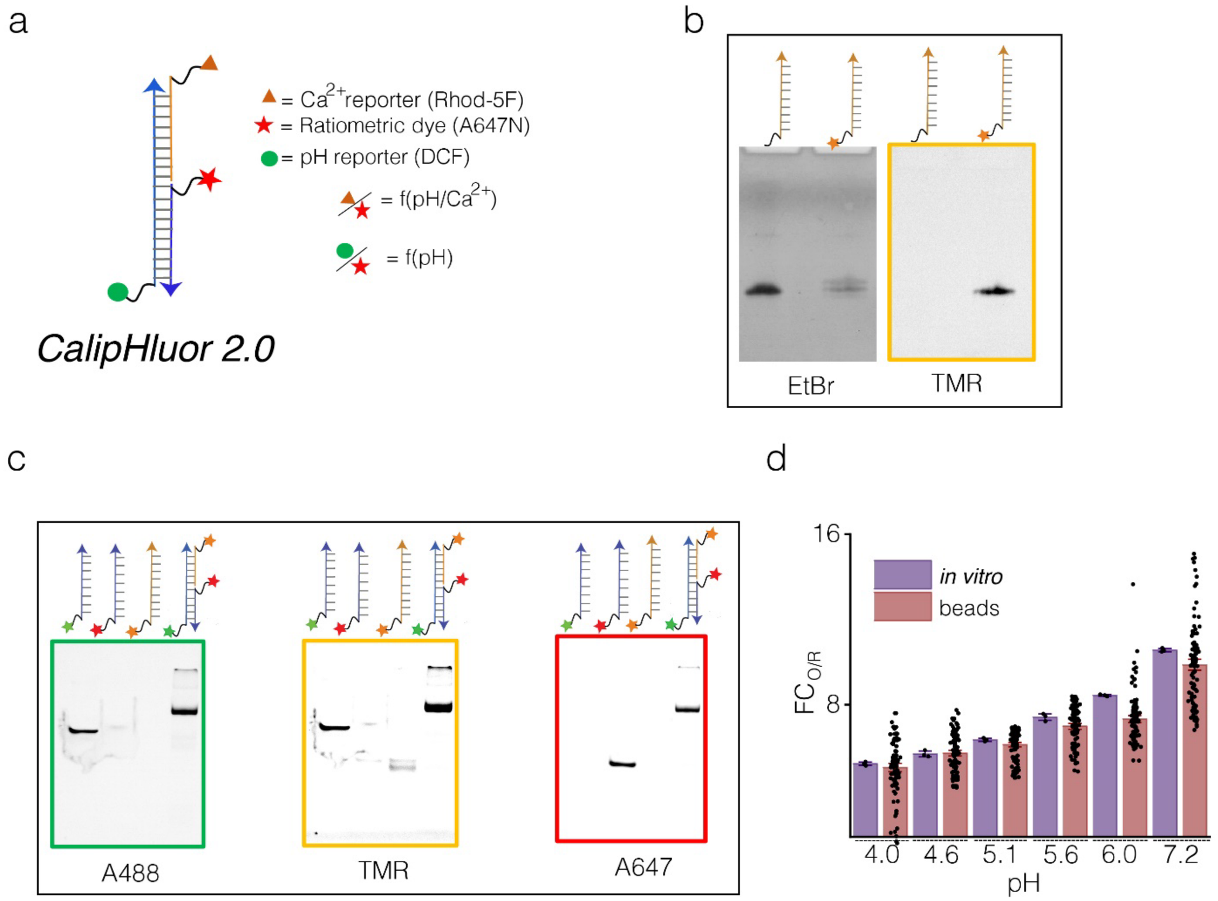


**Figure. S18: Characterization of *Caliphluor 2.0***: (a) Schematic of ratiometric fluorescent pH corrected $Ca^{2+}$ reporter, *CalipHluor 2.0*. It consists of $Ca^{2+}$ sensitive dye, Rhod-5F (orange triangle); pH sensing dye, DCF (green circle) and ratiometric dye, Atto 647N (red star). (b) Denaturing PAGE (15%) showing the conjugation of Rhod-5F to DBCO containing D3 oligo in EtBr and TMR channels. (c) Native PAGE (15%) showing gel mobility shift of *Caliphluor 2.0* in Alexa 488, TMR and Alexa 647 channels. (d) Comparison of *in vitro* (purple) and on beads (pink) fold change of O/R ($FC_{O/R}$) ratios of *Caliphluor 2.0* from pH 4.0 - 7.2 (n = 100 beads).

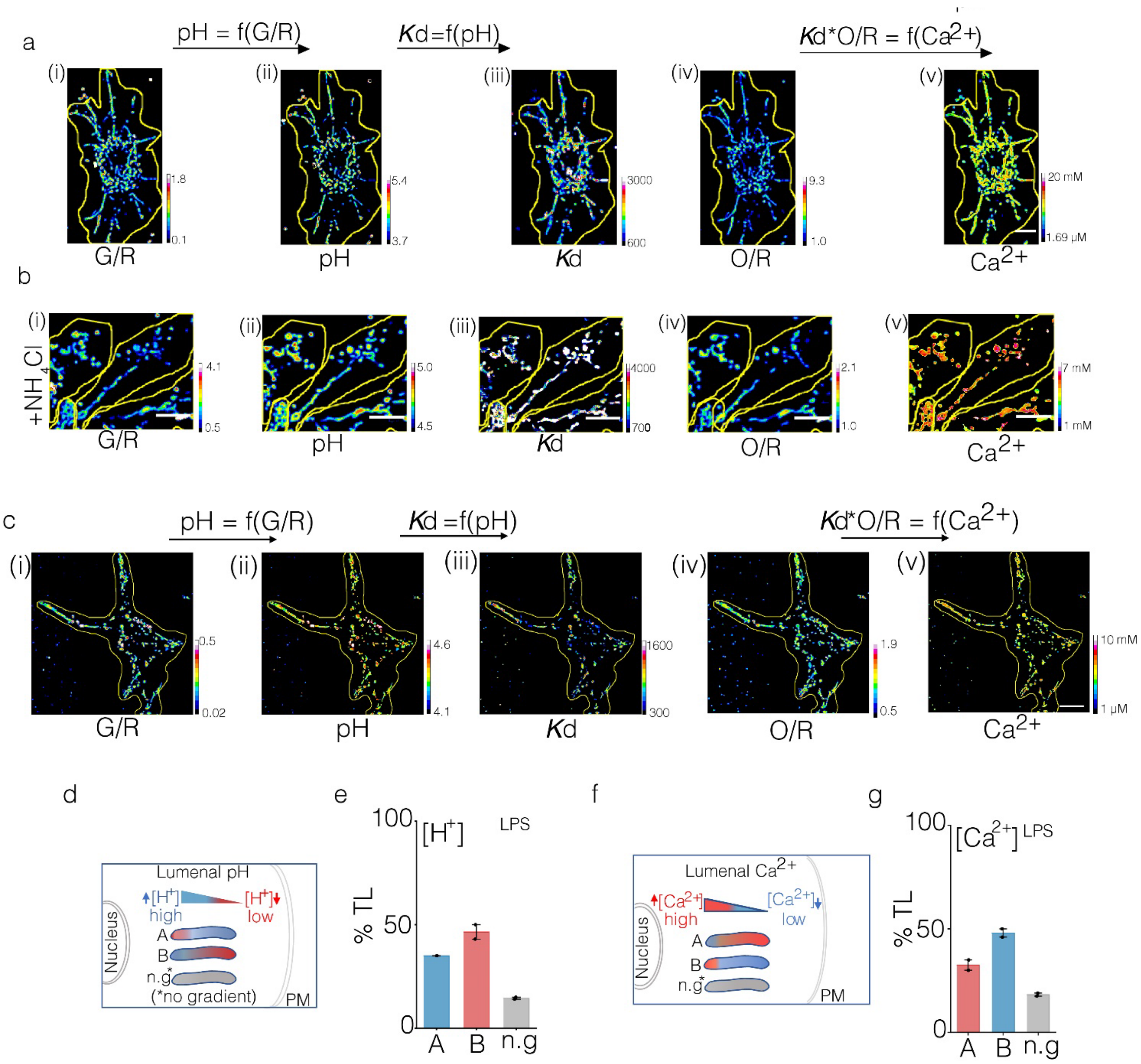


**Figure. S19: pH and Calcium images of RAW 264.7.** (a-b) Representative images from *Tudor* treated RAW 264.7 cells pulsed with *CalipHluor 2.0* followed by chase in complete media (a), media containing 10 mM $NH_4Cl$ (b). (c) Representative image of LPS treated RAW 264.7 cells pulsed with *CalipHluor 2.0* followed by chase in complete media. Pseudocolored maps of (i) G/R, (ii) pH, (iii), $K_d$, (iv) O/R (v) log [$Ca^{2+}$]. Scale bar: 10 µm. (d, f) Schematic of the lumenal pH and $Ca^{2+}$ gradients in tubular lysosomes oriented from nucleus to plasma membrane (PM). (e, g) % TLs showing pH and calcium gradient as per schematic in (d, f), (n=15 cells; m=50 TLs,). All error bars represent s.e.m from three independent experiments unless otherwise mentioned.

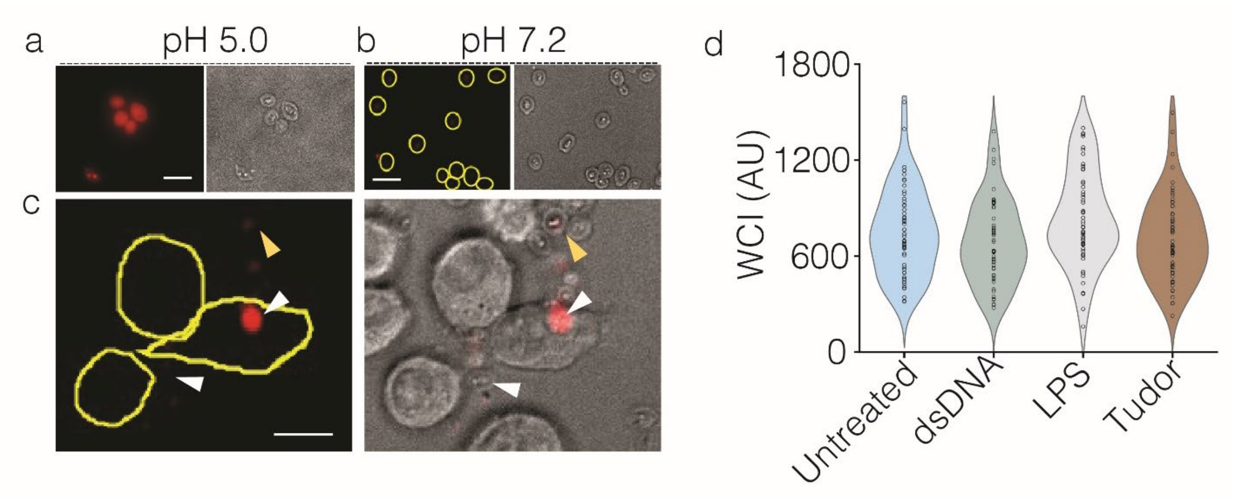


**Figure. S20: Phagocytosis of zymosan particles in RAW 264.7 cells.** Representative widefield images of pHrodo$^{TM}$ Red labeled zymosan imaged at (a) pH 5.0 and (b) pH 7.2. (c) Representative fluorescence (left) and brightfield images (right) of RAW 264.7 cells with phagocytosed pHrodo$^{TM}$ Red-zymosan. White arrowhead shows internalized zymosan. Yellow arrowhead shows the non-fluorescent zymosan outside the cells. Scale bar: 5µm. **(d)** Distribution of total cell intensity of fluid phase labeling of RAW 264.7 cells with Alexa 488 dextran. Cells were treated with either dsDNA, *Tudor* or LPS (n= 50 cells). Data represented from three independent experiments with similar results.

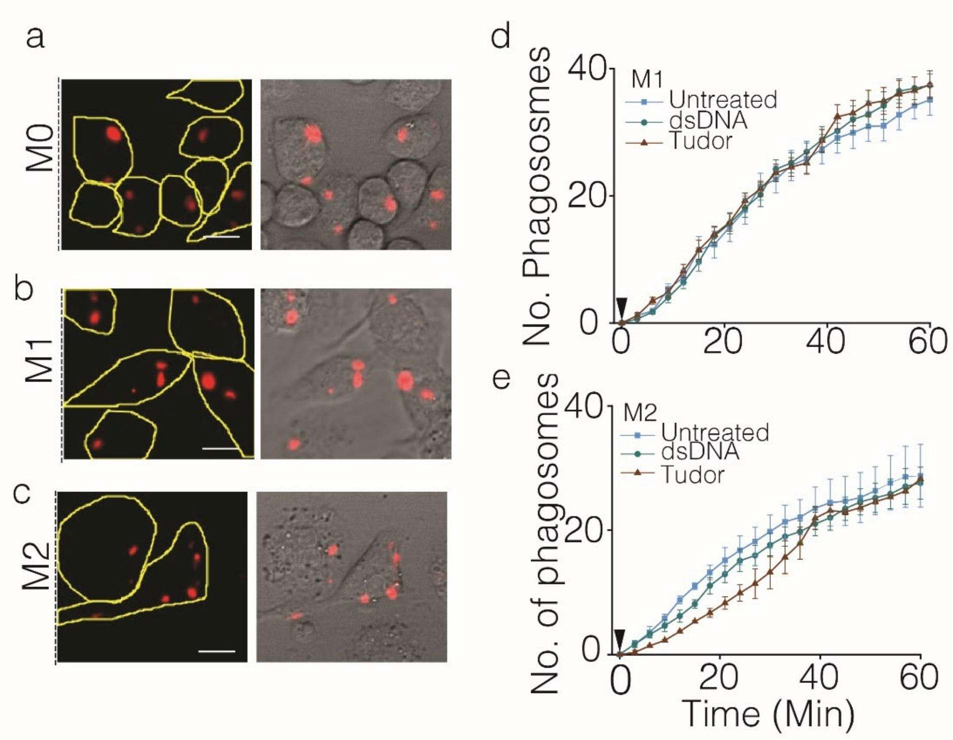

**Figure. S21: Phagocytic efficiency in M0, M1 and M2 macrophages of Pmacs.** (a-c) Representative widefield images of Pmacs showing pHrodo™ Red-zymosan uptake (fluorescence image, left and brightfield image, right). (d, e) Number of phagocytosed particles upon treatment with dsDNA or *Tudor* for 4 h. Arrowhead at t=0 min shows pHrodo™ Red-zymosan addition to cells (n = ~30 cells). Error bars represent s.e.m from three independent experiment. Scale bar: 10 µm.

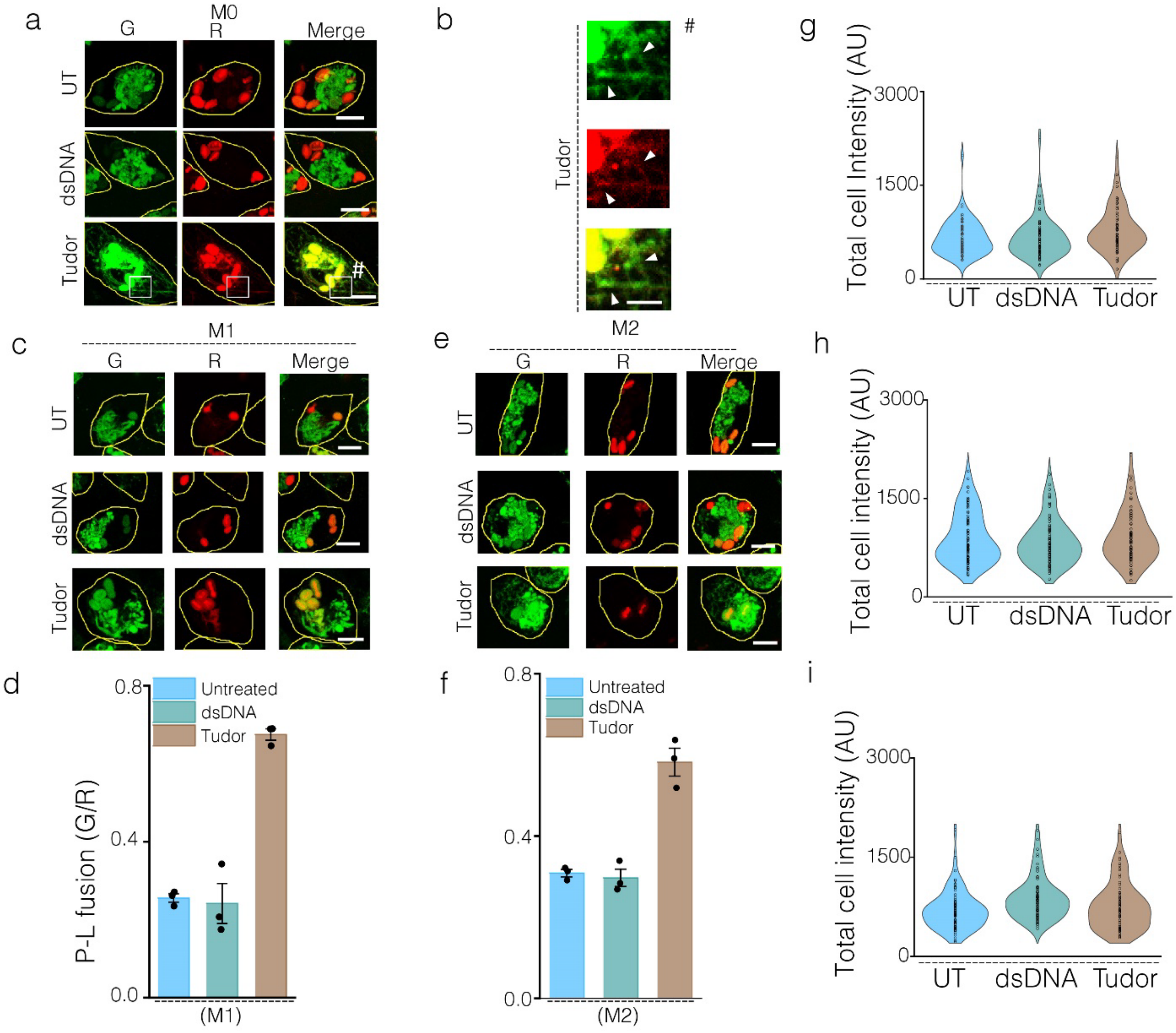


**Figure. S22: *Tudor* regulates phagosome lysosome fusion in M0, M1 and M2 of Pmac.** (a, c and e) Representative confocal images of lysosomes marked with Alexa 488 conjugated dextran (G) and pHrodo™ Red conjugated zymosan (R) in Pmacs upon treatment with culture media (untreated); dsDNA, *Tudor*. (b) Zoomed image of white box containing # in (a) with white arrow heads showing TL contacting phagosome. (d and f) Quantification of mean G/R showing the phagosome lysosome fusion (P-L fusion) in P macs (n = 50 cells, ~300 phagosomes). (g, h and i) Total cell intensity of Alexa 488 dextran containing lysosomes in Pmacs upon treatment with dsDNA and *Tudor* (n=50 cells). All errors showed here represent s.e.m from three independent experiments. Scale bars: 10 μm.

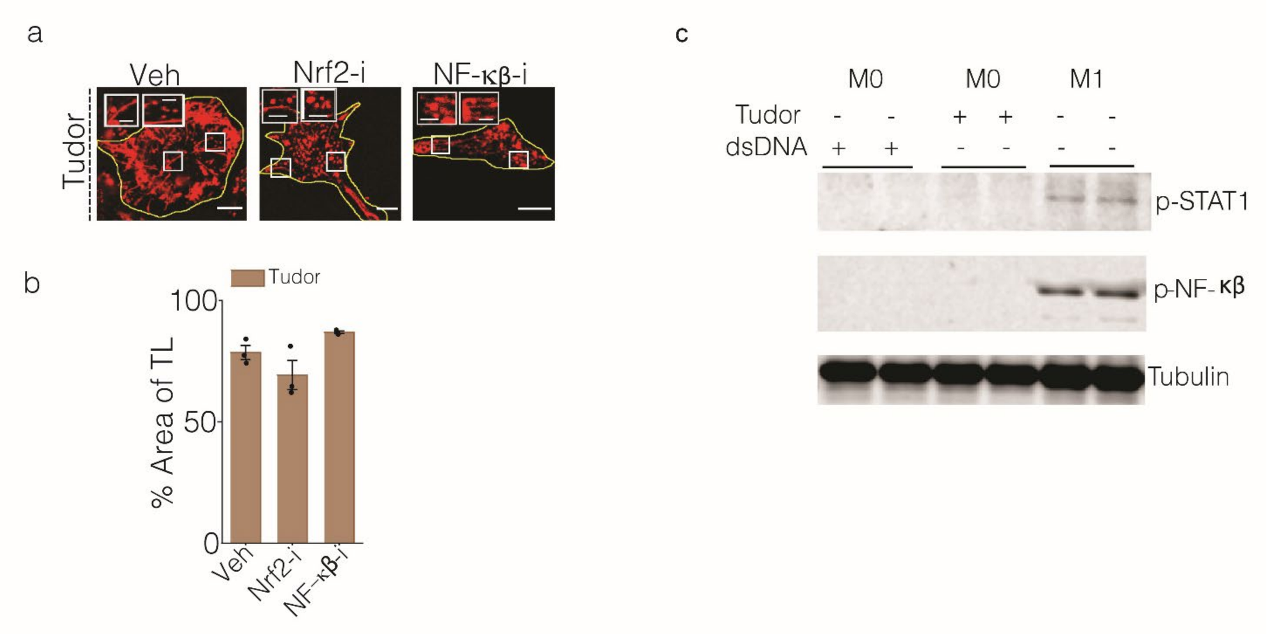


**Figure. S23: Nrf2 and NF-κβ activity in *Tudor* treated cells.** (a) Representative images of TMR dextran labeled lysosomes in *Tudor* treated RAW 264.7 cells in presence or absence of Nrf2 inhibitor (ML385) or NFkB inhibitor (JSH-23), (Veh=DMSO). (b) Quantification of % Area of TL in the presence or absence of inhibitors. Error bars represent s.e.m from three independent experiments (n =20 cells per experiment). Scale bar: 10 μm; Inset scale bars: 4 μm. (c) Representative western blots of p-STAT1, p-NFkB and tubulin in M0 BMDM treated with dsDNA or *Tudor* for 24 hours. M1 BMDM are shown as positive control for inflammatory pathway.

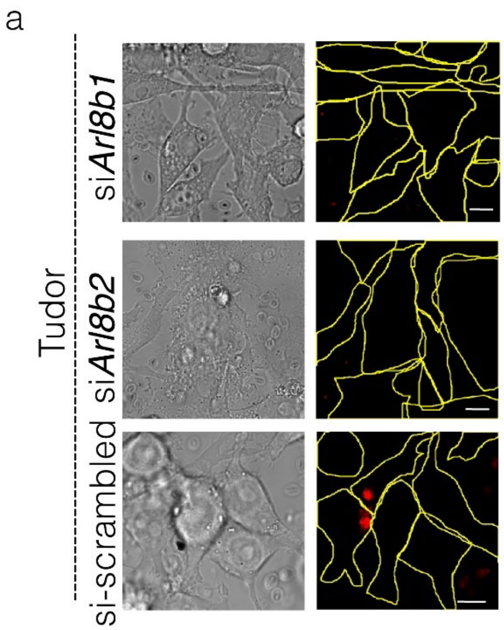


**Figure S24: Role of Arl8b in phagocytic uptake of zymosan in *Tudor* treated cells** (a) Representative images of RAW 264.7 cells showing its brightfield image (right) and pHrodo$^{TM}$ Red conjugated zymosan shown as red (left).

| Sequence name | DNA sequence information (5'-3') |
|---|---|
| SA43 | ACGTTACTCTTGCAACACAAACTTTAATAGCCTCTTATAGTT C |
| A1 | ACGTTACTCTTGCAACACAAACTTTAATAGCCTCTTATAGTT CTTCATCAACACTGCACACCAGACAGCA |
| A1-Atto647N | ACGTTACTCTTGCAACACAAACTTTAATAGCCTCTTATAGTT CTTCA/A647/TCAACACTGCACACCAGACAGCA |
| A2 | TGCTGTCTGGTGTGCAGTGTTGAT |
| A3 | ATCAACACTGCACACCAGACAGCA |
| A2-A647N | Alexa 647-TGCTGTCTGGTGTGCAGTGTTGAT |
| TRG2 | GGCTATAGCACATGGGTAAAACGACTTTGCT/Alexa 647/TGTCTGGTGTGCAGTGTTGAT |
| CpG | Atto 647-TGCTGTCTGGTGTGCAGTGTTGATTTtccatgacgttcctgacgtt |
| D1 | DBCO-ATCAACACTGCACACCAGACAGCAAGATCCTATATATA |
| D2 | Alexa 647-TATATATAGGATCTTGCTGTCTGGTGTGCAGTGTTGAT |

| | |
|---|---|
| C1 | Amino-ATAACACATAACACATAACAAAATATATATCCTAGAACGACAGACAAACAGTGAGTC |
| C2 | ATTO647-TATATTTGTTATGTGTTATGTGTTAT |
| C3 | DBCO-GACTCACTGTTTGTCTGTCGTTCTAGGATA |
| B1 | ATCAACACTGCACACCAGACAGCAAGATCCTATATATAACTAC |

**Table S1:** List of DNA nanodevices used in the study

| | |
|---|---|
| MUC1-dsDNA | 5-TRG2 aptamer linked to A2+A3 |
| CpG-dsDNA | CpG strand linked to A2+ A3 |
| dsDNA | A2 and A3 |
| ssDNA | B1 |

**Table S2:** Combinations of DNA used for specificity assays.

| **Protein** | **Inhibitor** | **Concentration/duration** | **Cat no.** | **Source** |
|---|---|---|---|---|
| TLR 4 | TAK-242 | 10 µM, 18 hours | 13871 | Cayman chemicals, USA |
| mTORC1 | Rapamycin | 100 nM, 1 hour | 13346 | Cayman chemicals, USA |
| mTORC2 | Torin 1 | 100 nM, 1 hour | 10997 | Cayman chemicals, USA |
| AMPK | Dorsomorphin | 20 µM, 30 mins | 21207 | Cayman chemicals, USA |
| PI3K | Zstk474 | 1 µM, 30 mins | 17381 | Cayman chemicals, USA |
| Akt | Akt inhibitor VIII | 5 µM, 30 mins | 14870 | Cayman chemicals, USA |
| Src1 | Dasatinib | 1 µM, 1 hour | 11498 | Cayman chemicals, USA |
| JAK | JAK inhibitor I | 1 µM, 48 hours | 15146 | Cayman chemicals, USA |
| Rac1 | NSC 23766 | 50 µM, 12 hours | 13196 | Cayman chemicals, USA |
| TAK1 | (5Z)-7-Oxo Zeaenol | 300 nM, 6 hours | 17459 | Cayman chemicals, USA |
| PLD1 | CAY10594 | 1 µM, 30 mins | 13207 | Cayman chemicals, USA |
| MMP9 | MMP-9 inhibitor I | 100 µM, 1 hour | 15942 | Cayman chemicals, USA |
| Integrin 1 | RGD peptide | 0.3 mg/mL, 4 hours | 14501 | Cayman chemicals, USA |
| Pan Cathepsin | E64 | | 10007963 | Cayman chemicals, USA |
| TLR3 | CuCPT-4a | 27 µM, 24 hours | 4884 | Tocris, USA |
| TLR5 | TH 1020 | 0.37 µM, 24 hours | 6191 | Tocris, USA |
| TLR2/6 | GIT-27 | 10ug/mL, 24 hours | 3270 | Tocris |

| | | | | |
|---|---|---|---|---|
| Myd88 | Myd88 Inhibitor peptide | 100 µM, 24 hours | NBP-2 29328 | Novus Biologicals, USA |
| LKB1 | LKB1-i | 380 nM, 24 hours | A3556 | APExBio |
| IRS1 | NT-157 | 1 µM, 72 hours | S8228 | Selleckchem |
| Sirtuin1 | EX527 | 1 µM, 24 hours | 100099798 | Cayman chemicals, USA |
| TLR1/2-i | CuCPT-22 | 8 µM, 24 hours | 4884 | Tocris, USA |
| Nrf2 | ML385 | 5 µM, 72 hours | 21114 | Cayman chemicals, USA |
| NFkB | JSH-23 | 300 µM, 1 hour | 481408 | Sigma Aldrich, USA |

**Table S3:** List of inhibitors used in the study.

| **Reagents** | **Catalog number** | **Source** |
|---|---|---|
| TMR conjugated 10kDa dextran | D1816 | Thermo Fisher Scientific |
| FITC conjugated to 10 kDa dextran | FD10S | Thermo Fisher Scientific |
| Amino dextran 10kDa | D3330 | Thermo Scientific |
| Zymosan A from Cerevisiae | Z4250 | Sigma |
| LPS | 2630 | Sigma |
| Bafilomycin | B1793 | Cayman Chemicals |
| Nigericin | 11437 | Cayman Chemicals |
| Ionomycin | I3909 | Cayman Chemicals |
| Monensin Sodium | 22373-78-0 | Cayman Chemicals |
| 40% Glyoxal solution | 128465 | Sigma |
| DQ™ BSA Red | D12051 | Invitrogen |
| ER Tracker™ Green | E34251 | Life technologies |
| LysoTracker™ Deep Red | L12492 | Thermo Scientific |
| Mito Tracker™ Green | M7514 | Thermo Scientific |
| SensoLyte 520 MMP-9 assay Kit | AS-71155 | AnaSpec Inc |
| 5(6)-Carboxy-2′,7′-dichlorofluorescein (DCF) | M1239 | Abcam |
| DMEM | 12400024 | Life technologies |
| Opti-MEM™ | 11058021 | Life technologies |
| FBS | 26140079 | Gibco |

**Table S4:** List of Reagents used in the study.

| **Antibodies** | **Catalog number** | **Source** |
|---|---|---|
| Ku 70 (1:100) | NB100-1915 | Novus Biologicals |

| | | |
|---|---|---|
| Cathepsin B (1:100) | CST 31718 | Cell Signaling technology |
| Lamp 1 (1:400) | ab24170 | Abcam |
| Pan Cadherin (1:500) | ab16505 | Abcam |
| PIP3(1:100) | Z-p345B | Echelon Biosciences |

**Table S5:** List of antibodies used in the study.

**Supplementary Video 1:** Timelapse images showing pH and Calcium gradient within the tubular lysosomes. Lysosomes in RAW 264.7 tubulated by *Tudor* and labeled with *CalipHluor 2.0* with change in pH in left and calcium on (right). Scale bar: 10 μM.

Bibliography

1. V. Prakash *et al.*, Quantitative mapping of endosomal DNA processing by single molecule counting. *Angew. Chem. Int. Ed. Engl.* **58**, 3073–3076 (2019).

2. S. Modi *et al.*, A DNA nanomachine that maps spatial and temporal pH changes inside living cells. *Nat. Nanotechnol.* **4**, 325–330 (2009).

3. A. Saminathan *et al.*, A DNA-based voltmeter for organelles. *Nat. Nanotechnol.* (2020), doi:10.1038/s41565-020-00784-1.

4. M. Kratz *et al.*, Metabolic dysfunction drives a mechanistically distinct proinflammatory phenotype in adipose tissue macrophages. *Cell Metab.* **20**, 614–625 (2014).

5. C. A. Reardon *et al.*, Obesity and Insulin Resistance Promote Atherosclerosis through an IFNγ-Regulated Macrophage Protein Network. *Cell Rep.* **23**, 3021–3030 (2018).

6. K. Dan, A. T. Veetil, K. Chakraborty, Y. Krishnan, DNA nanodevices map enzymatic activity in organelles. *Nat. Nanotechnol.* **14**, 252–259 (2019).

7. K. Leung, K. Chakraborty, A. Saminathan, Y. Krishnan, A DNA nanomachine chemically resolves lysosomes in live cells. *Nat. Nanotechnol.* **14**, 176–183 (2019).

8. J. Schindelin *et al.*, Fiji: an open-source platform for biological-image analysis. *Nat. Methods*. **9**, 676–682 (2012).

9. N. Narayanaswamy *et al.*, A pH-correctable, DNA-based fluorescent reporter for organellar calcium. *Nat. Methods*. **16**, 95–102 (2019).

10. D. N. Kalkofen, P. de Figueiredo, W. J. Brown, Methods for analyzing the role of phospholipase $A_2$ enzymes in endosome membrane tubule formation. *Methods Cell Biol.* **130**, 157–180 (2015).

11. Y. Sato *et al.*, Three-dimensional multi-scale line filter for segmentation and visualization of curvilinear structures in medical images. *Med Image Anal*. **2**, 143–168 (1998).

12. A. Saric *et al.*, mTOR controls lysosome tubulation and antigen presentation in macrophages and dendritic cells. *Mol. Biol. Cell*. **27**, 321–333 (2016).

13. K. N. Richter *et al.*, Glyoxal as an alternative fixative to formaldehyde in immunostaining and super-resolution microscopy. *EMBO J.* **37**, 139–159 (2018).

14. V. E. B. Hipolito *et al.*, Lysosome expansion by selective translation of lysosomal transcripts during phagocyte activation. *BioRxiv* (2018), doi:10.1101/260257.

15. M. Adamczyk, J. R. Fishpaugh, K. J. Heuser, Preparation of succinimidyl and pentafluorophenyl active esters of 5- and 6-carboxyfluorescein. *Bioconjug. Chem.* **8**, 253–255 (1997).

16. D. Moore, *Curr. Protoc. Immunol.*, in press, doi:10.1002/0471142735.im1001s8.

17. K. Chakraborty, K. Leung, Y. Krishnan, High lumenal chloride in the lysosome is critical for lysosome function. *Elife*. **6**, e28862 (2017).

18. A. T. Veetil, M. S. Jani, Y. Krishnan, Chemical control over membrane-initiated steroid signaling with a DNA nanocapsule. *Proc. Natl. Acad. Sci. USA*. **115**, 9432–9437 (2018).

19. J. Lahann, Ed., *Click chemistry for biotechnology and materials science* (John Wiley & Sons, Ltd, Chichester, UK, 2009).

20. S. Monferran, J. Paupert, S. Dauvillier, B. Salles, C. Muller, The membrane form of the DNA repair protein Ku interacts at the cell surface with metalloproteinase 9. *EMBO J.* **23**, 3758–3768 (2004).

21. Y. G. Y. Chan, M. M. Cardwell, T. M. Hermanas, T. Uchiyama, J. J. Martinez, Rickettsial outer-membrane protein B (rOmpB) mediates bacterial invasion through Ku70 in an actin, c-Cbl, clathrin and caveolin 2-dependent manner. *Cell Microbiol.* **11**, 629–644 (2009).

22. S. Monferran, C. Muller, L. Mourey, P. Frit, B. Salles, The Membrane-associated form of the DNA repair protein Ku is involved in cell adhesion to fibronectin. *J. Mol. Biol.* **337**, 503–511 (2004).

23. B. S. Prabhakar, G. P. Allaway, J. Srinivasappa, A. L. Notkins, Cell surface expression of the 70-kD component of Ku, a DNA-binding nuclear autoantigen. *J. Clin. Invest.* **86**, 1301–1305 (1990).

24. S. Aptekar *et al.*, Selective Targeting to Glioma with Nucleic Acid Aptamers. *PLoS One*. **10**, e0134957 (2015).

25. M. S. Jani, J. Zou, A. T. Veetil, Y. Krishnan, A DNA-based fluorescent probe maps NOS3 activity with subcellular spatial resolution. *Nat. Chem. Biol.* **16**, 660–666 (2020).

26. C. S. M. Ferreira, M. C. Cheung, S. Missailidis, S. Bisland, J. Gariépy, Phototoxic aptamers selectively enter and kill epithelial cancer cells. *Nucleic Acids Res.* **37**, 866–876 (2009).

27. A. P. West, A. A. Koblansky, S. Ghosh, Recognition and signaling by toll-like receptors. *Annu. Rev. Cell Dev. Biol.* **22**, 409–437 (2006).

28. A. T. Veetil *et al.*, DNA-based fluorescent probes of NOS2 activity in live brains. *Proc. Natl. Acad. Sci. USA*. **117**, 14694–14702 (2020).

29. D. Khatter, A. Sindhwani, M. Sharma, Arf-like GTPase Arl8: Moving from the periphery to the center of lysosomal biology. *Cell Logist*. **5**, e1086501 (2015).

30. N. A. Kaniuk *et al.*, Salmonella exploits Arl8B-directed kinesin activity to promote endosome tubulation and cell-to-cell transfer. *Cell Microbiol.* **13**, 1812–1823 (2011).

31. L. Yu *et al.*, Termination of autophagy and reformation of lysosomes regulated by mTOR. *Nature*. **465**, 942–946 (2010).
